# TXNDC15 modulated quality control at the endoplasmic reticulum shapes ciliogenesis

**DOI:** 10.64898/2026.04.01.715963

**Authors:** Vy N. Nguyen, Lena A. K. Bögeholz, Kit R. Page, Jichen Zhang, Alexa Ma, Melanie Ernst, Ting-Yu Wang, Natalie Chen, Adarsh Mayank, Maxine L. Wang, James Wohlschlegel, Tsui-Fen Chou, Alina Guna, Rebecca M. Voorhees

**Author notes:** These authors contributed equally to this work (Vy N. Nguyen, Lena A. K. Bögeholz). Correspondence (RMV) and (LAKB).

## Abstract

At the endoplasmic reticulum (ER), membrane protein quality control is tightly regulated to ensure excess subunits are recognized and degraded to protect cellular homeostasis. Using genome wide CRISPR screens, we identified a factor of unknown function, thioredoxin domain containing protein 15 (TXNDC15), and showed that it regulates membrane protein stability by tuning the activity of the E3-ubiquitin ligase MARCHF6. TXNDC15 modulates MARCHF6 in two opposing ways: first, it enhances the binding, ubiquitination, and degradation of membrane protein subunits with soluble cytosolic domains; and second, it prevents the inappropriate recruitment and ubiquitination of subunits with globular lumenal domains. Patient mutations to TXNDC15 that cause the ciliopathy Meckel-Gruber syndrome, disrupted its binding to MARCHF6, allowing degradation of critical ciliary proteins as they transit through the ER leading to defects in ciliogenesis. The regulatory function of TXNDC15 therefore exemplifies how protein quality control maintains the integrity of the proteome to prevent disease.

---

Nearly five thousand integral membrane proteins are encoded by the human genome, the majority of which must be inserted, modified, and assembled at the ER. This family includes proteins that will ultimately reside in diverse compartments of the cell, including ER-resident factors and those secreted to the Golgi, plasma membrane, and specialized compartments like the primary cilia. Defects in membrane protein biogenesis can lead to diseases ranging from neurodegenerative disorders to a suite of ciliopathies, many with incompletely understood molecular etiologies. To prevent cytotoxic effects, ER-associated protein degradation (ERAD) is responsible for the recognition, extraction and degradation of proteins that fail at any step of this biogenesis process.^1–3^ However, given the enormous biophysical and topological diversity in the human membrane proteome, the known ERAD factors alone cannot be responsible for surveillance of the thousands of nascent proteins that must transit through the ER.

One class of proteins that pose unique challenges for quality control are multisubunit membrane protein complexes.^4,5^ Complex assembly is highly regulated because unassembled subunits expose thermodynamically unfavorable intersubunit interfaces to the crowded cellular environment, which are prone to aggregation and non-specific interactions.^6,7^ In a healthy cell, a small number of orphan subunits, which are synthesized in excess or remain unassembled, must be identified and degraded.^8–10^ This problem is exacerbated in many diseases, including cancers and trisomies, where abnormal gene copy number results in unusually high levels of excess subunits.^11,12^

The cell therefore must have evolved mechanisms to tune protein homeostasis: subunits must be given sufficient time for assembly while excess subunits must be efficiently degraded.^13^ In mammals, ∼40% of membrane proteins must assemble into higher order oligomeric complexes, which include hundreds of ion channels and receptors, as well as members of large structures such as the nuclear pore complex and cilia transition zone.^14^ Despite the physiologic importance of the final assembled complexes, and the critical role subunit triage plays in cellular homeostasis, the factors responsible for the surveillance and triage of membrane protein subunits are not known.

## RESULTS

### Systematic mapping of factors that regulate orphan membrane protein subunit stability

To study the regulation of subunit assembly in human cells, we chose two distinct model systems that exemplify many of the common features of membrane protein complexes: GET1 and TMEM231. First, GET1 is a member of the essential, ubiquitously expressed, ER-resident membrane protein complex, GET1/2 (Figure 1A).^15–17^ The stability of GET1 depends on the presence of its binding partner, GET2: when unassembled, orphan GET1 is efficiently degraded in a proteasome dependent manner (Figure S1A).^18,19^ Second, GET1 contains an intramembrane interface composed of polar and charged residues that must be positioned within the hydrophobic core of the bilayer.^20,21^ These residues are likely required for assembly and function, as is commonly observed in many membrane protein subunits including ion channels and transporters.^22,23^ Finally, unlike more complex systems, GET1 contains only three TMs, and a structured cytosolic coiled coil domain, facilitating molecular analysis of its quality control.

**Figure 1.**
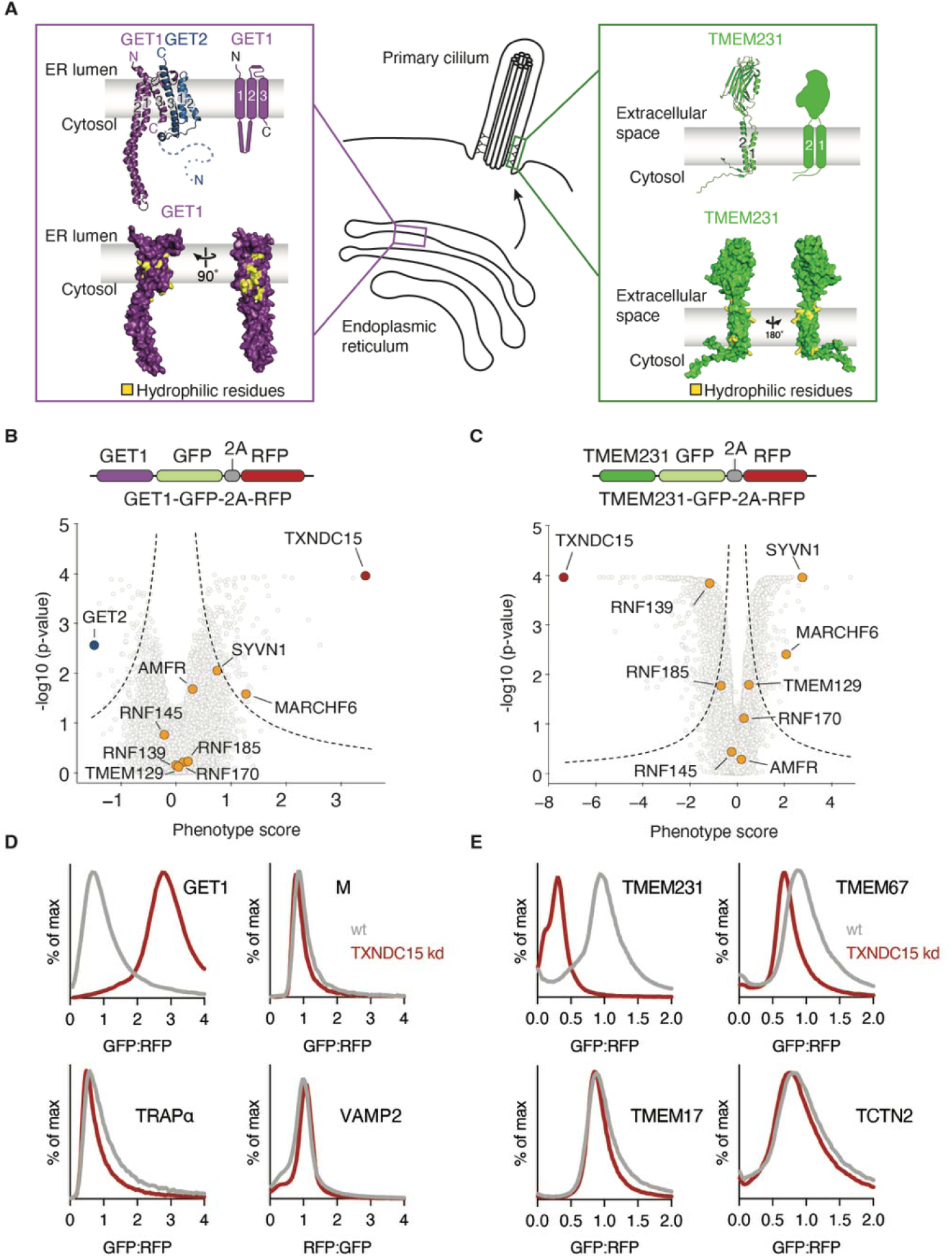
Systematic analysis of factors required for regulation of orphan GET1 and TMEM231. (A) Characterization of GET1 and TMEM231. (Left) AlphaFold3 predicted model of GET1 (purple) and GET2 (blue) aligned to the GET1/2 structure (PDB: 6SO5)^20^ compared to a topology diagram of GET1. Below, a surface representation of the GET1 model is shown with intramembrane hydrophilic residues highlighted in yellow that cluster at the intersubunit interface. (Middle) Schematic of the ER, where the GET1/2 complex resides and where TMEM231 is synthesized, and a primary cilium, the final localization of TMEM231 within the MKS complex located in the ciliary transition zone. (Right) Schematic of the AlphaFold3 predicted model of TMEM231 and its topology diagram. TMEM231 is predicted to contain two moderately hydrophobic TMs with intramembrane hydrophilic residues shown in yellow and a globular extracellular domain. (B) Genome wide CRISPRi screen for regulation of orphan GET1. Human K562 cells stably expressing GET1-GFP along with a translation normalization marker (RFP) from a single open reading frame were generated (Figure S1B). The volcano plot summarizes the phenotype for the three strongest single guide RNAs (sgRNAs) versus Mann-Whitney p-values calculated from two replicates of the genome wide CRISPRi screen. Individual genes are displayed in gray and specific factors highlighted and labeled, including select ER-resident E3 ubiquitin ligases (yellow). (C) Summary of genome wide CRISPRi screening results for factors that influence the stability of TMEM231 as described in (B). TXNDC15 (red) and select ER-resident E3 ubiquitin ligases (yellow) are highlighted as indicated. (D) Stability of the GET1 reporter described in (B) was assessed in wildtype (wt) and TXNDC15 depleted (kd) HEK293T cells. GET1-GFP expression relative to a normalization control was determined by flow cytometry and displayed as a histogram. To test whether depletion of TXNDC15 led to generalized dysregulation of the ER, we also determined the stability of a panel of membrane protein substrates with diverse topologies that rely on distinct targeting and insertion pathways into the ER including: a multipass membrane protein (the viral M protein from SARS-CoV-2), a single spanning membrane protein subunit (TRAPα), and a tail anchored protein (VAMP2). All data are normalized to the stability of each substrate in wildtype cells. (E) As in (D), but for TMEM231 and a panel of ciliary membrane proteins of the MKS complex. See also Figure S1, S2 and Table S1, S2.

Second, we chose TMEM231, a protein that must transit through the ER, but is ultimately localized to the primary cilium at the plasma membrane (Figure 1A).^24^ Primary cilia, unlike their motile counterparts, are sensory organelles found on most cell types. Their principal role is to function in both transducing environmental stimuli and regulating cellular signaling through the presentation of many essential receptors on their surface (e.g. sonic hedgehog).^25,26^ Consequently, a range of congenital ‘ciliopathies’ result from improper cilia function, and all share related developmental defects to the brain, kidneys, and limbs.^27^ TMEM231 is a member of the Meckel Syndrome (MKS) Complex, which contains at least eight subunits, and functions at the ciliary transition zone to regulate the diffusion of proteins between the bulk plasma and ciliary membrane.^28^ Genetic defects to TMEM231 are associated with several ciliopathies, including Meckel-Gruber syndrome and the more moderate Joubert syndrome.^29,30^ Though the structure of the MKS complex is still unknown, and the precise binding partners of TMEM231 are not defined, TMEM231 contains polar regions within its two membrane-spanning TMs that could form part of a subunit interface. Understanding how TMEM231 expression and stability is regulated could provide critical insight into the molecular basis of its role in ciliopathies.

To systematically identify factors involved in the regulation of these orphan subunits, we generated two human K562 cell lines that stably expressed green fluorescent protein (GFP)-tagged GET1 or TMEM231 along with a translation normalization marker (red fluorescent protein, RFP) and the CRISPR inhibition (CRISPRi) machinery (Figure S1B).^31–33^ We reasoned that depletion of factors that affect the stability or degradation of GET1 or TMEM231 would result in either a decrease or increase in GET1-GFP:RFP fluorescence, respectively. Following transduction with a genome-wide CRISPRi sgRNA library^34^, cells displaying altered GFP:RFP fluorescence were isolated by FACS and deep sequenced.

### Depletion of TXNDC15 leads to accumulation of GET1 and destabilization of TMEM231

In analysis of the GET1 screen, as expected, depletion of its binding partner, GET2, resulted in decreased levels of the fluorescent GET1 reporter (Figures 1B, S1D, Table S1). Loss of known ERAD factors such as the ubiquitin-activating enzyme 1 UBA1 or the AAA+ ATPase VCP and its cofactor FAF2 stabilized GET1 (Figure S1C).^35–37^ Notably however, the greatest stabilization of GET1 was caused by depletion of a protein of unknown function, thioredoxin domain containing 15 (TXNDC15).

TXNDC15 is a single spanning membrane protein predicted to contain an N-terminal signal sequence followed by a thioredoxin domain and C-terminal TM (Figure S1E). While its molecular function is undefined, it was initially identified in patients where mutations to TXNDC15 lead to one of the most severe ciliopathy disorders, Meckel-Gruber syndrome.^38–40^ Notably, mutations to TMEM231 are also associated with the same disorder.^28^ Depletion of TXNDC15 in cells has also been shown to effect cilia morphology and function.^41,42^

Despite this association with the primary cilia, we confirmed using an arrayed approach that loss of TXNDC15 stabilized our ER-resident GET1 reporter (Figure 1D). Because this ratiometric reporter acutely expresses both GET1 and a normalization marker from a single open reading frame, we concluded that the effect of TXNDC15 must occur post-translationally on newly synthesized GET1, as would be expected for a quality control factor. We further validated that depletion of TXNDC15 (Figure S1F) specifically affected the stability of unassembled GET1, but not the assembled GET1/2 complex (Figure S1H) and confirmed that the observed phenotype is independent of the fluorescent labeling strategy utilized (Figure S1G). Finally, a panel of other membrane protein substrates with diverse topologies was unaffected by TXNDC15 depletion, excluding generalized dysregulation of the ER (Figure 1D).

To further probe the function of TXNDC15, we next performed a genetic modifier screen using our GET1 reporter cell line (Figures S2A-B, Table S1).^43^ We observed that depletion of several known ERAD factors— including DERL2 (Der1 in yeast), SYVN1 (Hrd1 in yeast), SEL1L (Hrd3 in yeast)—had an enhanced phenotype in the TXNDC15 knockdown compared to the control screen.^44–48^ The increased effects of these canonical ERAD factors upon depletion of TXNDC15 are indicative of function in parallel pathways, consistent with a potential role for TXNDC15 in membrane protein quality control.

However, upon analysis of the TMEM231 CRISPRi screen, we found that depletion of TXNDC15 appeared to have the opposite effect on TMEM231: loss of TXNDC15 destabilized TMEM231 (Figure 1C, Table S2). We validated this phenotype in an arrayed format (Figures 1E, S2C), and found that some, but not all members of the MKS complex were also destabilized by depletion of TXNDC15. Together these results suggested that TMEM231 and GET1 represented two specific and distinct classes of TXNDC15 substrates. We thereby set out to try to determine the apparently contradictory roles of TXNDC15 in regulating the stability of these two membrane protein subunits.

### TXNDC15 regulation of subunit stability depends on binding MARCHF6 in the ER

We reasoned that one potential explanation for the differing effects of TXNDC15 on the ER-resident GET1 versus the ciliary TMEM231 was that TXNDC15 has distinct functions in these two compartments. To study TXNDC15 localization, we first generated a human cell line in which it was fluorescently tagged at its endogenous locus (Figure S3A). We found that the majority of TXNDC15 is localized to the ER (Figure 2A), and we were unable to identify a population in the primary cilia, even when overexpressed (Figure S3C). This is consistent with an earlier study that reported that overexpressed TXNDC15 was localized to the ER where it is anchored by ERp44.^41,49^ We therefore concluded that TXNDC15 must be exerting its effect on GET1 and TMEM231 in the ER, excluding the possibility of a compartment-specific function.

**Figure 2.**
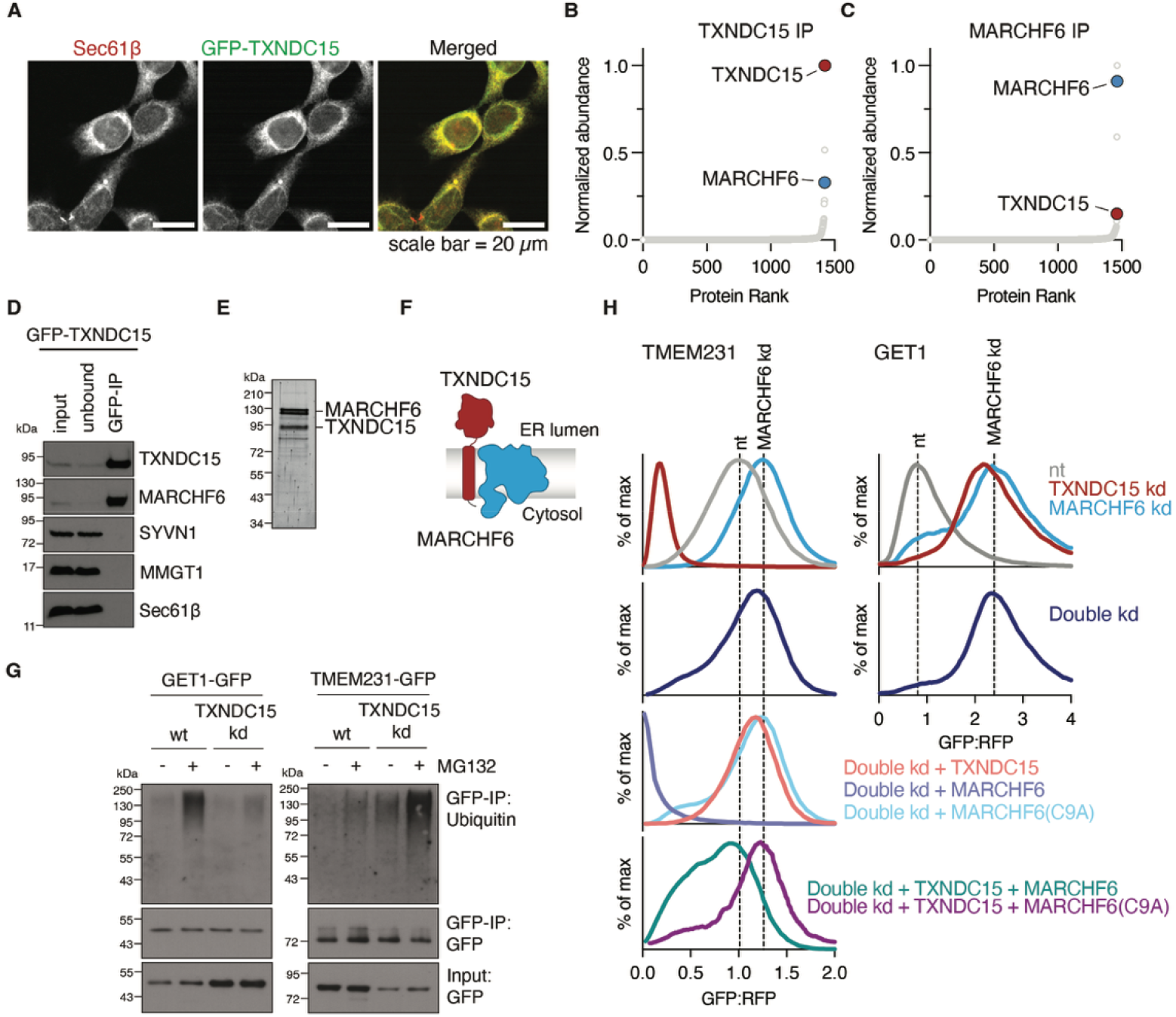
TXNDC15 is an ERAD factor that binds and functions with MARCHF6. (A) The localization of TXNDC15 was determined by imaging HEK293T cells in which an N-terminal GFP tag was inserted at the endogenous TXNDC15 locus (described in Figure S3A). Staining for SEC61β was used as an ER marker. Imaging of cilia was subsequently performed using RPE1 cells overexpressing TXNDC15-GFP (Figure S3C). (B) Analysis of the interactome of endogenously expressed TXNDC15 using mass spectrometry. GFP-TXNDC15 was solubilized and purified from the endogenously tagged HEK293T cell line under native conditions in the detergent GDN. Co-purifying proteins were ranked by average abundance from two replicates and plotted against their normalized abundance (See materials and methods for more details). Background controls for non-specific interactions are shown in Figure S3D. (C) As in (B) for exogenously expressed MARCHF6(C9A)-GFP isolated from Expi293 cells. MARCHF6(C9A) was used as it is the more stable, catalytically inactive variant of MARCHF6. Background controls are shown in Figure S3E. (D) To study the interaction partners of TXNDC15, we generated HEK293T cells in which TXNDC15 and MARCHF6 were tagged at their endogenous loci. Following immunoprecipitation of GFP-TXNDC15, samples were analyzed by western blot for co-purification of MARCHF6 and several other abundant ER-resident specificity controls. (E) Purification of a recombinant MARCHF6(C9A)-TXNDC15 complex visualized using Sypro Ruby. Expi293 cells expressing TXNDC15-ALFA and MARCHF6(C9A)-GFP were solubilized and immunoprecipitated using first the anti-ALFA-nanobody and then the anti-GFP-nanobody, sequentially. (F) Schematic summarizing our model that TXNDC15-MARCHF6 interact directly at ∼1:1 ratio. (G) Analysis of the ubiquitination of GET1-GFP and TMEM231-GFP upon TXNDC15 depletion. K562 cell lines stably expressing GET1 and TMEM231 were transduced with either a TXNDC15 sgRNA or a non-targeting (nt) control. After a 6 hr treatment with the proteasome inhibitor MG132 to allow for accumulation of ubiquitinated species, cells were lysed and GET1-GFP and TMEM231-GFP were immunoprecipitated. Elution samples were normalized by GFP levels as determined by fluorescence spectroscopy and ubiquitinated species were analyzed by western blotting. (H) Analysis of GET1 and TMEM231 stability using the ratiometric fluorescent reporter system described in Figure 1 following depletion with the indicated sgRNAs in K562 cells. (Left) For TMEM231, depletion of MARCHF6 and TXNDC15 had the opposite effects on subunit stability: loss of MARCHF6 stabilized TMEM231, while loss of TXNDC15 destabilized TMEM231. Simultaneous depletion of both TXNDC15 and MARCHF6 (double knockdown) phenocopied loss of MARCHF6 alone, leading to stabilization of TMEM231. This double knockdown could not be rescued by expression of TXNDC15 alone, a catalytically inactive variant of MARCHF6 (MARCHF6(C9A)), or TXNDC15 together with MARCHF6(C9A). Conversely, overexpression of MARCHF6 in this double knockdown background led to a strong destabilization of the TMEM231 reporter. Only the addition of TXNDC15 together with functional MARCHF6 can rescue the double knockdown phenotype. (Right) Depletion of either TXNDC15 or MARCHF6 individually stabilized GET1. Their simultaneous depletion did not cause an additive effect on GET1 stability, consistent with their function in the same degradation pathway. See also Figure S3 and Table S3.

To study its role in the ER, we next used our endogenously tagged cell line to immunoprecipitate TXNDC15 under native conditions to identify co-purifying factors using mass spectrometry (Figures 2B, S3D, Table S3). We found that one of the strongest interaction partners is the ER-resident E3-ubiquitin ligase, MARCHF6 (Doa10 in yeast), a component of the ERAD-C and M pathways. The interaction of MARCHF6 with TXNDC15 appeared to be highly specific, as other abundant ER-resident membrane proteins did not display similar enrichment (Figure 2D).^50–54^ Immunoprecipitation of exogenously expressed MARCHF6 from whole cells similarly resulted in a pronounced enrichment of TXNDC15 (Figures 2C, S3E, Table S3). Recombinant co-expression and purification suggested that TXNDC15-MARCHF6 could form a stable, direct, and nearly stoichiometric complex (Figure 2E, F). Depletion experiments suggested that TXNDC15 is an obligate binding partner of MARCHF6, as loss of MARCHF6 led to subsequent destabilization of TXNDC15 over time (Figure S3F, S3G).

The association with MARCHF6 suggested TXNDC15 could play a role in protein degradation, which was consistent with the observed stabilization of GET1 upon depletion of TXNDC15. However, the effect of TXNDC15 on TMEM231 was clearly more complicated, as its depletion destabilized TMEM231. To test whether TXNDC15 could also be involved in regulating the degradation of TMEM231, we analyzed the ubiquitination of GET1 and TMEM231 upon TXNDC15 depletion (Figure 2G). As expected, loss of TXNDC15 led to a decrease in ubiquitinated GET1, explaining GET1’s accumulation upon TXNDC15 depletion. In contrast, we observed an accumulation of ubiquitinated TMEM231 in the absence of TXNDC15, both in the presence and absence of proteasome inhibition, which would explain the apparent destabilizing effects of loss of TXNDC15 on TMEM231. We therefore concluded that TXNDC15 is exerting both its stabilization and degradation effects through modulation of the ubiquitin-proteasome pathway at the ER.

To test whether this role of TXNDC15 in quality control relied on MARCHF6, we performed a series of genetic interaction experiments using reporters for both GET1 and TMEM231. First, we found for GET1 that the effects of TXNDC15 and MARCHF6 depletion were epistatic, suggesting they functioned together in its degradation (Figure 2H). For TMEM231, we similarly observed that depletion of both MARCHF6 and TXNDC15 phenocopied MARCHF6 depletion alone. Rescue of this double knockdown with MARCHF6 destabilized TMEM231 independent of TXNDC15, while expression of TXNDC15 alone had no effect. Rescue of the MARCHF6 and TXNDC15 dual depletion required expression of both TXNDC15 and a catalytically active variant of MARCHF6 (Figure 2H). These results suggested that the ubiquitination activity of MARCHF6 was regulated by TXNDC15. Consistent with this model, expression of MARCHF6 alone in wildtype cells destabilized TMEM231, presumably stimulating its degradation (Figure S3H). Conversely, overexpression of TXNDC15 alone phenocopied MARCHF6 knockdown, resulting in stabilization of TMEM231 in a dose dependent manner (Figure S3H-I). Based on these experiments, we concluded that the occupancy of TXNDC15 at MARCHF6 dictates the stability of TMEM231. Therefore, given the observed genetic and biochemical interactions, we propose that TXNDC15’s role in both the stabilization and degradation of membrane subunits in the ER must depend on regulating the catalytic activity of MARCHF6.

### TXNDC15 requires its lumenal domain, but not thioredoxin reductase activity for its role in ERAD

We next sought to define what features of TXNDC15 were required to regulate nascent subunit stability in the ER. First, we used a series of mutational experiments to identify which domains of TXNDC15 were required for its activity (Figures 3A, S4A-B). Leveraging our reporter assay, we established that while anchoring of the TXNDC15 lumenal domain to the ER membrane is required for its effects on GET1 and TMEM231, the sequence of the TM itself is dispensable. We found that mutations to TXNDC15 that disrupted its binding to MARCHF6, e.g. by removing the lumenal domain from the immediate vicinity of the membrane through addition of a rigid linker, could not rescue effects of TXNDC15 knockdown on GET1 or TMEM231. The stability of the TXNDC15-MARCHF6 complex is therefore critical for its function. From these experiments, we concluded that the integrity and localization of the TXNDC15 lumenal domain is both necessary and sufficient for its role in both quality control and stabilization of membrane protein subunits in the ER.

**Figure 3.**
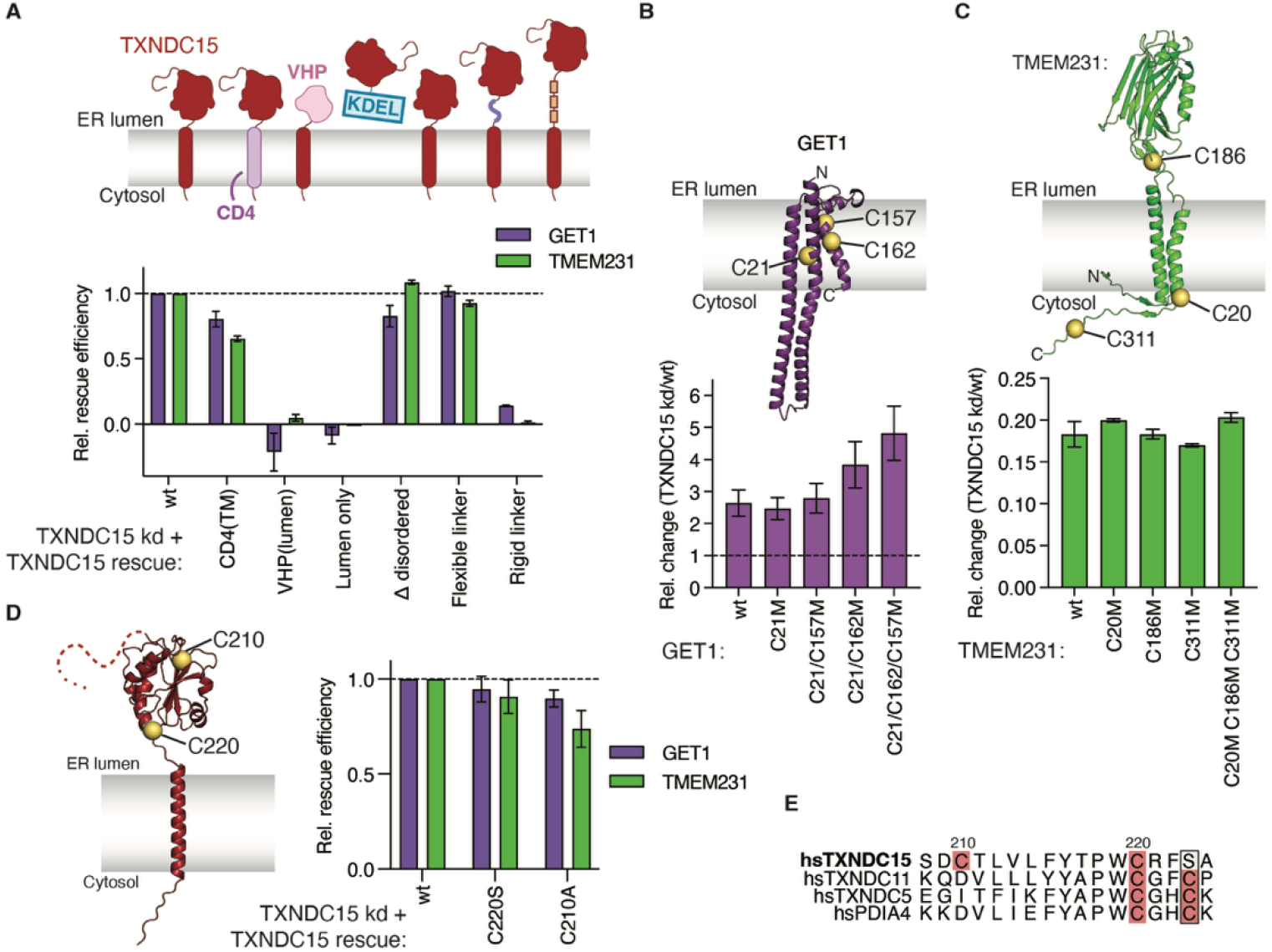
TXNDC15 does not require thioredoxin reductase activity for its role in regulating subunit stability in ERAD. (A) Rescue assay to identify domains essential for TXNDC15’s function in regulating GET1 and TMEM231 stability in HEK293T cells. The following constructs were transduced in TXNDC15 depleted cells: wildtype (wt) TXNDC15; the TXNDC15 lumenal domain fused to an unrelated TM from CD4 (CD4-TM); TXNDC15 with its lumenal domain replaced with the unrelated globular domain of bVHP (bVHP-lumen); the TXNDC15 lumenal domain alone fused to a KDEL ER-retention sequence (lumen-KDEL); TXNDC15 without its disordered region (Δ disordered); TXNDC15 with a flexible or rigid linker between the thioredoxin-like domain and TM (flexible/rigid linker). The dashed line at 1 indicates rescue to wildtype levels. Relative rescue efficiency was calculated based on the percent rescue relative to wildtype cells and normalized to rescue with wt TXNDC15 (described in methods). Assays were performed in triplicate and error bars indicate one standard deviation. Associated western blots are shown in Figure S4A. (B) To assess whether substrate cysteines were required for TXNDC15’s function, we generated the indicated GET1 cysteine mutants (yellow spheres). The relative change in stability of each reporter was calculated as: median GFP:RFP fluorescence in TXNDC15 depleted cells/median GFP:RFP in wt cells. Error bars represent the standard deviation calculated from 3 replicates. The dashed line at 1 is the relative change we would expect for substrates that are not dependent on TXNDC15. (C) Cysteine mutations in TMEM231 analyzed by fluorescence flow cytometry as in (B) with associated western blots shown in Figure S4C. (D) As in (A) for the indicated TXNDC15 cysteine mutants at residues 210 and 220 (shown as yellow spheres). Note that while TXNDC15 contains a cysteine at position 210, this is not within the putative thioredoxin reductase active site. (E) Multiple sequence alignment of the putative catalytic domain of human TXNDC15 with the indicated proteins in the disulfide isomerase (PDI) family (TXNDC11, TXNDC5, PDIA4). The positions of the two catalytic cysteines (residues 220 and 223) are highlighted in red. See also Figure S4.

Given the critical function of the TXNDC15 lumenal domain, which contains a thioredoxin reductase fold, we tested whether enzymatic activity was required for TXNDC15-mediated membrane subunit regulation. For example, another thioredoxin domain containing protein, TXNDC11, is an established ERAD factor that acts to reduce its substrates’ disulfide bonds to enable their degradation.^55^ Therefore, if TXNDC15 was functioning as a canonical thioredoxin reductase, we would expect that its function would depend on the presence of cysteine residues within its substrates. However, mutation of all three cysteines in either GET1 or TMEM231 did not decrease the effects of TXNDC15 depletion on their stability (Figures 3B-C). Further, sequence alignment of TXNDC15 suggested it is missing one of the required catalytic cysteines (Figure 3E), and mutation of its remaining active-site cysteine did not affect its quality control function (Figures 3D, S4C). These results suggested that the role of TXNDC15 in ERAD does not require covalent client trapping, though TXNDC15 may rely on this strategy for other purposes.^49^ We therefore concluded that TXNDC15 must have evolved a distinct role in recognition and/or degradation of its clients that relies on a non-enzymatic function of its lumenal domain.

### The molecular features that lead to substrate degradation vs stabilization by TXNDC15

To understand the potential role of TXNDC15 in modulating MARCHF6, we dissected which properties of GET1 and TMEM231 conferred dependence on TXNDC15. To do this, we exploited the observation that the stability of TMEM237, a ciliary membrane protein, was dependent on MARCHF6 but not TXNDC15 (Figure S5A). We reasoned that because TMEM237 was efficiently recognized by MARCHF6 alone, introducing defined domains of GET1 or TMEM231 into TMEM237 (our TXNDC15 independent substrate) could delineate which features require TXNDC15 for either degradation or stabilization, respectively. Importantly, all chimeras of GET1 or TMEM231 with TMEM237 remained MARCHF6 dependent, excluding the possibility that the observed phenotypes could reflect degradation by an alternative E3-ubiquitin ligase pathway (Figures S5A-B).

We found that successive fusions of TMEM237 with GET1 became most dependent on TXNDC15 for their degradation when they included both the cytosolic coiled coil and TMs 1-2 of GET1 (Figures 4A, S5A).

**Figure 4.**
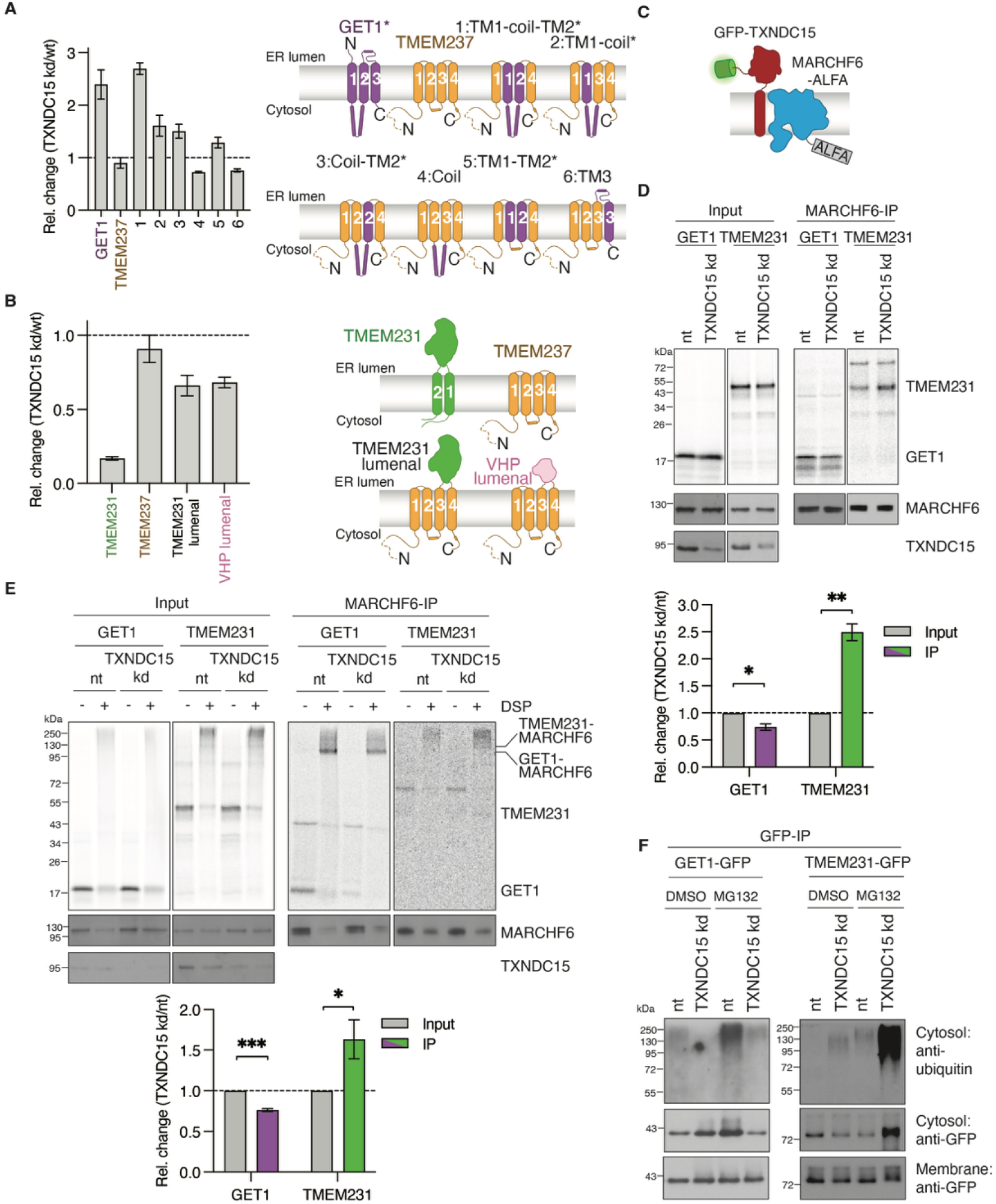
TXNDC15 regulates recruitment and ubiquitination of substrates to MARCHF6 based on their structural features. (A) To identify regions of GET1 that confer TXNDC15 dependence for degradation by MARCHF6, we generated a series of chimeras between GET1 (requires TXNDC15 for degradation) and TMEM237 (TXNDC15 independent). A ratiometric fluorescent reporter assay was used to analyze the stability of GET1, TMEM237, and the indicated chimeras in wildtype (wt) or TXNDC15 knockdown (TXNDC15 kd) HEK293T cells. Introduction of TMs 1-2 and the cytosolic coiled coil of GET1 into TMEM237 conferred TXNDC15 dependence for degradation by MARCHF6 and TXNDC15 dependent chimeras were labelled with *. Mean and standard deviation were calculated from 3 replicates. The dashed line at 1 is the relative change we would expect for chimeras that are not dependent on TXNDC15. All fusion constructs remained MARCHF6 dependent as shown in Figure S5A. (B) As in (A) but to identify features of TMEM231 that lead to its stabilization by TXNDC15 from MARCHF6 degradation. Chimeras of TMEM237 (TXNDC15 independent) and TMEM231 (stabilized by TXNDC15) were generated as indicated. Additionally, to test the effect of fusion with an unrelated globular domain on TMEM237’s degradation by MARCHF6, we also tested the effects of fusion with the soluble Villin Head Piece (VHP) domain. Error bars represent the standard deviation (n = 3). The dashed line at 1 is the relative change we would expect for chimeras that are not dependent on TXNDC15. Associated histograms are shown in Figure S5B. (C) Schematic of endogenously labeled GFP-TXNDC15 and MARCHF6-ALFA. (D) Reconstitution of the function of TXNDC15 *in vitro* by measuring nascent substrate association to MARCHF6. An ^35^S-methionine labeled TMEM231 or GET1 was translated in rabbit reticulocyte lysate (RRL) in the presence of human derived ER microsomes (hRMs) prepared from HEK293T cells containing endogenously tagged GFP-TXNDC15 and MARCHF6-ALFA. To acutely deplete TXNDC15, cells were treated with either nontargeting (nt) or TXNDC15 shRNA prior to purification of ER microsomes. MARCHF6-ALFA was then immunoprecipitated and eluted under native conditions using the anti-ALFA-nanobody (MARCHF6-IP). Co-purification of either TMEM231 or GET1 with MARCHF6 was analyzed by SDS-PAGE, autoradiography and western blot. The relative change in amounts of immunoprecipitated substrates between nt and TXNDC15 kd samples was normalized to the relative change in the input. Mean and standard deviation are displayed for 3 replicates and a one-sample t-test was calculated to determine if the mean significantly differed from 1 (p(GET1) = 0.016; p(TMEM231) = 0.004). The dashed line at 1 is the relative change we would expect for substrates whose binding is not affected by TXNDC15. (E) Using a similar *in vitro* platform as described in (D), we measured recruitment of GET1 or TMEM231 to MARCHF6 using chemical crosslinking to visualize binding that could not be captured by co-immunoprecipitation. After insertion of ^35^S-methionine labeled GET1 or TMEM231 into human ER microsomes, the isolated membrane fraction was incubated with the primary amine-specific crosslinker, DSP, or a DMSO control. The endogenously tagged MARCHF6-ALFA was enriched by ALFA-IP. Quantification was performed as in (D) for GET1 (n = 4; p = 0.0001) and TMEM231 (n = 3; p = 0.045). (F) Measurement of the ubiquitination and retrotranslocation of GET1-GFP and TMEM231-GFP. K562 cell lines stably expressing GET1 and TMEM231 were transduced with either a TXNDC15 sgRNA or a nt control. After a 6 hr treatment with the proteasome inhibitor MG132 to allow for accumulation of ubiquitinated species, cells were lysed and the soluble and membrane fractions were isolated by centrifugation. GET1 and TMEM231 were affinity purified from each fraction. Samples were normalized by GFP levels in the membrane fraction as determined by fluorescence spectroscopy and ubiquitinated species were analyzed by western blotting. Input samples are shown in Figure S5C and analysis of harvested cells using the ratiometric fluorescence reporter assay is displayed in Figure S5D. See also Figure S5.

Introduction of neither the coiled coil nor TMs 1-2 alone was sufficient to confer TXNDC15 dependence. These experiments indicated that it is the combination of these intramembrane and cytosolic features of GET1 that required TXNDC15 for MARCHF6-dependent degradation.

Chimeras with TMEM231 showed that introducing its globular lumenal domain into TMEM237 was sufficient to confer a protective effect of TXNDC15 (Figures 4B, S5B). Because the N-termini of these chimeras remained unchanged, TXNDC15 dependent regulation of MARCHF6 cannot depend on recognition of an N-terminal acetylation degron.^56^ Inserting an unrelated globular domain also led to a similar TXNDC15-dependent stabilization, suggesting it was the presence of a globular domain, and not necessarily its sequence or biophysical properties, that was important. We concluded that TXNDC15 must prevent the MARCHF6-dependent degradation of substrates that contain a globular lumenal domain, resulting in their stabilization.

### TXNDC15 is an ERAD factor that tunes the recruitment and ubiquitination of substrates by MARCHF6

Having identified the critical domains of TXNDC15 and its substrates, we finally addressed how TXNDC15 confers these opposing effects on TMEM231 and GET1. We hypothesized that if TXNDC15 was not playing an enzymatic role in subunit regulation, it could instead affect substrate recruitment to MARCHF6. To test this, we translated orphan GET1 and TMEM231 in a cell-free system in the presence of ER-microsomes purified from an endogenously tagged MARCHF6 cell line (Figure 4C, S3A,B). Prior to microsome purification, partial TXNDC15 depletion was achieved through acute treatment with shRNA. Following insertion of each nascent substrate into the membrane, the reactions were solubilized in detergent, and MARCHF6 was affinity purified to determine the efficiency of substrate recruitment. We found that orphan GET1 could be specifically enriched by immunoprecipitation of MARCHF6 in the presence of TXNDC15, validating that nascent substrates physically associated with the TXNDC15-MARCHF6 complex as would be expected for a bona fide quality control factor (Figure 4D). However, in the absence of TXNDC15, orphan GET1 was less efficiently immunoprecipitated by MARCHF6, suggesting that TXNDC15 is required to mediate its recruitment and interaction. Conversely, depletion of TXNDC15 led to an increase in immunoprecipitation of TMEM231, suggesting that for this class of substrates, TXNDC15 prevented its recruitment to MARCHF6.

Using a similar in vitro system, we used chemical crosslinking to stabilize the potentially transient interactions between substrates and MARCHF6 (Figure 4E). Consistent with our immunoprecipitation results, the depletion of TXNDC15 decreased crosslinking of GET1, but increased crosslinking of TMEM231 to MARCHF6. Together, these results provide a biochemical explanation for the apparently contradictory effects of TXNDC15 on the stability of GET1 and TMEM231: TXNDC15 enhances recruitment of GET1 to the E3-ubiquitin ligase, while preventing recruitment of TMEM231.

To test whether these changes in substrate recruitment in vitro resulted in functional consequences in cells, we next queried the effects of TXNDC15 depletion on ubiquitination and retrotranslocation of GET1 and TMEM231 (Figures 4F, S5C, S5D). Depletion of TXNDC15 resulted in a marked decrease in ubiquitination of both the membrane and cytosolic pools of GET1, consistent with TXNDC15 being required for the degradation of orphan GET1. The observed decrease in the ubiquitinated GET1 in the cytosol would further suggest that TXNDC15-mediated ubiquitination by MARCHF6 is required for its retrotranslocation from the ER. We also observed that the ubiquitination of TMEM231 was enhanced by loss of TXNDC15, consistent with its protective effect on TMEM231. The striking changes in ubiquitination may be the result of the cumulative effect of a multiple turnover reaction in cells, which may not be fully reproduced in vitro.

Together these data in cells and in vitro suggested that TXNDC15 modulates client recruitment to MARCHF6—both enhancing and inhibiting binding of distinct classes of substrates—which in turn alters their ubiquitination and stability.

### Mapping the suite of TXNDC15-dependent substrates

Having established a role of TXNDC15 in modulating recruitment of GET1 and TMEM231 to MARCHF6, we next sought to identify other TXNDC15-dependent substrates. To do this, we used two complementary approaches. First, we performed an unbiased proteomics experiment to identify endogenous proteins where depletion of TXNDC15 led to an increase (similar to GET1) or decrease (similar to TMEM231) in their steady-state levels (Figure 5A). Comparison across several replicates indicated that many membrane proteins are dependent on TXNDC15, underscoring its general role in regulating protein homeostasis in the ER. We validated a subset of these proteomics hits using our ratiometric fluorescent reporter assay, which specifically queries post-translational effects on the stability of newly synthesized proteins, suggesting these were bona fide TXNDC15-dependent substrates (Figure S6).

**Figure 5.**
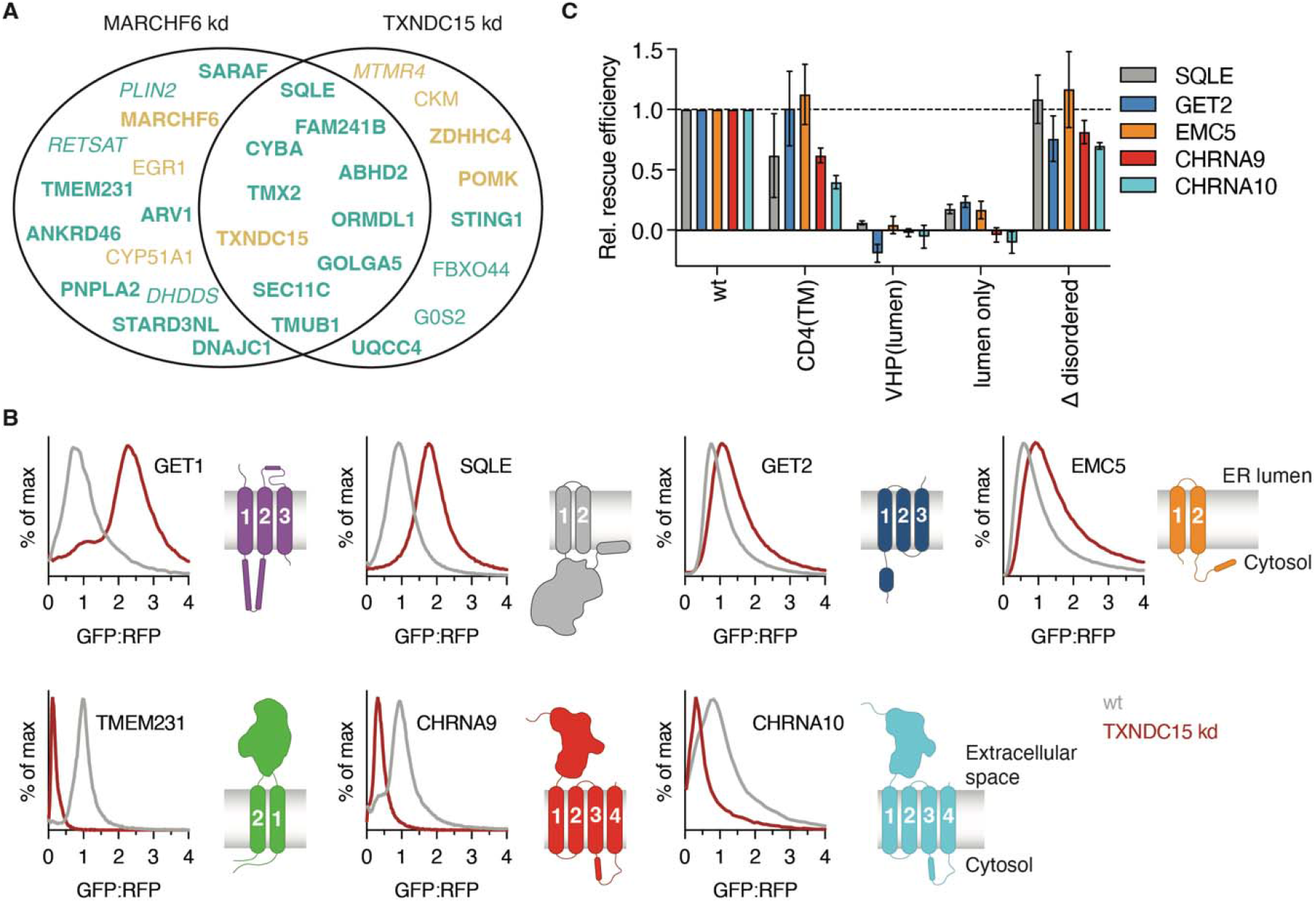
TXNDC15 plays a general role in regulating protein homeostasis in the ER. (A) Quantitative proteomics to identify potential endogenous TXNDC15 substrates. K562 cells were transduced with either a TXNDC15, MARCHF6, or non-targeting control sgRNA. Cells were lysed and proteins were enzymatically digested, and the resulting peptides were purified and analyzed via mass spectrometry. Two identical experiments with 4 technical replicates each were compared and proteins that showed significant changes in abundance upon depletion of TXNDC15 or MARCHF6 are displayed (i.e. a log2 abundance ratio <-0.5 and >0.5 and more than 1 different peptide was detected). Proteins displayed in teal were found at increased levels in knockdown compared to wildtype cells, while proteins labeled in tan were at decreased levels. Proteins in bold indicate membrane proteins, those in italics are peripheral membrane proteins, and soluble proteins are shown in standard font. (B) Analysis of the stability of several additional TXNDC15 substrates. Membrane proteins with varying topologies were analyzed using our ratiometric fluorescent reporter in HEK293T wt versus TXNDC15 knockdown cells. These experiments identified additional GET1-like and TMEM231-like substrates, either degraded or stabilized by TXNDC15. (C) To determine if TXNDC15 used a concerted mechanism across all identified substrates, we performed a rescue assay with the TXNDC15 mutants described in Figure 3A. Rescue efficiencies for the indicated substrates were calculated relative to wt and TXNDC15 kd cells. Mean and standard deviation are shown (n = 3). The dashed line at 1 indicates where mutants able to fully rescue either the degradation or stabilization phenotype would be expected. See also Figure S6 and Table S4.

Second, because many endogenous membrane proteins are poorly detected by mass spectrometry, we also used our knowledge of the molecular features of GET1 and TMEM231 to rationally identify a panel of potential substrates (Figure 5B). For example, we reasoned that other subunits that contained multiple TMs and cytosolic domains might behave like GET1 (e.g. GET2 and EMC5). Equally, integral membrane proteins with globular lumenal domains might behave like TMEM231 (e.g. the nicotinic acetylcholine subunits, CHRNA9 and CHRNA10, members of the physiologically important cys-loop receptor family).^57^ Comparison of these substrates to a panel of other membrane protein subunits suggested that indeed, presence of a soluble cytosolic domain was a common feature of substrates that rely on TXNDC15 for their degradation by MARCHF6. A globular lumenal domain was also common to substrates that were stabilized by TXNDC15, suggesting that it more generally limited access to MARCHF6 for substrates with this feature. We also observed that the majority of GET1-like substrates of TXNDC15 were ER-resident proteins, while TMEM231-like substrates were exported to the plasma membrane. However, not all ER-resident or exported proteins that are MARCHF6 dependent are substrates of TXNDC15, so specific molecular features are also critical to confer TXNDC15 dependence.

Third, to test if the mechanism of TXNDC15 function was consistent across all identified substrates, we performed a similar mutational experiment previously used to identify the domains of TXNDC15 required for its activity (Figure 5C). As we observed for GET1 and TMEM231, we found that the lumenal domain of TXNDC15, provided that it was anchored at the membrane, was both necessary and sufficient for its degradation and stabilization effects on all tested substrates. We therefore concluded that TXNDC15 used a consistent mechanism for its role in regulating protein homeostasis in the ER across the proteome.

### Patient mutations to TXNDC15 cause inappropriate degradation of TMEM231 in the ER and thereby defects in ciliogenesis

Finally, we sought to directly establish the connection between TXNDC15’s function in modulating MARCHF6 activity in the ER to its role in ciliogenesis at the plasma membrane. Earlier work has established that depletion of TXNDC15 and TMEM231 in cells leads to defects in cilia function, morphology, and signaling.^28,42,58^ For these reasons, mutations in both proteins in patients lead to a severe ciliopathy known as Meckel-Gruber syndrome, which is characterized by malformation of the central nervous system, polycystic kidneys, and polydactyly that together lead to death in utero or soon after birth.^38–40,59^ In the case of TMEM231, its critical role in organizing the MKS complex at the ciliary transition zone provides a clear explanation for its connection to disease. However, for TXNDC15, how an ER protein might regulate ciliogenesis was not clear.

To further define this connection, we selected a patient mutation characterized by a five amino acid deletion to the TXNDC15 thioredoxin domain (Figure 6A).^38^ We found that expression of wildtype, but not the mutant TXNDC15, was able to rescue defects in cilia formation and morphology in human RPE1 and mouse NIH/3T3 cells consistent with earlier experiments (Figures 6B-E, S7A-E).^41,42^ However, loss of MARCHF6 did not affect ciliogenesis, and its depletion rescued the TXNDC15 knockdown phenotype (Figure 6F-G, 2H). Therefore, the effects of TXNDC15 on ciliogenesis must occur via its role in regulating MARCHF6. Indeed, while this patient mutation of TXNDC15 accumulated to higher levels than wildtype in cells (Figures S7F-G), we showed that its binding to MARCHF6 is partially disrupted (Figure S7H), explaining the loss of its protective function in the ER.

**Figure 6.**
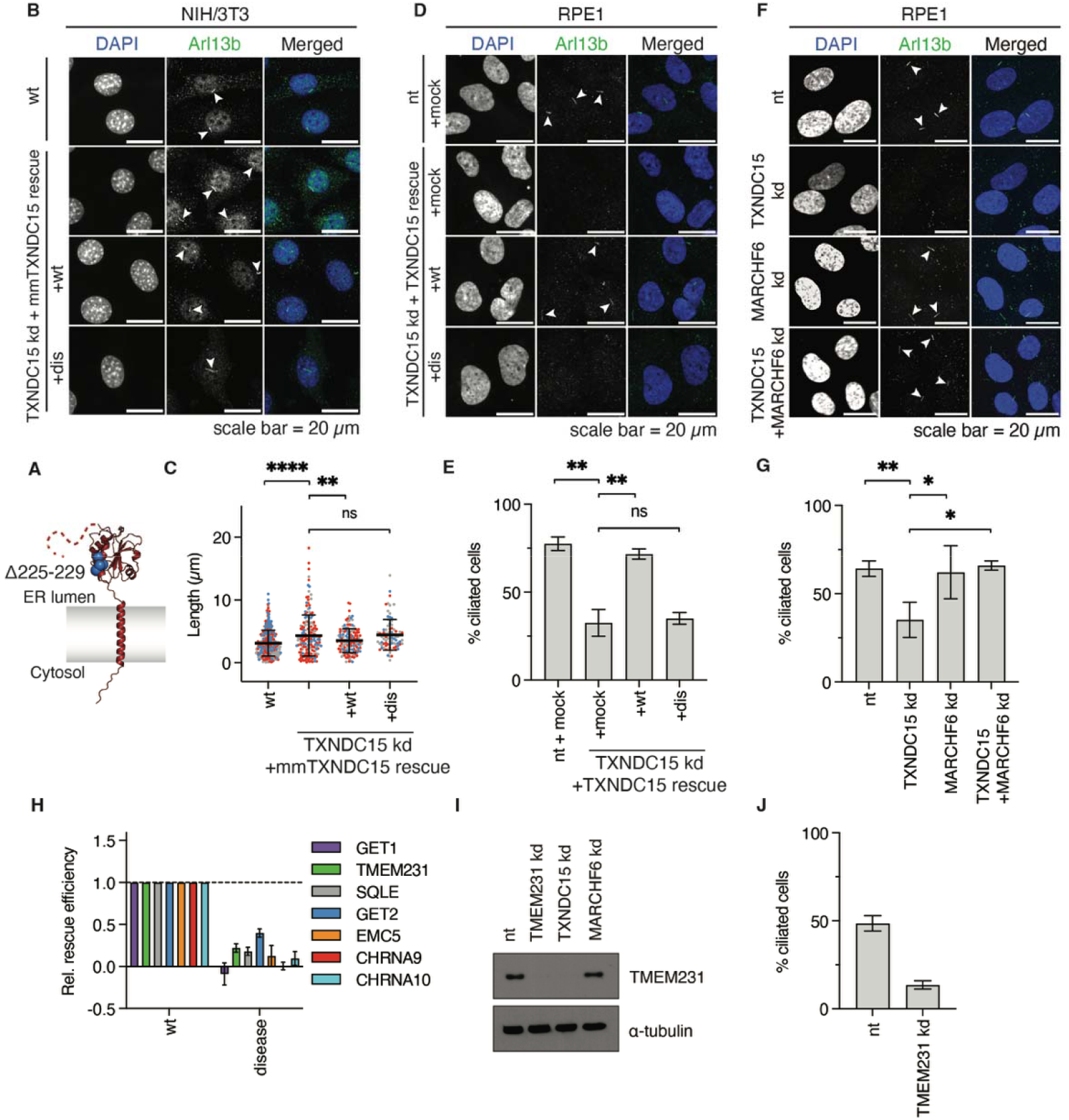
Patient mutations to TXNDC15 destabilize TMEM231 in the ER, leading to defects in ciliogenesis. (A) AlphaFold3 predicted model of TXNDC15. A patient mutation to TXNDC15 that causes the ciliopathy Meckel-Gruber syndrome involves a deletion within the lumenal domain (hs: Δ225-229^38^, mm: Δ209-213) and is highlighted as blue spheres. (B,C) Confocal microscopy of cilia in serum starved murine NIH/3T3 cells. Cilia length was determined from either wildtype (wt), TXNDC15 knockdown (kd), or TXNDC15 kd cells expressing wt murine TXNDC15 or the murine homolog of the human TXNDC15 ciliopathy-mutant (dis, Δ209-213). DAPI (blue) stains nuclear DNA and Arl13b (green) is used as ciliary marker. (C) Cilia lengths of NIH/3T3 cells are reported with the average and standard deviation for wt cells (338 cilia), TXNDC15 kd (182 cilia), murine TXNDC15 rescue (147 cilia) and murine TXNDC15 ciliopathy mutant rescue (85 cilia). Colors represent different replicates (n=3), percentage of ciliated cells is shown in Figure S7E, where we also observed a decrease in cilia in TXNDC15 kd cells. Statistical significance was determined using Welch’s t-test (p(nt/TXNDC15 kd) < 0.0001; p(TXNDC15 kd/TXNDC15wt rescue) = 0.005; p(TXNDC15 kd/TXNDC15dis rescue) = 0.81). (D,E) Confocal microscopy of cilia in serum starved RPE1 wt cells and TXNDC15 kd cells with either mock (BFP only), or wt TXNDC15 rescue, or TXNDC15 disease-mutant rescue (dis, Δ225-229). (E) Percent ciliated RPE1 cells, determined by quantifying the number of cilia compared to the total number of cells. Mean and standard deviation for 3 separate slides per sample are shown (nt kd: 759 cells; TXNDC15 kd: 742 cells; TXNDC15 wt rescue: 488 cells; TXNDC15(dis) rescue: 520 cells). Statistical significance was determined using Welch’s t-test (p(nt/TXNDC15 kd) = 0.0029; p(TXNDC15 kd/TXNDC15wt rescue) = 0.006; p(TXNDC15 kd/TXNDC15dis rescue) = 0.64). (F,G) Confocal microscopy of cilia in serum starved RPE1 wt cells, TXNDC15 kd cells, MARCHF6 kd cells or TXNDC15 + MARCHF6 double kd cells. (G) As in (E) for samples shown in (F). (nt kd: 250 cells; TXNDC15 kd: 126 cells, MARCHF6 kd: 268 cells; TXNDC15 + MARCHF6 kd: 241 cells). Statistical significance was determined using Welch’s t-test (p(nt/TXNDC15 kd) = 0.0055; p(TXNDC15 kd/MARCHF6 kd) = 0.029; p(TXNDC15 kd/Double kd) = 0.0064). (H) Analysis of the activity of the ciliopathy-causing mutation (Δ225-229) of TXNDC15 on a panel of TXNDC15 substrates in HEK293T. Rescue experiments in wt and TXNDC15 depleted cells were carried out using the indicated reporters as described in Figure 5B and analyzed using flow cytometry. (I) RPE1 cells were treated with either a TMEM231, TXNDC15, MARCHF6, or non-targeting control (nt) sgRNA. After an eight-day knockdown, cells were harvested and endogenous TMEM231 levels were analyzed by western blot. While depletion of TXNDC15 decreases TMEM231 levels to a similar extent as TMEM231 kd, depletion of MARCHF6 maintains the steady-state levels of TMEM231. (J) Percent of ciliated RPE1 cells upon TMEM231 kd quantified as described in (E). Error bars represent the standard deviation of three separate slides (nt kd: 678 cells; TMEM231 kd: 654 cells). See also Figure S7.

Consistent with this model, using our reporter system, we found that the patient mutant of TXNDC15 was unable to rescue both its quality control and stabilization effects on a panel of substrates (Figures 6H, S7I-K). Notably, this disease mutant also cannot rescue the destabilization of TMEM231, which is depleted to a similar extent upon loss of TXNDC15 as in a direct TMEM231 knockdown (Figure 6I-J, S7L). Furthermore, expression of TMEM231 alone was able to partially rescue the ciliogenesis defect observed upon TXNDC15 depletion (Figure S7M). We therefore concluded that in patients, loss of TXNDC15 regulation of MARCHF6 leads to the inappropriate degradation of TMEM231, providing a biochemical explanation for how mutations to the ER-resident TXNDC15 cause Meckel Gruber syndrome.

## DISCUSSION

Taken together, our data support a model in which TXNDC15 is an ERAD factor that regulates membrane protein levels in the ER through its tuning of MARCHF6 activity in two ways (Figure 7). For proteins with globular lumenal domains like TMEM231, mutations or deletions to TXNDC15 that disrupt its binding and/or occupancy at MARCHF6 lead to their inappropriate recruitment to the E3 ubiquitin ligase in the ER. This leads to their ubiquitination, extraction, and proteasome-dependent degradation. Indeed, our data suggested that TMEM231 levels may normally be regulated by a MARCHF6-independent pathway, and it is only when the protective effects of TXNDC15 are lost or diminished, that TMEM231 is aberrantly recognized by MARCHF6. The degradation of TMEM231 at the ER depletes the protein from cilia, resulting in defects in the transition zone that lead to deficits in ciliary morphology and function, causing Meckel-Gruber syndrome. Our data suggest that this aberrant decrease in TMEM231 levels is sufficient to cause defects in ciliogenesis and therefore is the critical link between TXNDC15 and ciliopathies. Indeed, previous studies have observed that depletion of both TMEM231 and TXNDC15 have identical effects on the integrity of the MKS complex and thereby the localization of TMEM67.^28,38,41^ However, without TXNDC15 to regulate MARCHF6, we observed a more general dysregulation of proteostasis at the ER, which could also affect the ciliary proteome and further contribute to the severity of disease. Importantly, the genetic link between TXNDC15 and ciliogenesis in patients validates the role of TXNDC15 on endogenous substrates in an organismal model.

**Figure 7.**
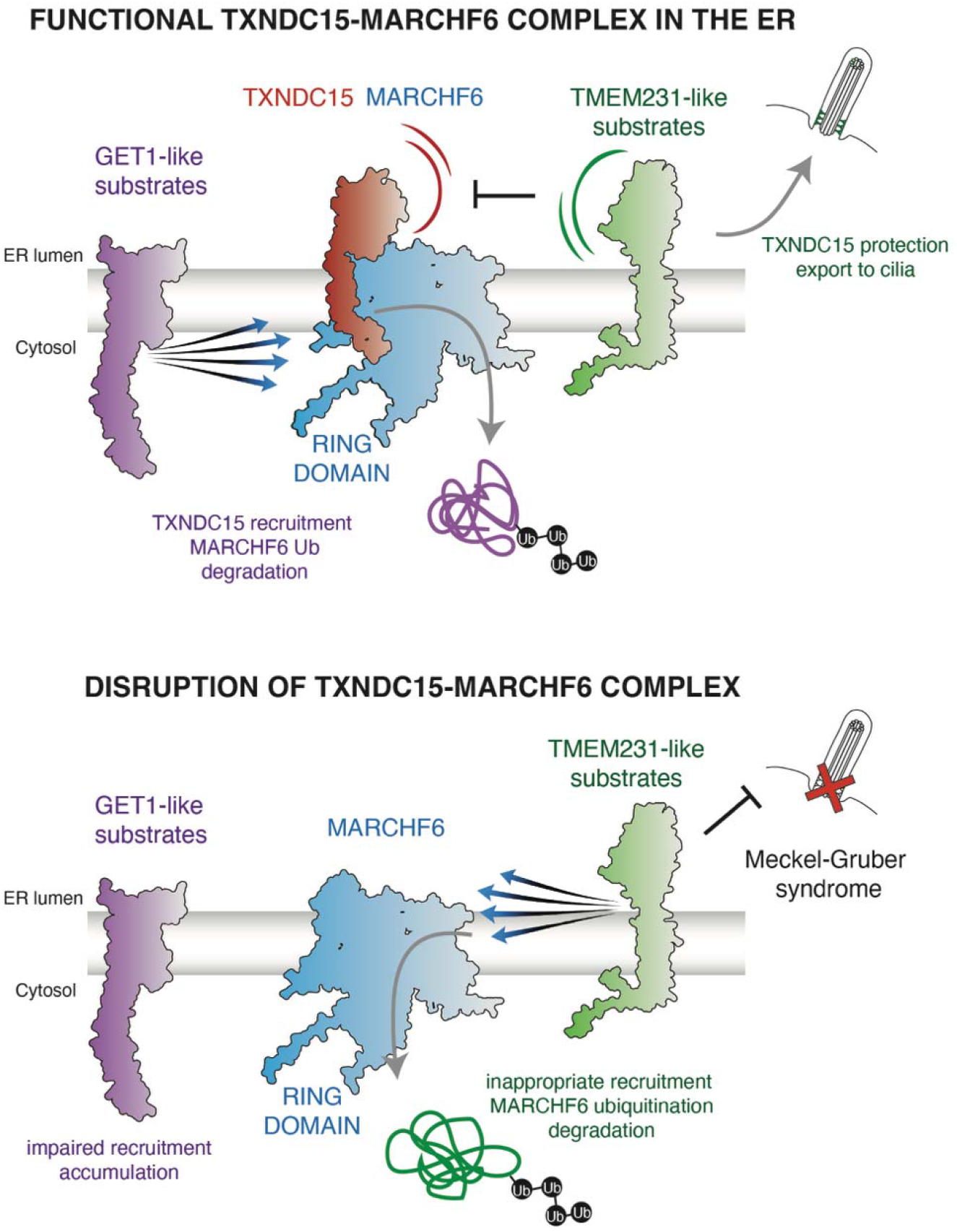
Proposed model for TXNDC15-mediated regulation of protein homeostasis in the ER via modulation of MARCHF6. Wildtype TXNDC15 forms a functional complex with the E3 ubiquitin ligase MARCHF6. Binding of TXNDC15 facilitates the recruitment, ubiquitination, retrotranslocation, and degradation of GET1-like substrates, while also inhibiting the inappropriate recruitment of TMEM231-like substrates. TMEM231 can then be trafficked to the plasma membrane where it supports the formation of functional cilia. However, if TXNDC15-MARCHF6 complex formation is disrupted, through either deletion or loss of function mutations to TXNDC15, recruitment and thereby degradation of GET1-like substrates is impaired. In the absence of TXNDC15 binding at MARCHF6, its protective effect is lost and TMEM231-like substrates can be inappropriately recruited and ubiquitinated by MARCHF6. Severe reduction of TMEM231 levels in the ER and at the plasma membrane disrupt ciliogenesis, leading to Meckel-Gruber syndrome.

Conversely, for membrane subunits with soluble cytosolic domains like GET1, TXNDC15 enhances their recruitment to MARCHF6, leading to their ubiquitination, extraction, and degradation. We observed that recognition of this class of substrates relied on a composite degron within the membrane and cytosol, reminiscent of earlier work on Doa10 (the yeast homolog of MARCHF6) that recognized degrons spanning the membrane and lumen.^51^ In the absence of TXNDC15, this class of substrates is less efficiently recruited to MARCHF6, leading to their aberrant accumulation. Because we found that many substrates that relied on TXNDC15 for their degradation were ER-resident, it is possible that their longer dwell time in the ER necessitates inefficient recognition by MARCHF6. TXNDC15 could be fine-tuning this interaction, regulating the kinetics of their turnover to allow sufficient time for complex assembly.

Given these observations, we can begin to hypothesize how binding of TXNDC15 might modulate MARCHF6 activity in these two opposing ways. For example, AlphaFold3 predicts that the lumenal domain of TXNDC15 binds near the pore of MARCHF6, which could partially occlude access from the ER lumen for substrates with globular domains like TMEM231. Equally, AlphaFold3 predicts binding of TXNDC15 could change the degree of opening of MARCHF6’s flexible lateral gate, which could facilitate access from within the membrane to its cytosolic RING domain for substrates like GET1.^56^ These hypotheses would be consistent with recent structures of Doa10 that proposed that substrates must access its central cavity and lateral tunnel for their ubiquitination.^60,61^ Notably, we observed more modest effects of TXNDC15 on substrate recruitment to MARCHF6 compared to their ubiquitination, suggesting TXNDC15 could play an additional role in correctly orienting substrates like GET1 at the RING domain of MARCHF6. Regardless, it is clear that the expression and levels of TXNDC15, and thereby its occupancy at MARCHF6, controls protein homeostasis in the ER.

Global analysis of TXNDC15 across tissues suggests that there is substantial variability in its expression.^62^ TXNDC15 levels may therefore be titrated to regulate membrane subunit levels depending on the unique requirements of particular cell types. Our experiments in K562s cells suggested that at endogenous levels of TXNDC15, MARCHF6 is not fully occupied, and therefore overexpression of TXNDC15 can drive an additional protective effect. In contrast, the highest levels of TXNDC15 are reported in the choroid plexus, a region of the brain that relies on highly ciliated epithelial cells that regulate cerebrospinal fluid production. Choroid plexus cysts have been reported in Meckel-Gruber syndrome,^63^ underscoring the critical role of TXNDC15 in regulating the ciliary proteome and how its dysregulation leads to the pathogenesis of disease.

These observations have two important consequences, both for our understanding of ciliary protein biogenesis and more generally membrane protein quality control. First, this role of ERAD in regulating ciliary subunit levels, implies that some aspects of quality control and very likely assembly of ciliary complexes occur in the ER. To what extent membrane protein complex assembly occurs prior to trafficking, particularly of architecturally complicated structures like the MKS complex, is an important area for future research.

Second, the molecular characterization of TXNDC15’s regulation of MARCHF6 provides an explanation for how the cell can rely on a small number of E3 ligases to surveil the enormous diversity within the human membrane proteome. The use of adaptors that act as molecular switches to mediate or inhibit interactions with specific classes of substrates is an efficient strategy to regulate the repertoire of a single E3 ligase.

Variable expression of such adapters in particular cell types could be used to tune membrane protein homeostasis. Indeed, this is analogous to the many adaptors that have been characterized in the cytosol that function with HSP70, VCP, and Skp1-Cullin E3 ubiquitin ligases to maintain proteostasis.^64–66^ However, it is clear that while a pathway may specialize in a particular region and set of substrates, there is substantial overlap within the protein quality control network. Compensatory and redundant pathways are critical for robustness and may be particularly important in post-mitotic cell types that must survive throughout an eighty-to ninety-year lifespan.

## MATERIALS AND METHODS

### EXPERIMENTAL MODEL DETAILS

#### Cell lines

HEK293T cells were cultured in Dulbecco’s Modified Eagle Medium (DMEM; 11965092, Thermo Scientific, USA) supplemented with 10% fetal bovine serum (FBS; S11150, Bio-Techne, USA), 2 mM glutamine. RPE1 cells were cultured in Dulbecco’s Modified Eagle Medium/Nutrient Mixture F-12 (DMEM/F-12; 11320033, Thermo Scientific, USA) supplemented with 10% FBS, 2 mM glutamine. NIH/3T3 cells were cultured in DMEM supplemented with 10% FBS, 32 mM glutamine, 4.5 g/L D-glucose and 110 mg/L sodium pyruvate (10569010, Thermo Scientific, USA). K562 cells containing KRAB-BFPdCas9^33^ were cultured in RPMI-1640 with 25 mM HEPES, 2.0 g/L NaHCO3, and 0.3 g/L L-glutamine (SH30255.01, Cytiva, USA) supplemented with 10% Tet System Approved FBS (S10350, Bio-Techne, USA), 2 mM glutamine, 100 units/mL penicillin, and 100 μg/mL streptomycin. Expi293 cells were cultured in Expi293 Expression Medium (A1435101, Thermo Scientific, USA). K562 and Expi293 cells were maintained between 0.25 × 10^6^ –1 × 10^6^ cells/mL. Adherent cells were detached from culture dishes (trypsinized) using Trypsin-EDTA (0.25%), phenol red (25200056, Thermo Scientific, USA). HEK293T, RPE1, NIH 3T3, and K562 cells were grown at 37°C and 5% CO_2_. Expi293 cells were grown at 37°C and 8% CO_2_ with shaking at 125 rpm. Cell lines were not authenticated.

#### Biochemical preparations

GFP-nanobodies were purified from the E. coli strain AVB101 (CVB101, Avidity, USA) in Superbroth media (35 g/L tryptone, 20 g/L yeast extract, 5 g/L NaCl). ALFA-nanobodies were purified from E. coli Rosetta-gami2 (71350-3, Novagen/Millipore-Sigma, USA). SUMO proteases were purified in E. coli NEBExpress cells (C3037I, New England Biolabs, USA).

## METHOD DETAILS

### Plasmids

#### Protein sequences

The sequences used in cell-based and in vitro experiments were derived from UniProtKB/SwissProt. These include: guided entry of tail-anchored proteins factor 1 (GET1; O00258), guided entry of tail-anchored proteins factor 2 (GET2; P49069), transmembrane protein 231 (TMEM231; Q9H6L2), SARS-CoV-2 membrane glycoprotein M (VME1_SARS2; P0DTC5), vesicle associated membrane protein 2 (VAMP2; P63027-1), translocon-associated protein subunit alpha (SSR1/SSRA/TRAPα; P43307), ER membrane protein complex subunit 3 (EMC3; Q9P0I2), thioredoxin domain-containing protein 15 (TXNDC15; human: Q96J42, mouse: Q6P6J9), E3 ubiquitin-protein ligase MARCHF6 (MARCHF6; O60337), T-cell surface glycoprotein CD4 (CD4; P01730), villin-1 (βV; chicken: P02640), squalene monooxygenase (SQLE; Q14534), ER membrane protein complex subunit 5 (EMC5/MMGT1; Q8N4V1), neuronal acetylcholine receptor subunit alpha-9 (CHRNA9; Q9UGM1: Variant p.Asn442Ser), neuronal acetylcholine receptor subunit alpha-10 (CHRNA10; Q9GZZ6), transmembrane protein 17 (TMEM17; Q86X19), transmembrane protein 67 (TMEM67; Q5HYA8), transmembrane protein 107 (TMEM107; Q6UX40), transmembrane protein 216 (TMEM216; Q9P0N5), transmembrane protein 237 (TMEM237; Q96Q45), tectonic-2 (TCTN2; Q96GX1), monoacylglycerol lipase (ABHD2; P08910).

#### General plasmid backbones

The 2nd generation lentiviral packaging plasmid pCMV-VSV-G was a gift from Bob Weinberg (Addgene plasmid # 8454). The 2nd generation lentiviral packaging plasmid psPAX2 was a gift from Didier Trono (Addgene plasmid # 12260). The pHAGE2 lentiviral transfer plasmid was a gift from Magnus A. Hoffmann and Pamela Bjorkman. The dual guide lentiviral vector pJR103 was a gift from Jonathan Weissman (Addgene plasmid # 187242). The SFFV-tet3G backbone was used for K562 cell expression during CRISPRi screens.^67^ For transduction of NIH 3T3 cells, we used the plasmid backbone pMCB306 (Addgene plasmid # 89360), which was a gift from Michael Bassik. RFP is used in the text and figures and refers to the mCherry variant that was used in this study; GFP refers to the EGFP variant. The ratiometric fluorescence GFP:RFP reporter system was used as previously described.^9^

#### Inducible reporter construct in K562 cells

The fluorescent reporter used for genome wide CRISPRi screens contained either GET1 or TMEM231 cloned downstream of the doxycycline-inducible TET3G promoter with a C-terminal GFP fusion followed by a 2A ribosomal skipping sequence (2A) and RFP normalization marker. The same construct was used for reporter assays in K562s using an inducible reporter.

#### Constitutive reporter construct in K562 cells

For select reporter assay experiments, the constitutive reporter construct of GET1 was cloned downstream of an EF1α promoter with a C-terminal GFP fusion followed by a 2A and RFP in the pHAGE2 lentiviral vector.

#### Reporter constructs in HEK293T cells

For reporter assays, the following reporter constructs were cloned downstream of a CMV promoter with a C-terminal GFP fusion followed by a 2A and RFP in the pHAGE2 lentiviral vector: GET1, EMC3, EMC5, M, SQLE, TRAPα, TMEM17, TMEM107, TMEM216, TMEM231, TMEM237, TCTN2. Chimeras of TMEM237, GET1, TMEM231 and VHP were cloned in the same fluorescence cassette but expressed under a EF1a promoter.

Due to their topology, CHRNA9 and CHRNA10 were labelled using an internal GFP in the cytoplasmic loop between TMs 3 and 4, which was inserted after amino acid 412 for CHRNA9 and after amino acid 391 for CHRNA10 as previously described. Previous experiments demonstrated that this internal labeling strategy did not perturb protein function.^68^ In each case, the protein of interest was then followed by a 2A skipping sequence and the normalization marker RFP.

To ensure the fluorescent tags were positioned in the cytosol to avoid perturbing insertion efficiency, GET2 was cloned downstream of a CMV promoter with an N-terminal GFP fusion followed by a 2A and RFP in the pHAGE2 lentiviral vector. The tail-anchored membrane protein reporter VAMP2 was labelled on the N-terminus using the GFP-2A-RFP cassette in the pHAGE2 lentiviral vector. In this case GFP was used as translation normalization marker.

For overexpression of GET2 in the presence of GET1-GFP-2A-RFP, GET2 was expressed along with a BFP control separated by a 2A site.

For the split fluorescence reporter assay, monomeric neon green 2 (mNG2) was used as a fluorophore.^69^ The eleventh beta sheet of the fluorescent protein (mNG2(11)) was fused to the C-terminus of GET1, SQLE or ABHD2 followed by a 2A sequence and the translation normalization marker RFP. The first ten beta sheets (mNG2(1-10)) were expressed in the cytosol.

#### GET2 overexpression in HEK293T cells

For overexpression of GET2, the protein of interest was cloned downstream of a CMV promoter and followed by a 2A-BFP cassette. This allowed to select for GET2 expressing cells.

#### TXNDC15 rescue constructs in HEK293T cells

The TXNDC15 rescue constructs were cloned downstream of an EF1α promoter with a C-terminal ALFA fusion followed by a 2A and BFP-3xFLAG in the pHAGE2 vector. The ALFA epitope (PSRLEEELRRRLTEP) has been previously described.^70^ Mutant variants of TXNDC15 were cloned such that the native TXNDC15 TM (residues 322-342) was replaced with the TM of CD4 along with flank residues (GGPGMALIVLGGVAGLLLFIGLGIFFGPGG), the ER lumenal domain of TXNDC15 (residues 33-315) was replaced by the globular, inert, βVHP domain (LSDEDFKAVFGMTRSAFANLPLWKQQNLKKEKGLF), or only the thioredoxin domain residues 1-321 fused to a C-terminal ER-retention sequence KDEL.^71^ Deletion of the predicted disordered region of TXNDC15 was performed to preserve the N-terminal signal sequence, deleting amino acids 36-163). Mutant variants of TXNDC15 with an exogenous linker between the lumenal thioredoxin domain and the TM were cloned such that flexible (GS)_8_ or rigid (EAAAK)_5_ repeats were inserted at TXNDC15 residue 322. The rigid linker motif (EAAAK)_n_ (n = 3-5) has been previously characterized to rigidly increase the distance between protein domains.^72^ To ensure the helix was continuous, amino acids 322 to 324 were deleted from the rigid linker. TXNDC15 cysteine mutants (C220S, C210A) were cloned using site-directed mutagenesis.

TXNDC15(dis) is a TXNDC15 construct with residues 225-229 deleted using site-directed mutagenesis, as described previously to cause Meckel-Gruber syndrome.^38^ In all TXNDC15 rescue variants, its native signal sequence remained intact.

#### TXNDC15 and MARCHF6 rescue constructs in K562 cells

TXNDC15 and MARCHF6 constructs were cloned downstream of a PGK promoter with a C-terminal ALFA fusion followed by a 2A and BFP-3xFLAG in the pHAGE2 vector. Double rescue constructs contained MARCHF6-ALFA-2A-TXNDC15-ALFA-2A-BFP-3xFLAG. As previously mentioned, a point mutation was introduced to MARCHF6 to create the catalytically inactive form MARCHF6(C9A).

#### TXNDC15 rescue constructs in murine NIH 3T3 cells

To rescue the TXNDC15 kd phenotype in NIH 3T3 cells in immunofluorescence experiment, the mmTXNDC15 sequence was fused to a C-terminal ALFA-tag followed by a 2A and RFP in the pHAGE2 vector downstream of an EF1α promoter. To rescue the TXNDC15 kd phenotype in NIH 3T3 cells in the rescue reporter assay, the mmTXNDC15 sequence was fused to a C-terminal ALFA-tag followed by a 2A and BFP-3x-FLAG in the pHAGE2 vector downstream of an EF1α promoter. The ciliopathy mutant (mmTXNDC15(dis)) was created by deleting amino acids 209-213 which are equivalent to the human disease mutation as described above.

#### TXNDC15 rescue constructs in RPE1 cells

TXNDC15 rescue constructs (wt and dis) used for immunofluorescence were cloned downstream of an EF1α promoter with a C-terminal ALFA fusion followed by a 2A and RFP in the pHAGE2 vector. TXNDC15 rescue constructs used for the TXNDC15 rescue assay are the same constructs that were used in HEK293T cells as described above.

#### Constructs for stable expression in Expi293 cells

For exogenous expression of MARCHF6 to be used for immunoprecipitation-mass spectrometry in Figure 2C, we used the catalytically inactive variant MARCHF6(C9A) – this mutation resides in its RING domain – as MARCHF6 had previously been shown to catalyze its own degradation.^50^ The sequence of MARCHF6(C9A) was cloned downstream of a CMV promoter with a C-terminal TEV cleavage site (ENLYFQG) followed by a GFP-2A-RFP in the pHAGE2 vector. Construct for “MARCHF6-GFP” is the MARCHF6(C9A)-TEV-GFP-2A-RFP as described above. The TXNDC15 expression construct, “TXNDC15-ALFA”, was cloned downstream of a CMV promoter with a C-terminal ALFA followed by a 2A and BFP-3xFLAG in the pHAGE2 vector.

#### Constructs for in vitro translations

Constructs for expression of GET1 and TMEM231 in rabbit reticulocyte lysate (RRL) were based on the SP64 vector (Promega) using an SP6 promoter.

#### CRISPRi knockdown guides

Programmed single and dual guides were generated to assay depletion of one or two genes using the CRISPRi system.^73^ The following sgRNA protospacer sequences were used to clone into pLG1 (single guides) or pJR103 (dual guides):

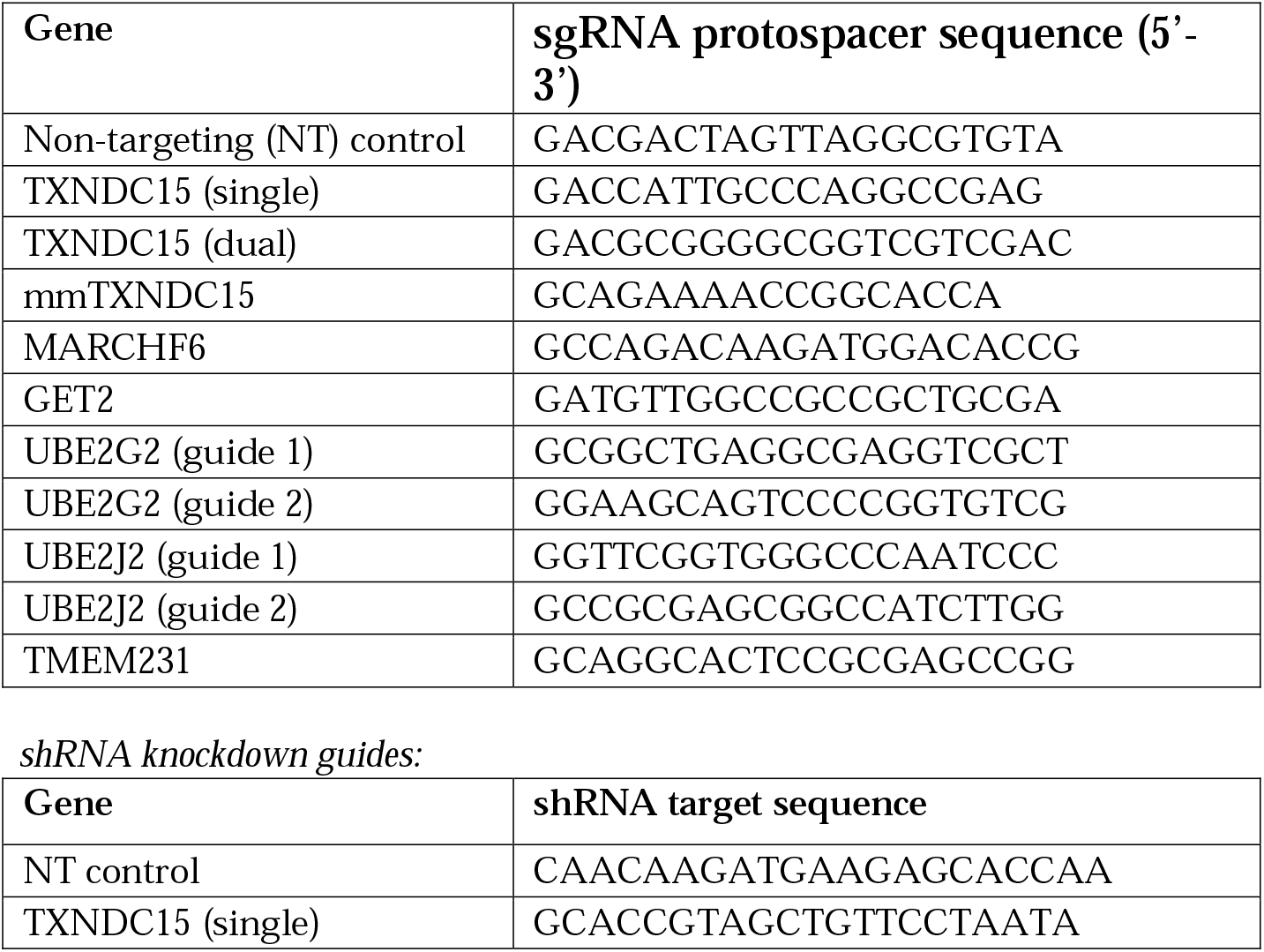

All plasmids are available upon request.

### Antibodies for western blotting and immunofluorescence (catalog number, manufacturer, RRID; dilution)

*Primary antibodies* mouse monoclonal anti-α-tubulin (T9026, Sigma-Aldrich, USA, RRID:AB_477593; 1:2,000); rabbit anti-Sec61β (gift from Ramanujan Hegde; 1:50-1:1,000); rabbit polyclonal anti-GFP (A11122, Thermo Scientific, USA, RRID:AB_221569; 1:1,000); rabbit polyclonal anti-Arl13b (17711-1-AP, Proteintech, USA, RRID:AB_2060867; 1:100); mouse monoclonal anti-Arl13b (66739-1-Ig, Proteintech, USA, RRID:AB_2882088; 1:100); mouse monoclonal anti-acetylated-α-tubulin (66200-1-Ig, Proteintech, RRID:AB_2722562; 1:100); rabbit anti-polyubiquitin-B (SPA-200F, Enzo Life Sciences, RRID:AB_312481; 1:1,000); rabbit polyclonal anti-TMEM231 (23731-1-AP, Proteintech, USA, RRID: AB_2879312; 1:1,000); rabbit anti-Bag6 (gift from Ramanujan Hegde; 1:30,000); rabbit anti-MMGT1 (A305-833A, Bethyl, RRID:AB_2890207; 1:1,000); rabbit anti-SYVN1 (A302-946A, Bethyl, RRID:AB_10690984; 1:1,000).

*Secondary antibodies used for western blotting:* goat anti-mouse-HRP (172-1011, BioRad, USA, RRID:AB_11125936; 1:2,500 or 1:5,000); goat anti-rabbit-HRP (170-6515, BioRad, USA, RRID:AB_11125142; 1:2,500).

*Secondary antibodies used for immunofluorescence:*goat anti-mouse IgG Alexa Fluor 488 (115-545-003, Jackson ImmunoResearch Inc, USA, RRID:AB_2338840; 1:500); goat anti-rabbit IgG Alexa Fluor 488 (111-545-003, Jackson ImmunoResearch Inc, USA, RRID:AB_2338046; 1:500); goat anti-mouse IgG Alexa Fluor 647 (115-605-003, Jackson ImmunoResearch Inc, USA, RRID:AB_2338902; 1:500).

#### Others

anti-FLAG-HRP (A8592, Sigma-Aldrich, USA, RRID:AB_439702; 1:10,000); anti-ALFA nanobody-HRP (this paper; 1:10,000-80,000), GFP antibody Alexa Fluor 488 (A-21311, Thermo Scientific, USA, RRID:AB_221477; 1:100).

### Conjugation of the ALFA nanobody to HRP for western blotting

ALFA nanobody was coupled to horseradish peroxidase (HRP)-maleimide via an engineered C-terminal cysteine, as previously described.^74^ Briefly, equimolar amounts of the ALFA nanobody and EZ-Link™ maleimide activated HRP (31485, Thermo Scientific, USA) are incubated for 1 hour at room temperature (RT).

### Visualization of western blots

Western blot signal was developed after 1 minute incubation with SuperSignal™ West Pico PLUS Chemiluminescent Substrate (34580, Thermo Scientific) or SuperSignal™ West Femto Maximum Sensitivity Substrate (34094, Thermo Scientific).

### Lentiviral production

Lentivirus was produced by co-transfection of packaging plasmid psPAX2 and envelope plasmid VSV.G, along with a transfer plasmid of interest, into HEK293T cells using TransIT-293 transfection reagent (2705, Mirus, USA) as per manufacturer’s protocol. Lentivirus was harvested 48 hours after transfection and stored at -80°C for future use.

### Cell culture conditions

| **Cell Lines** | **Source** |
|---|---|
| K562 KRAB-BFP-dCAS9 | <sup>33</sup> |
| HEK293T/17 | CRL-11268, ATCC, USA |
| K562-CRISPRi-Tet-ON-(tet-GET1-GFP-2A-RFP, KRAB(Kox1)) | This study |
| K562-CRISPRi-Tet-ON-(tet-TMEM231-GFP-2A-RFP, KRAB(zim3)) | This study |
| HEK293T TXNDC15 stable knockdown | This study |
| HEK293T MARCHF6 stable knockdown | This study |
| HEK293T SS-GFP-TXNDC15 knockin | This study |
| HEK293T MARCHF6-TEV-ALFA-2A-BFP knockin | This study |
| HEK293T SS-GFP-TXNDC15 MARCHF6-TEV-ALFA-2A-BFP knockin | This study |
| HEK293T MARCHF6-TEV-ALFA-2A-BFP knockin TXNDC15 stable knockdown | This study |
| Expi293 MARCHF6(C9A)-GFP | This study |
| Expi293 MARCHF6(C9A)-GFP TXNDC15-ALFA | This study |
| RPE1 KRAB-BFP-dCAS9 | <sup>67</sup> |
| NIH 3T3 Cas9 | Gift from Maxence Nachury |
| NIH 3T3 Cas9 TXNDC15 stable knockdown | This study |

HEK293T cells were cultured in Dulbecco’s Modified Eagle Medium (DMEM; 11965092, Thermo Scientific, USA) supplemented with 10% fetal bovine serum (FBS; S11150, Bio-Techne, USA), 2 mM glutamine. RPE1 cells were cultured in Dulbecco’s Modified Eagle Medium/Nutrient Mixture F-12 (DMEM/F-12; 11320033, Thermo Scientific, USA) supplemented with 10% FBS, 2 mM glutamine. NIH 3T3 cells were cultured in DMEM supplemented with 10% FBS, 32 mM glutamine, 4.5 g/L D-glucose and 110 mg/L sodium pyruvate (10569010, Thermo Scientific, USA). K562 cells containing KRAB-BFPdCas9^33^ were cultured in RPMI-1640 with 25 mM HEPES, 2.0 g/L NaHCO3, and 0.3 g/L L-glutamine (SH30255.01, Cytiva, USA) supplemented with 10% Tet System Approved FBS, 2 mM glutamine, 100 units/mL penicillin, and 100 μg/mL streptomycin. Expi293 cells were cultured in Expi293 Expression Medium (A1435101, Thermo Scientific, USA). K562 and Expi293 cells were maintained between 0.25 × 10^6^ –1 × 10^6^ cells/mL. Adherent cells were detached from culture dishes (trypsinized) using Trypsin-EDTA (0.25%), phenol red (25200056, Thermo Scientific, USA). HEK293T, RPE1, NIH 3T3, and K562 cells were grown at 37°C and 5% CO_2_. Expi293 cells were grown at 37°C and 8% CO_2_ with shaking at 125 rpm.

For preparations of ER microsomes using HEK293T wt cells and those with endogenously tagged GFP-TXNDC15, these cell lines were adapted for growth in suspension. For adaptation, HEK293T cell lines were trypsinized, pelleted, and resuspended in Expi293 Expression Medium, and maintained between 0.25 × 10^6^ – 1 × 10^6^ cells/mL.

### Endogenously tagged cell line construction and stable knockdown cell lines

Tagging of TXNDC15 at its endogenous locus in HEK293T cells was performed as previously described.^75^ Briefly, to endogenously tag TXNDC15 with a GFP positioned between its signal sequence and the mature N-terminus, a protospacer within exon two was cloned into pX459 (CCTGGCCCATCGGGTCATG). A vector encoding GFP and 800 bp of homology on either side of the cut site of the TXNDC15 locus was ordered from TWIST Biosciences. The GFP insertion site was designed such that the native signal sequence and signal cleavage site of TXNDC15 would remain intact.

To create a double tagged GFP-TXNDC15, MARCHF6-ALFA cell line, polyclonal HEK293T cells tagged with GFP-TXNDC15 were used and a TEV-ALFA-2A-BFP tag was inserted after MARCHF6s. A monoclonal cell line was sorted based on GFP and BFP fluorescence.

Stable knockdowns of TXNDC15 and MARCHF6 were obtained by transfecting HEK293T cells with pX459 encoding the respective sgRNA using TransIT-293 transfection reagent using the manufacturer’s protocol (2705, Mirus, USA). The following sgRNAs were cloned into pX459: TXNDC15 (CGGGCATTTCCAGCTCTTCA), MARCHF6 (CCATGGACTATCTGCTCTCT).

GFP positive single cells of either the stable knockdown or fluorescently tagged knockin were sorted into 96-well plates using a SONY SH800S cell sorter (Sony Biotechnology, USA) 72 hours post transfection.

Depletion of TXNDC15 and MARCHF6 were verified with qPCR as described below (Figures S1F, S3H). Endogenous tagging of TXNDC15 was verified by western blotting.

In NIH 3T3, a stable knockdown cell line was created by transducing NIH 3T3 cells expressing dCas9 with either non-targeting (CGAGGTATTCGGCTCCGCG) or TXNDC15 (GCAGAAAACCGGCACCA) guides in cloned in the pMCB306 backbone. After selection with puromycin (incubation with 3 µg/mL puromycin [A1113803, Thermo Scientific, USA] for three days), knockdown was verified by qPCR (Figure S7A) and polyclonal cell lines were maintained.

### Expi293 stable expression cell line generation

Expi293 cells were transduced by adding 2.5 mL of lentivirus of corresponding construct to 10 million cells along with 8 µg/mL final concentration of polybrene in a volume of 30 mL in a 125-mL cell culture flask. After 8-12 hours, the media was exchanged to remove lentiviral particles and prevent cell clumping. Cells were maintained until they were sorted for GFP positive cells using the SONY SH800S cell sorter (Sony Biotechnology, USA).

The MARCHF6(C9A)-GFP cell line was generated by transducing wt Expi293 cells with MARCHF6(C9A)-GFP-2A-RFP lentivirus and sorting for GFP positive cells.

The MARCHF6(C9A)-GFP TXNDC15-ALFA Expi293 cell line was generated by transducing the sorted MARCHF6(C9A)-GFP with TXNDC15-ALFA-2A-BFP lentivirus and sorting for GFP and BFP positive cells.

### CRISPRi single sgRNA screen and dual genetic modifier screen

*CRISPRi single sgRNA screen:* The CRISPRi screen to identify factors for the recognition and degradation of GET1 and TMEM231 was performed as previously described.^33,34^ In short, the TOP5 CRISPRi v2 library was transduced at a multiplicity of infection 0.3 into 330 million K562-CRISPRi-Tet-ON cells stably expressing the GET1-GFP-2A-RFP reporter under an inducible promoter. Transduced cells were enriched using puromycin treatment (addition of 1 µg/mL puromycin for three consecutive days). Cells were maintained at a density of 0.5 × 10^6^ cells/mL throughout the course of the screen. On day 8, the reporter was induced with doxycycline (100-1000 ng/mL) for 36 hours and sorted on a SONY SH800S cell sorter (Sony Biotechnology Inc., USA).

During sorting, cells were first gated based on BFP expression, as a proxy for the expression of an sgRNA, followed by gating for RFP and GFP to select cells expressing the GET1 reporter. From this sgRNA and reporter positive population, cells with perturbed GFP:RFP fluorescence were sorted based on their GFP:RFP ratio. Approximately 25 million cells with GFP:RFP ratios in the highest and lowest 30% were collected, pelleted, and flash-frozen. Genomic DNA was extracted and purified using a Nucleospin Blood XL kit (740950.10, Takara Bio, JPN). Guides were amplified and barcoded by PCR using NEB Next Ultra ii Q5 MM (M0544L). The DNA library (279 bp for single guide library, 349 bp for dual guide libraries) was purified using SPRISelect beads (B23317, Beckman Coulter, USA), and purified DNA was analyzed on an Agilent 2100 Bioanalyzer prior to sequencing using an Illumina HiSeq2500 using the standard CRISPRi-v2 library sequencing primer (5’-GTGTGTTTTGAGACTATAAGTATCCCTTGGAGAACCACCTTGTTG). The results of the screen were analyzed using published protocols (https://github.com/mhorlbeck/ScreenProcessing).^34^ To ensure sufficient coverage of the guide libraries, only guides with more than 50 counts were assessed and phenotype scores were computed based on the 2 (GET1) or 3 (TMEM231) sgRNAs displaying the strongest phenotypes. The Mann-Whitney p-value was calculated using the 5 sgRNAs targeting the same gene compared to the negative controls.

#### Cloning the TXNDC15 dual-guide CRISPRi library

Cloning of the genome-wide CRISPRi dual library with a guide targeting TXNDC15 in the anchor position (see Figure S2A) was performed as previously described ^43^. Briefly, the TXNDC15 guide (5’-GACGCGGGGCGGTCGTCGAC-3’) was cloned into a hU6-CR3 cassette (pJR152, Addgene # 196280). The anchor guide with custom overhangs (see table below) was ligated into the pJR152 vector digested with BstXI/BlpI. This vector containing the hU6-TXNDC15 sgRNA-CR3 element was digested with BamHI and NotI to yield a 400 bp fragment that was gel purified. The CRISPRi-v2 library (Addgene Pooled Libraries # 83969) was digested with BamHI and NotI in the presence of shrimp alkaline phosphatase (rSAP) followed by gel purification to yield the 8,800 bp vector. This vector was ligated with the hU6-TXNDC15 sgRNA-CR3 element using T4 (NEB # M0202M) at an insert to vector ratio 1:2 for 16 hours at 16°C. This construction yielded the mU6-CR1-hU6-CR3 vector, which was purified with SPRISelect beads (Beckman Coulter B23317) and electroporated into MegaX cells at scale (ThermoFisher # C640003). The cells were incubated in 200 mL LB supplemented with 100 µg/mL carbenicillin overnight. The resulting TXNDC15 dual plasmid library was amplified and barcoded for deep sequencing (NGS) by PCR using NEB Next Ultra ii Q5 MM (M0544L) and a reverse primer (CAAGCAGAAGACGGCATACGAGATggaatcatgggaaataggccctc) and index primer. The PCR libraries were purified with SPRISelect beads to yield dual the TXNDC15 dual DNA library (349 bp), which was sequenced using an Illumina HiSeq2500 with the sequencing primer listed previously. The non-targeting (NT) dual guide genome-wide CRISPRi library was described previously (cite paper) (Addgene Pooled Library # 197348). The TXNDC15 dual guide library read counts were compared to the NT dual guide library, which showed consistent guide coverage and distribution between the libraries.

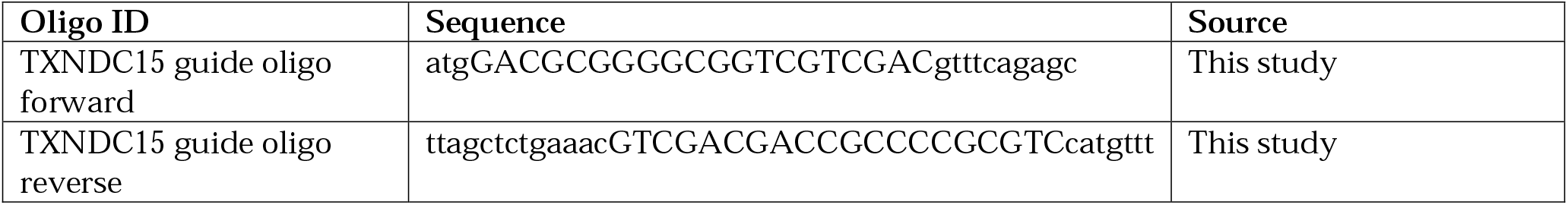

#### Dual guide genetic modifier screen

The genetic modifier screen was conducted in a similar manner, using a non-targeting dual library or TXNDC15 dual library, as previously described.^43^ The cells were sorted on a FACSAria Fusion Cell Sorter (BD Biosciences). Following isolation of cells displaying perturbed GET1-GFP:RFP ratios, amplification of the guide region was performed using a unique forward index primer in combination with a reverse primer complementary to the hU6 region upstream of the fixed guide in the dual guide vector (5’-CAAGCAGAAGACGGCATACGAGATGGAATCATGGGAAATAGGCCCTC). Sequencing and analysis were performed using the same strategy as for the single sgRNA screen.

### Ratiometric fluorescent reporter assays to test individual depletion of genes and TXNDC15 rescue

Throughout the manuscript we rely on depletion of individual genes to assess the effect of loss of specific factors on a fluorescent reporter. These reporters serve two purposes: First, because the reporters express a substrate (e.g. GET1-GFP) and a normalization marker (e.g. RFP) from a single open reading frame separated by a viral 2A ribosomal skipping site, changes in GFP:RFP ratio necessarily reflect changes that occur post-translationally. Second, because the reporter is acutely expressed, changes to the GFP:RFP ratio reflect effects to a newly synthesized protein. In both cases, these features are particularly useful for studying the function of quality control factors that must act post-translationally on nascent proteins.

#### K562 cells

To perform reporter assays in K562 cells, we transduced lentiviral vectors expressing programmed guides into K562 dCas9BFP-KRAB cells. All spinfections throughout the manuscript were performed as follows. Briefly, 250,000 cells were mixed with 200 µL of lentivirus and RPMI medium in the presence of 8 µg/mL polybrene (TR-1003, Sigma-Aldrich, USA) in a total volume of 1 mL. K562 cells in 12-well plates were spinfected at 1,000 xg for 1.5 hours at 30°C, resuspended, and cultured in 12-well or 6-well plates. Approximately 48 hours after spinfection, cells containing guides were selected by incubation with 3 µg/mL puromycin for 3 days (treated once). Cells recovered from puromycin treatment for 1.5 days and then were transduced with reporter lentivirus. 48 hours after reporter transfection, the cells were analyzed using flow cytometry, as described below.

#### HEK293T cells

For reporter assays in HEK293T cells, 0.3 x 10^6^ of wt or TXNDC15 stable knockdown or MARCHF6 stable knockdown cells were seeded into each 6-well plate. 24 hours later, cells were transduced with 300 μL lentivirus of TXNDC15 rescue construct and 8 μg/mL final concentration of polybrene. 24 hours later, the media was exchanged to remove excess polybrene. 48 hours after reporter transduction, the cells were harvested, washed, pelleted, and resuspended in 500 μL 1x Dulbecco’s Phosphate Buffered Saline (1x PBS, Thermo Scientific, USA), and analyzed by flow cytometry.

For rescue assays in HEK293T cells, 0.3 x 10^6^ of WT or TXNDC15 depleted or MARCHF6 depleted cells were seeded into each 6-well plate. 24 hours later, cells were transduced with 300 μL lentivirus of each TXNDC15 rescue construct and 8 μg/mL final concentration of polybrene. 24 hours later, the media was exchanged to remove excess polybrene, and cells were transduced with 150 μL reporter lentivirus the in presence of 8 μg/mL final concentration of polybrene. 24 hours later, cells were split 1:2 (half of the sample is used for flow cytometry and the other half is used for western blotting). 24 hours later (72 hours after rescue construct transduction and 48 hours after reporter transduction), cells were harvested, pelleted, and resuspended in 500 μL 1x PBS and analyzed by flow cytometry or washed, pelleted, and frozen for analysis via western blot.

#### RPE1 cells

For CRISPRi knockdown reporter assay experiment in RPE1, cells were transduced with sgRNA/dual guide lentiviral vectors. 6 days after guide transduction and selection for cells with guides by puromycin treatment (analogous to K562 cell treatment), cells were transduced with reporter lentiviral vectors. After 48 hours, cells were harvested, pelleted, and resuspended in 500 µL 1x PBS or DMEM/F-12 with 10% FBS and 2 mM glutamine and analyzed by flow cytometry. For rescue assays in RPE1, the experiment was performed similarly, except for the addition of rescue lentiviral vectors to the cells 72 hours before analysis as described below. Knockdown of TXNDC15 in RPE1 cells was verified using qPCR, as described below.

#### NIH 3T3 cells

Rescue assays in NIH 3T3 cells were performed using polyclonal TXNDC15 stable knockdown cell lines. 48 hours after transduction with lentiviral reporter constructs, cells were harvested and analyzed by flow cytometry as described below.

### Flow cytometry to analyze fluorescence reporter assays

HEK293T, RPE1, and NIH 3T3 cells were treated with trypsin, pelleted, and resuspended in 1x PBS for flow cytometry analysis. K562 cells were analyzed directly in RPMI media. Cells were analyzed using an Attune NxT Flow Cytometer (Thermo Scientific, USA).

Flow cytometry data was analyzed using FlowJo v10.8 Software (BD Life Sciences, USA) or by Python using the FlowCytometryTools package.^76^ For reporter assays, cells were gated based on RFP expression for cells expressing the reporter constructs. For TXNDC15 rescue assays, cells were first gated based on BFP expression and then RFP expression for cells expressing both the rescue constructs and the reporter. The relative change (kd/wt) of each reporter was calculated by taking the ratio of the median GFP:RFP intensities between that of the respective knockdown cells and the wildtype/non-targeting control cells.

Relative change (kd/wt) is either displayed in bar plots or histograms. Normalized histograms of these assays are generated by setting the median GFP:RFP ratio (or RFP:GFP ratio in the case of VAMP2) of wildtype or non-targeting control to 1. Histograms, charts, and plots were generated using Python and GraphPad Prism v.10.6.1.

### Calculating % rescue of TXNDC15 rescue constructs

To calculate how well a particular mutant of TXNDC15 can rescue GET1 stability in HEK293T TXNDC15 stable kd compared to wt cells, the following formula (1) was used:

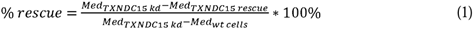

“Med” represents the median GFP:RFP value of the GET1 reporter, determined from the ratio of GFP to RFP fluorescence measurements obtained from flow cytometry.

Additionally, the relative rescue efficiency was calculated to compare rescue efficiencies among different reporters:

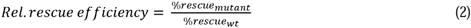

### K562 treatment with quality control factor inhibitors

To test the effect of inhibition of the VCP AAA ATPase or E1 ubiquitin-activating enzymes on TXNDC15 dependent degradation, K562 wt and TXNDC15 kd cells were prepared as described above. 48 hours after transduction with the GET1-GFP-2A-RFP reporter construct, cells were treated with either DMSO control, or 250 nM E1 inhibitor MLN7243 (S8341, Selleckchem, USA) in DMSO, or 1.25 µM VCP inhibitor CB-5083 (S8101, Selleckchem, USA) in DMSO for 6 hours. Afterwards, cells were analyzed by flow cytometry as described above.

### Mass spectrometry sample preparation of TXNDC15 and MARCHF6 immunoprecipitations (IP-MS)

Native immunoprecipitation of endogenously expressed GFP-TXNDC15 from HEK293T cells or exogenously expressed MARCHF6-GFP from Expi293 cells were performed as described previously.^77^ The control sample for background binding of TXNDC15 immunoprecipitation in Figure 2B was wt HEK293T cells that were not endogenously tagged with GFP (Figure S3D). The control sample for background binding of MARCHF6 immunoprecipitation in Figure 2C was the same Expi293 cells overexpressing MARCHF6(C9A)-GFP but the immunoprecipitation was carried out as described below without addition of GFP-nanobody (Figure S3E). Briefly, cells were incubated in Solubilization Buffer (50 mM HEPES, pH 7.5, 200 mM NaCl, 2 mM MgOAc_2_, 1% [w/v] glycol-diosgenin [GDN, GDN101, Anatrace, USA], 1x Roche cOmplete EDTA-free protease inhibitor cocktail [COEDTAF-RO, Sigma-Aldrich, USA], 1 mM DTT) for 1 hour on ice. Solubilized lysate was clarified by centrifugation at 34,540 xg in an SS-34 rotor in a Sorvall RC6+ Superspeed centrifuge for 45 min at 4°C. At the same time, biotinylated His14-Avi-SUMOEu1-tagged anti-GFP nanobody was incubated with Pierce™ Streptavidin Magnetic Beads (88817, Thermo Scientific, USA) head-over-tail in Wash Buffer (50 mM HEPES, pH 7.5, 200 mM NaCl, 2 mM MgOAc_2_, 1 mM DTT, 0.0053% GDN) for 30 min at 4°C. Beads were washed and incubated with 50 mM HEPES pH 7.5 containing 100 µM biotin on ice for 5 min to occupy unbound streptavidin sites on the magnetic beads. Then, beads were washed in Solubilization Buffer twice before being incubated with clarified lysate head-over-tail for 1-1.5 hours at 4°C. After incubation, beads were washed 3 times in Wash Buffer. Proteins were eluted by addition of 0.5 µM SUMOEu1 protease (Addgene # 149333) and incubated for 30-45 min at 4°C with rotating. Elution protein samples were precipitated by adding 1:10 volume of 100% trichloroacetic acid (TCA), followed by a 10 min incubation on ice before being pelleted at maximum speed in a benchtop centrifuge at 4°C. Protein pellets were washed with ice-cold acetone twice before being air dried at RT. TCA-precipitated protein pellets were dissolved in 8 M urea prepared in 50 mM HEPES, pH 8.0. Samples were reduced by incubating with 4 mM Tris(2-carboxyethyl)phosphine hydrochloride (TCEP) (20490, Thermo Scientific, USA) for 20 min at 37°C, alkylated by incubating with 12 mM 2-chloro-acetamide (CAA) (ICN15495580, MP Biomedicals) for 15 min at 37°C.

Samples were digested with 2 ng/µL Lysyl Endoproteinase (Lys-C) (125-05061, Wako Chemicals) for 4 hours at 37°C. Then, samples were diluted 4-fold so that final concentration of urea reached 2 M, and digested overnight with 0.6 ng/µL Trypsin (90057, Thermo Scientific, USA). Samples were desalted using the Pierce™ C18 Spin Columns (89870, Thermo Scientific, USA) as per manufacturer’s instructions. After being eluted off the desalting columns, the samples were lyophilized before mass spectrometry analysis.

### Mass spectrometry analysis for TXNDC15 and MARCHF6 interaction partners

LC-MS/MS analysis was performed with an EASY-nLC 1200 (Thermo Scientific, USA) coupled to a Q Exactive HF hybrid quadrupole-Orbitrap mass spectrometer (Thermo Scientific, USA). Peptides were separated on an Aurora UHPLC Column (25 cm × 75 μm, 1.7 μm C18, AUR3-25075C18, Ion Opticks, AUS) with a flow rate of 0.35 μL/min for a total duration of 75 min and ionized at 1.6 kV in the positive ion mode. The gradient was composed of 6% solvent B (3.5 min), 6-25% B (42 min), 25-40% B (14.5 min), and 40–98% B (2 min) and 98% B (13 min); solvent A: 2% acetonitrile (ACN; A9554, Fisher Scientific, USA) and 0.2% formic acid (FA, A11750, Fisher Scientific) in LC-MS grade water (W6212, Fisher Scientific, USA); solvent B: 80% ACN and 0.2% FA. MS1 scans were acquired at the resolution of 60,000 from 375 to 1600 m/z, AGC target 3e6, and maximum injection time 15 ms. The 12 most abundant ions in MS1 scans are selected for fragmentation via higher-energy collisional dissociation (HCD) with a normalized collision energy (NCE) of 28. MS2 scans were acquired at a resolution of 30,000, AGC target 1e5 and a maximum injection time 45 ms. Dynamic exclusion was set to 45 s and ions with charge +1, +6, +8 and >+8 were excluded. The temperature of ion transfer tube was 275°C and the S-lens RF level was set to 55. RAW files were searched with Proteome Discoverer SEQUEST (version 2.5, Thermo Scientific, USA) against in silico tryptic digested the UniProt Human proteome Swiss-Prot database (UP000005640). The maximum missed cleavages were set to 2. Dynamic modifications were set to oxidation on methionine (M, +15.995 Da), deamidation on asparagine and glutamine (N and Q, +0.984 Da) and protein N-terminal acetylation (+42.011 Da). Carbamidomethylation on cysteine residues (C, +57.021 Da) was set as a fixed modification. The maximum parental mass error was set to 10 ppm, and the MS2 mass tolerance was set to 0.03 Da. The maximum false peptide discovery rate was specified as 0.01 using the Percolator Node validated by q-value. Depending on the IP sample, the average abundance over 1 or 2 replicates was calculated and protein hits were ranked in ascending order. The average abundance was normalized to the most abundant protein in the sample.

### Whole cell proteomics

#### Sample preparation

K562 dCas9BFP-KRAB cells were transduced with non-targeting, TXNDC15 kd or MARCHF6 kd sgRNA and selected by puromycin treatment as described above. After an 8-day knockdown, four technical replicates of 750,000 cells each were harvested per sample. Samples were prepared according to the EasyPep Mini MS Sample Prep Kit (Thermo Scientific). Cell pellets were resuspended in lysis buffer and sonicated for 3 s. 100 μg of protein were further reduced and alkylated at 95°C followed by overnight digestion with Trypsin and Lys-C at 4°C. Using the Peptide Clean-up column, peptides were bound, washed, and eluted. After drying the eluate using a vacuum centrifuge at room temperature, samples were resuspended in 2% acetonitrile and 0.2% formic acid.

#### Sample processing for replicate 1

LC-MS/MS analysis was performed on an Orbitrap Eclipse Tribrid mass spectrometer (Thermo Fisher Scientific, USA) coupled to a Vanquish Neo UHPLC system (Thermo Fisher Scientific, USA). Peptides were separated on an analytical Aurora Frontier™ column (60 cm × 75 μm, 1.7 μm C18, AUR3-60075C18-CSI, Ion Opticks) with a flow rate of 0.3 μL/min for a total duration of 130 min and ionized at 1.6 kV in the positive ion mode. The gradient was composed of 6% solvent B (7 min), 6-25% B (88 min), 25-40% B (25 min), 40–98% B (1 min) and 98% B (9min); solvent A: 2% ACN and 0.2% FA in water; solvent B: 80% ACN and 0.2% FA. The gradient was composed of 6% solvent B (8.5 min), 6-25% B (90.5 min), 25-40% B (25 min), and 40–98% B (16 min); solvent A: 0.2% formic acid in water; solvent B: 80% ACN and 0.2% formic acid. MS1 scans were acquired at the resolution of 120,000 from 375 to 2,000 m/z, AGC target 1e6, and maximum injection time 50 ms. MS2 scans were acquired in the ion trap using fast scan rate on precursors with 2-7 charge states and quadrupole isolation mode (isolation window: 1.2 m/z) with higher-energy collisional dissociation (HCD, 30%) activation type. Dynamic exclusion was set to 30 s. The temperature of ion transfer tube was 300°C and the S-lens RF level was set to 30.

RAW files were analyzed using Proteome Discoverer (PD) (version 3.0, Thermo Fisher Scientific) based on the CHIMERYS algorithm with Inferys 2.1. fragmentation prediction against the UniProt human proteome database (UP000005640, 2022 version, 79,740 entries). The maximum missed cleavages was set to 2. Dynamic modifications were set to oxidation on methionine (M, +15.995 Da) and carbamidomethylation on cysteine residues (C, +57.021 Da) was set as a fixed modification. The maximum parental mass error was set to 10 ppm, and the MS2 mass tolerance was set to 0.3 Da. The false discovery threshold was set strictly to 0.01 using the Percolator Node validated by q-value. The relative abundance of parental peptides was calculated by integration of the area under the curve of the MS1 peaks using the Minora LFQ node.

#### Sample processing for replicate 2

LC-MS/MS data were acquired in data-independent acquisition (DIA) mode using a Thermo Fisher Scientific Vanquish Neo UHPLC system coupled to an Orbitrap Astral mass spectrometer. A total of 0.5 µg peptides was injected onto a PepMap Neo Trap Cartridge (5 cm x 300 µm,5 µm C18, ThermoFisher Scientific) before refocusing on an Aurora UHPLC column (25 cm x 75 µm, 1.7 µm C18, AUR3-25075C18-XT, Ion Opticks) with a flow rate of 0.25 µL/min in a 33.5-min gradient. The gradient was composed of 5-12% B (0.9 min, 0.65uL/min), 12.3% B (0.1 min, 0.25uL/min), 12.3-30% B (24 min, 0.25uL/min), 30–44% B (2.5 min,0.25uL/min), 44-99% B (1 min,0.25uL/min) and 99% B (5 min, 0.5uL/min); mobile phase A: 0.1% FA in water; mobile phase B: 80% ACN and 0.1% FA. The Orbitrap Astral MS was operated at a full MS resolution of 240,000 with a full scan range of 380–980 m/z. The full MS AGC was set to 500%, IT was set to 5 ms and RF lens was set to 40. MS/MS scans were recorded with a 2 Th isolation window and 3.5 ms maximum ion injection time. The normalized AGC target was set to 500% and the isolated ions were fragmented using HCD with 25% NCE.

RAW files were processed using the CHIMERYS intelligent search algorithm with Inferys 4.7.0 fragmentation prediction (version 4.0, MSAID GmbH) in the Proteome Discoverer (version 3.2, Thermo Scientific) against the UniProt human proteome Swiss-Prot database (UP000005640, 2023 version, 20,411 entries). The maximum missed cleavages was set to 2. Dynamic modifications were set to oxidation on methionine (M). Carbamidomethylation on cysteine residues (C) was set as a fixed modification. Peptides between 7 and 30 amino acids were considered and the false discovery rate (FDR) for peptide spectra matches (PSMs) identifications was set to 1%. Quantification type was set to MS2 Apex.

#### Post-analysis

Differential expression analysis was performed using the tidyproteomics package v1.7.3 ^78^. Statistically significant differences between groups were calculated using two-sided Student t-test. Volcano and heatmap plots were generated using the ggplot2 package v.3.4.4.^79^ The mass spectrometry proteomics data have been deposited to the ProteomeXchange Consortium via the PRIDE partner repository with the dataset identifier PXD063158.^80^

#### Comparison of replicates

Proteins were filtered to identify only those that showed significant changes in abundance ratio between wt and kd cells, displaying a log2 abundance ratio >0.5 or <-0.5 and more than 1 detected peptide. This pool of hits was then compared between the two replicates and only proteins that appeared in both replicates and showed changes in the same direction between replicates were included in the final Venn diagram.

### Expression and purification of a recombinant TXNDC15-MARCHF6 complex

To determine if MARCHF6 and TXNDC15 interacted directly, we expressed and purified a recombinant complex. Expi293 cells stably expressing MARCHF6(C9A)-GFP and TXNDC15-ALFA were subjected to first an anti-ALFA immunoprecipitation and then anti-GFP immunoprecipitation. Briefly, 500 mL of cells were harvested at 4500 rpm and washed with ice-cold 1x PBS. Cell pellets were resuspended in solubilization buffer (50 mM HEPES pH 7.5, 2 mM MgOAc_2_, 200 mM KOAc, 1% [w/v] GDN, 1x Roche cOmplete EDTA-free protease inhibitor cocktail [COEDTAF-RO, Sigma-Aldrich, USA], 1 mM DTT) and incubated at 4°C for 1 hour. Solubilized lysate was clarified by centrifugation at 34,540 xg in an SS-34 rotor in a Sorvall RC6+ Superspeed centrifuge for 45 at 4°C. Simultaneously, biotinylated His14-Avi-SUMOStar-tagged anti-ALFA nanobody was incubated with Pierce™ Streptavidin Magnetic Beads (88817, Thermo Scientific, USA) head-over-tail in Wash Buffer (50 mM HEPES, pH 7.5, 200 mM NaCl, 2 mM MgOAc_2_, 1 mM DTT, 0.0053% GDN) for 30 min at 4°C. After this immobilization, beads were washed and incubated with 50 mM HEPES pH 7.5 containing 100 µM biotin on ice for 5 min to occupy unbound biotin binding sites on the magnetic beads.

Then, beads were washed in Solubilization Buffer twice before being incubated with clarified lysate head-over-tail for 1.5 hours at 4°C. After incubation, beads were washed 3 times in Wash Buffer. Proteins were eluted by addition of 0.3 µM SUMOStar protease and incubated for 45 min at 4°C with rotating. Eluted proteins were then subjected to anti-GFP immunoprecipitation as described above. Finally, eluted proteins were analyzed using SDS-PAGE.

### Preparation of human ER microsomes

Endogenously tagged GFP-TXNDC15 and MARCHF6-ALFA HEK293T cells were transduced with non-targeting or TXNDC15 kd shRNA and selected by puromycin treatment at 1 μg/mL for 2 days. Cells were adapted to suspension, harvested and washed in 1x PBS. Cell pellets resuspended in 4 times the pellet volume of sucrose buffer (10 mM HEPES, pH 7.5, 250 mM sucrose, 2 mM, 1x Roche cOmplete EDTA-free protease inhibitor cocktail [COEDTAF-RO, Sigma-Aldrich, USA]) and lysed by douncing at 4°C. Cells were diluted to twice the lysis volume in sucrose buffer and pelleted at 3,214 xg for 35 min at 4°C. Supernatant from the cell lysate was transferred to a new tube and pelleted again at 3,214 xg for 35 min at 4°C. Samples were pelleted in an ultracentrifuge in an MLA-80 rotor (Beckman-Coulter, USA) at 75,000 xg for 1 hour at 4°C to isolate the microsomal fraction. The microsomal pellet was resuspended in microsome buffer (10 mM HEPES, pH 7.5, 250 mM sucrose, 1 mM MgOAc_2_, 0.5 mM DTT) to an A280 of 75. To remove contaminating RNAs from human rough microsomes (hRMs), hRMs were mixed with CaCl_2_ (1 mM) and micrococcal nuclease (0.125 U/μL) on ice before incubating for 6 min at 25°C. Nucleased hRMs were transferred to ice and immediately quenched with EGTA (2 mM). Nucleased hRMs were flash frozen and stored at -80°C prior to use in in vitro translations. Only aliquots of nucleased hRMs with one freeze-thaw cycle were used in experiments.

### Mammalian in vitro translations

Translation reactions were prepared using nucleased RRL supplemented with hRMs, as described above and previously.^81,82^ DNA templates for in vitro transcription were made by PCR using primers within the SP6 promoter (5’ end) and after the stop codon (3’ end). Transcription reactions were carried out by combining 4.8 µL T1 mix^81^, 0.1 µL RNAsin (N251, Promega, USA), 0.1 µL SP6 polymerase (M0207L, New England Biolabs, USA) and 50 ng PCR product. Transcription reactions were incubated at 37°C for 2 hours, and then used directly in a translation reaction, which was incubated for 45 min at 32°C. Radioactive ^35^S-methionine (NEG709A005MC, Perkin Elmer, USA) was used to label nascently translated proteins, unless otherwise indicated. Reactions were analyzed on SDS-PAGE followed by autoradiography. In vitro translation reactions for mass spectrometry were treated with 1 mM puromycin for 10 min at 32 C immediately following translation in order to fully release nascent proteins from the ribosome.

#### Analysis of MARCHF6 interaction with orphan GET1 and TMEM231 in presence or absence of TXNDC15

Translation reactions were performed as described above in the presence of hRMs prepared from endogenously tagged GFP-TXNDC15 MARCHF6-ALFA HEK293T cells subjected to non-targeting or TXNDC15 kd shRNA for 8 days. Translation reactions were pelleted through a sucrose cushion (50 mM HEPES, pH 7.5, 2 mM MgOAc_2_, 100mM KOAc, 20% sucrose [w/v]) for 20 min at 55,000 RPM at 4°C in a TLA-55 rotor (Beckman-Coulter). Membrane fractions were resuspended in solubilization buffer (50 mM HEPES pH 7.5, 2 mM MgOAc_2_, 200 mM KOAc, 1% [w/v] GDN) and incubated on ice for 20 min. Solubilized membrane was pelleted at 14,000 xg for 10 min in a table-top centrifuge for 10 min at 4°C and the clarified solubilized membrane was transferred to a fresh tube. Anti-ALFA nanobody was immobilized onto magnetic streptavidin beads as described above. Beads were incubated with solubilized membrane for 2 hours at 4°C with rotating. Unbound fraction was removed by pipetting and beads were washed 3 times before being incubated with wash buffer containing 0.25 µM SUMOstar protease for 30 min at 4°C. Input and elution samples were analyzed on SDS-PAGE followed by autoradiography or western blotting. The relative change between nt and TXNDC15 kd samples was calculated by quantification of the relative change in band intensity between input samples containing nt (I_nt, input_) or TXNDC15 kd (_kd, input_) membranes. Elution samples were normalized to contain the same amount of MARCHF6 by western blot and the relative change in band intensities was calculated between nt (I_nt, IP_) or TXNDC15 kd (I_kd, IP_) membranes. Individual replicates were normalized on the change detected in the input sample and mean and standard deviations were calculated. Statistical significance was determined by a One-sample t-test to test whether the mean of the relative change between nt and TXNDC15 kd samples equals 1.

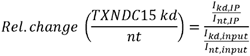

#### Crosslinking of MARCHF6 with GET1 or TMEM231

GET1 or TMEM231 were synthesized using the rabbit reticulocyte in vitro translation system in the presence of hRMs as described above. The amounts of added in vitro transcription product were titrated to achieve similar amounts of translation product. The hRMs were obtained from endogenously labeled GFP-TXNDC15, MARCHF6-ALFA HEK293T cells that were treated with nt or TXNDC15 kd shRNA for 8 days. To remove translation products that were not inserted, microsomes were isolated by pelleting through a sucrose cushion (50 mM HEPES, pH 7.5, 2 mM MgOAc_2_, 100mM KOAc, 20% sucrose [w/v]) for 30 min at 55,000 RPM at 4°C (TLA-55, Beckman-Coulter). Pellets were resuspended in resuspension buffer (50 mM HEPES, 2 mM MgOAc_2_, 150 mM KOAc) and incubated with 250 uM dithiobis(succinimidyl propionate) (DSP, Thermo Scientific) for 30 min on ice. As a control, the equivalent volume of DMSO was added. The reaction was quenched with 20 mM Tris/HCl at pH 7.5 and membranes were solubilized under denaturing conditions by addition of 1% SDS in 0.1 M Tris/HCl at pH 8. Samples were diluted 10-fold in IP buffer (50 mM HEPES, 300 mM NaCl, 0.5% Triton X-100) and incubated with Anti-ALFA-nanobody immobilized onto magnetic streptavidin beads for 2 hours at 4°C. After washing with wash buffer (50 mM HEPES, 300 mM NaCl, 0.05% Triton X-100), elution was performed using 1 μM SUMOstar protease and samples were analyzed by SDS-PAGE followed by autoradiography or western blotting. If indicated, western blot samples were treated with 100 mM DTT for 15 min at room temperature to reduce the cysteines of DSP and cleave the crosslink. Quantification was performed as described above. For quantification of the crosslinked sample, the crosslinking band including the intensity above was analyzed.

### Ubiquitination assay in human cells

K562 cells lines stably expressing inducible GET1-GFP or TMEM231-GFP reporters were spinfected with non-targeting or TXNDC15 sgRNA as described above. 48 hours after transduction, cells were selected with 3 μg/mL puromycin and then incubated for a further four days. Cells were then induced by addition of 0.5 μg/mL doxycycline and, 24 hours later, treated with 10 μM proteasome inhibitor MG132 (10012628, Cayman Chemical) or DMSO for 6 hours. Cells were harvested and washed with PBS. The pellet was resuspended in solubilization buffer (50 mM HEPES pH 7.5, 200 mM NaCl, 2 mM MgOAc_2_, 1 mM DTT, 1 % GDN, protease inhibitor tablet (Roche)) and incubated for 30 min on ice. Cell debris was pelleted at 18,000 xg for 15 min at 4°C and the supernatant was normalized based on the A280 absorbance using IP buffer (50 mM HEPES, 200 mM NaCl, 2 mM MgOAc_2_, 1 mM DTT, 1 % GDN). Pierce™ magnetic streptavidin beads were equilibrated in IP buffer and loaded with anti-GFP nanobody as described above. Normalized samples were incubated at 4°C for 1 hour. After washing the beads with wash buffer (50 mM HEPES, 200 mM NaCl, 2 mM MgOAc_2_, 1 mM DTT, 0.53 % GDN), samples were eluted with 0.3 μM SUMOEu1 protease and analyzed in triplicate via SDS-PAGE and western blotting.

### Retrotranslocation assay in human cells

K562 cells lines stably expressing inducible GET1-GFP or TMEM231-GFP reporters were spinfected with non-targeting or TXNDC15 sgRNA and treated with MG132 proteasome inhibitor as described above. The pellet was resuspended in lysis buffer (50 mM HEPES, 10 mM KOAc, 2 mM MgOAc_2_, 1 mM DTT, 50 mM digitonin, protease inhibitor tablet (Roche)) and swollen by addition of 12% final volume of water for 10 min at 4°C. Cells were lysed by passage through a 27 ¼ gauge needle 12 times and 0.19 M NaCl was added. The soluble cytosol fraction was separated by centrifugation at 18,000 xg for 15 min at 4°C and the membrane pellet was washed with lysis buffer. It was further resuspended in IP buffer (50 mM HEPES, 200 mM NaCl, 2 mM MgOAc_2_, 1 mM DTT, 1 % GDN), incubated at 4°C for 30 min and centrifuged at 18,000 xg for 15 min at 4°C. The remaining supernatant suspension represents the membrane fraction. Cytosol and membrane samples were normalized based on A280 absorbance within the same compartment and subjected to an anti-GFP immunoprecipitation. Pierce™ magnetic streptavidin beads were equilibrated in IP buffer and loaded with anti-GFP nanobody as described above. Normalized samples were incubated at 4°C for 1 hour. After washing the beads with wash buffer (50 mM HEPES, 200 mM NaCl, 2 mM MgOAc_2_, 1 mM DTT, 0.53 % GDN), samples were eluted with 0.3 μM SUMOEu1 protease and analyzed in triplicate via SDS-PAGE and western blotting.

### Immunofluorescence (IF) confocal microscopy

#### Preparation and coating of microscopy cover glass

Circular cover glass (72222-01, Electron Microscopy Sciences, USA) was sterilized, placed in the bottom of each 12-well plate, and coated by incubating with 1 mL of 0.01 mg/mL poly-D-lysine hydrobromide (P6407, Sigma-Aldrich, USA) for 1 hour at RT. Coated cover glass was washed 3 times with nanopure water and dried overnight at RT.

#### Seeding conditions for endogenously tagged GFP-TXNDC15 HEK293T cells

For Figure 2A, 275,000 endogenously tagged GFP-TXNDC15 HEK293T cells were seeded into each 12-well with coated cover glass. Cells were grown on cover glass for 48 hours before preparation of imaging slides.

#### Seeding conditions for NIH 3T3 cells

For the imaging rescue assay in Figure 6B, D, and Figure S7C, 100,000 NIH 3T3 cells (wt or TXNDC15 stable kd) were seeded in 6-well plates and treated with 8 µg/mL final concentration of polybrene or transduced with the respective lentivirus and polybrene. After 48 hours, 65,000 cells were seeded into each 12-well with coated cover glass. To induce cilia formation, NIH 3T3 cells were serum starved by incubation with DMEM supplemented with 0.5% FBS, 32 mM glutamine, 4.5 g/L D-glucose and 110 mg/L sodium pyruvate for 48 hours before preparation of imaging slides.

#### Seeding conditions for RPE1 cells

For the imaging rescue assay in Figure 6D, E, RPE1 cells were transduced with sgRNA guide lentiviral vectors targeting TXNDC15 or a non-targeting control on Day 1 (24 hours after seeding). On Day 2 (24 hours after lentiviral transduction), cells were subjected to puromycin treatment (8 µg/mL final concentration) to select for cells with guides for 48 hours. On Day 7, 100,000 cells were seeded into each 6-well plate. On Day 8, 150 µL of lentivirus of TXNDC15 rescue constructs were added to corresponding 6-well plates in the presence of 8 µg/mL polybrene. On Day 10, 90,000 cells (either wildtype or from knockdown-rescue experiment) were seeded into each 12-well with coated cover glass. On Day 11, to induce cilia formation, adherent RPE1 cells were serum starved by incubation in DMEM/F12 media supplemented with 2 mM glutamine and 0% FBS for 48 hours before preparation of imaging slides on Day 13.

For the TXNDC15 and MARCHF6 knockdown imaging experiment in Figure 6F, G, RPE1 cells were transduced with dual gRNA guides targeting TXNDC15, MARCHF6, TXNDC15 and MARCHF6, or a non-targeting control on Day 1 (24 hours after seeding). For the TMEM231 knockdown imaging experiment in Figure 6J, RPE1 cells were transduced with sgRNA guide targeting TMEM231 on Day 1 (24 hours after seeding). For both experiments, on Day 2 (24 hours after lentiviral transduction), cells were subjected to puromycin treatment (8 µg/mL final concentration) to select for cells with guides for 48 hours. On Day 7, 90,000-100,000 cells were seeded into each 12-well with coated cover glass. On Day 8, to induce cilia formation, adherent RPE1 cells were serum starved by incubation in DMEM/F12 media supplemented with 2 mM glutamine and 0% FBS for 48 hours before preparation of imaging slides on Day 10.

#### Imaging slides preparation

Cells were fixed in 4% paraformaldehyde (PFA; 15714, Electron Microscopy Sciences) in 1x PBS for 15 min and permeabilized cells in 0.1% Triton X-100 in PBS for 5 min. Samples were blocked in IF Blocking Buffer (10% FBS, 5% Normal Donkey Serum [017-000-121, Jackson ImmunoResearch] in 1x PBS) for 30 min.

For subsequent incubation steps, cover glasses are inverted onto droplets on parafilm in a humidified environment to avoid evaporation. Samples were incubated in 60-µL droplets of primary antibodies diluted in IF Blocking Buffer for 1-2 hours and washed 3 times with 1x PBS. Then, samples were incubated in 60-µL droplets of secondary antibodies diluted in IF Blocking Buffer for 1 hour and washed 3 times with 1x PBS. Specifically, for imaging of the endogenously tagged HEK293T cells in Figure 2A, the samples were first stained with rabbit anti-Sec61β (1:50), washed, stained with goat anti-rabbit IgG Alexa Fluor 488 (1:500), washed, stained with anti-GFP antibody Alexa Fluor 488 (1:100), and washed. For imaging of the RPE1 cells in Figure S3C, samples were first stained with mouse anti-Arl13b (1:100), washed, stained with goat anti-mouse IgG Alexa Fluor 647 (1:500), and washed. For imaging of the NIH 3T3 cells in Figures 6B and S7A, samples were first stained with rabbit anti-Arl13b (1:100) and mouse anti-acetylated tubulin (1:100), washed, stained with goat anti-rabbit IgG Alexa Fluor 488 (1:500) and goat anti-mouse IgG Alexa Fluor 647 (1:500), and washed. For imaging of the RPE1 cells in Figures 6D, F, and S7B, samples were first stained with rabbit anti-Arl13b (1:100) and mouse anti-acetylated tubulin (1:100), washed, stained with goat anti-rabbit IgG Alexa Fluor 488 (1:500) and goat anti-mouse IgG Alexa Fluor 647 (1:500), and washed.

Samples were stained with 0.2 µg/mL DAPI (MBD0015, Sigma-Aldrich, USA) in 1x PBS for 5 min and washed 2 times with 1x PBS, followed by 2 washes with nanopure water. To mount samples onto microscope slides, each stained cover glass was inverted onto 7 µL of SlowFade™ Gold Antifade Mountant (S36936, Invitrogen, USA) was added on a glass microscope slide. Cover glasses were sealed and secured in place on microscope slide with nail polish. Prepared microscope slides were stored at RT overnight to dry and stored at 4°C until imaging.

#### Imaging conditions

Samples were imaged using the Leica STELLARIS 8 FALCON confocal fluorescence microscope (Leica Microsystems, USA) using the 63x oil-immersion objective.

#### Imaging data analysis

All imaging analyses were performed in FIJI.^83^ Cilia data analysis was done using the CiliaQ package.^84^

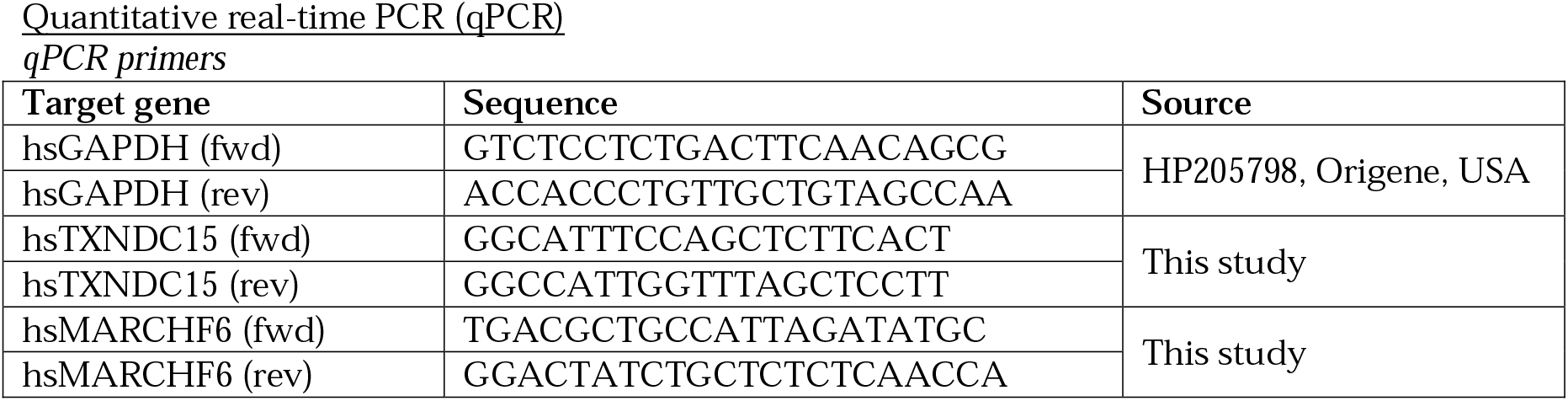
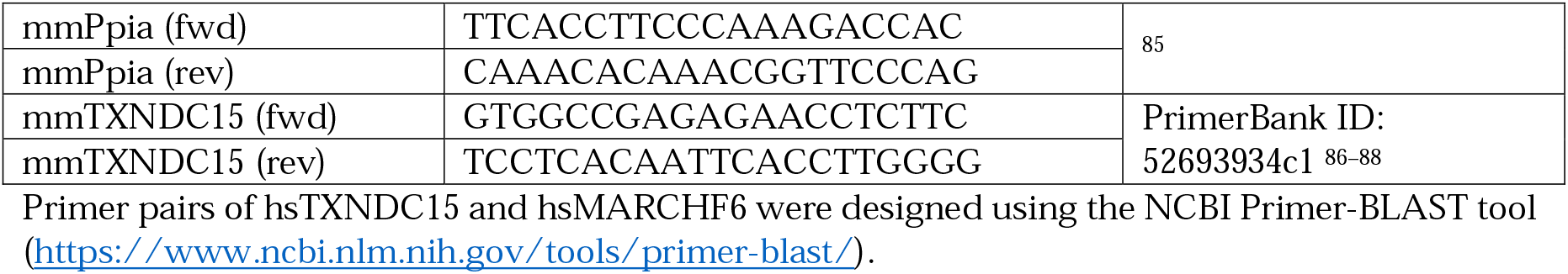

Primer pairs of hsTXNDC15 and hsMARCHF6 were designed using the NCBI Primer-BLAST tool (https://www.ncbi.nlm.nih.gov/tools/primer-blast/).

*qPCR experiment:* Cells were harvested, washed with 1x PBS, pelleted, and flash frozen. Total RNA from these cells were extracted using the Direct-zol RNA Miniprep Kit (R2050, Zymo Research, USA) and TRIzol Reagent (15596018, Invitrogen, USA). Reverse transcription of the total RNA was performed using the SuperScript™ III First-Strand Synthesis SuperMix (18080400, Invitrogen, USA). qPCR reactions were set up in triplicates using SYBR™ Green Universal Master Mix (4334973, Applied Biosystems, USA) and ran on the StepOnePlus™ Real-Time PCR System (4376600, Applied Biosystems, USA). GAPDH or Ppia were used as the housekeeping gene control and relative expression of each protein was calculated using the Pfaffl method.^89^ Results are represented as relative expression to control cells (either wt or nt kd).

### Sequence alignment

Sequence alignments were performed using Jalview 2.11.4.1.^90^ The sequences used for alignment were: hsTXNDC15 (Q96J42), hsTXNDC11 (Q6PKC3), hsTXNDC5 (Q8NBS9), hsPDIA4 (P13667).

### Other software

Flow cytometry data was analyzed using FlowJo 10.10.0.

Histograms, charts, and plots were generated using Python and GraphPad Prism v.10.6.1.

Molecular models were generated using the PyMOL Molecular Graphics System, Version 3.0 Schrödinger, LLC.

AlphaFold 3 predicted models were generated using the AlphaFold Server.^91^ Illustrations were made using Adobe Illustrator by Adobe Inc.

### QUANTIFICATION AND STATISTICAL ANALYSIS

For analysis of the CRISPRi screen, sequencing reads were aligned to the hCRISPRi-v2 compact library (5 sgRNAs per gene, Addgene pooled library # 83969) sequences using Python 3.7 as described previously.^33,34^ Only perfect matches to a guide strand sequence from our library were counted and overall alignments were generally only considered if over 70%. Generation of negative control genes and calculation of phenotype scores and Mann–Whitney p-values was performed as described previously using the ScreenProcessing software available on Github (https://github.com/mhorlbeck/ScreenProcessing).^33,34^ The sgRNA-level phenotype scores were calculated as log2 enrichment. The Mann-Whitney p-value was calculated by comparing the phenotypes of all sgRNAs targeting the gene to all negative control sgRNAs. To ensure sufficient coverage of the guide libraries, only guides with more than 50 counts were assessed and phenotype scores were computed based on the 2 (GET1) or 3 (TMEM231) sgRNAs displaying the strongest phenotypes. For GET1, 2 replicates were analyzed together.

Flow cytometry data was analyzed using FlowJo software from where the median of the distribution was determined. Three biological replicates were recorded and the mean and standard deviation were calculated where described in the figure legends.

For the in vitro translation followed by immunoprecipitation (Figure 4D) and the crosslinking experiment (Figure 4E), quantification is described in the method details. Statistical significance was calculated by a one-sample t-test using the GraphPad Prism webtools for 3 or 4 replicates as indicated in the figure legends.

For imaging (Figures 6C, E, G), 3 or 4 biological replicates were analyzed with a total number of cells as indicated in the figure legends. Statistical significance was tested by Welch’s t-test using GraphPad Prism software.

Significance levels are defined as: * p-value <0.05; ** p-value<0.01; *** p-value<0.001; **** p-value<0.0001.

## ACKNOWLEDGEMENTS

We thank all members of the Voorhees lab especially Lucas Humayun for assistance in construct generation, the Caltech Flow Cytometry facility, the Caltech Millard and Muriel Jacobs Genetics and Genomics Laboratory and the Caltech Biological Imaging Facility. This work was supported by: the Heritage Medical Research Institute (RMV), an NSF-CAREER award 2145029 (RMV), the Burrough’s Wellcome Fund (RMV), the Pew-Stewart Foundation (RMV), the Sontag Foundation (RMV), a Rosen Family fellowship (KRP), an Arie Jan Haagen-Smit Fellowship (KRP), and a Human Frontier Science Program long-term fellowship (LAKB). RMV is a Howard Hughes Medical Institute Freeman Hrabowski Scholar.

## AUTHOR CONTRIBUTIONS

Conceptualization, R.M.V., V.N.N., L.A.K.B., K.R.P.; methodology, V.N.N., L.A.K.B., K.R.P., A.G.; Investigation, V.N.N., L.A.K.B., K.R.P., J.Z, M.E, T.W., N.C., A.M., M.L.W.; writing—original draft, R.M.V.; writing—review & editing, R.M.V., V.N.N., L.A.K.B., A.M., T.W.; funding acquisition, R.M.V., L.A.K.B., K.R.P.; resources, J.W. and T.C.; supervision, R.M.V., J.W. and T.C..

## DATA AND CODE AVAILABILITY

The genome wide CRISPR screens datasets are provided as table S1, and S2. Mass spectrometry datasets are provided as table S3 and S4. The mass spectrometry proteomics data have additionally been deposited to the ProteomeXchange Consortium via the PRIDE partner repository with the dataset identifier PXD063158. A table of all oligonucleotides used is provided as table S5. All other data are available in the main text or the supplementary materials.

## REFERENCES

1. McCracken, A.A., and Brodsky, J.L. (1996). Assembly of ER-associated protein degradation in vitro: dependence on cytosol, calnexin, and ATP. Journal of Cell Biology 132, 291–298. 10.1083/jcb.132.3.291.

2. Werner, E.D., Brodsky, J.L., and McCracken, A.A. (1996). Proteasome-dependent endoplasmic reticulum-associated protein degradation: An unconventional route to a familiar fate. Proc. Natl. Acad. Sci. U.S.A. 93, 13797–13801. 10.1073/pnas.93.24.13797.

3. Neal, S., Jaeger, P.A., Duttke, S.H., Benner, C. K., Glass, C., Ideker, T., and Hampton, R.Y. (2018). The Dfm1 Derlin Is Required for ERAD Retrotranslocation of Integral Membrane Proteins. Molecular Cell 69, 306–320.e4. 10.1016/j.molcel.2017.12.012.

4. Huttlin, E.L., Bruckner, R.J., Paulo, J.A., Cannon, J.R., Ting, L., Baltier, K., Colby, G., Gebreab, F., Gygi, M.P., Parzen, H., et al. (2017). Architecture of the human interactome defines protein communities and disease networks. Nature 545, 505–509. 10.1038/nature22366.

5. Marsh, J.A., and Teichmann, S.A. (2015). Structure, Dynamics, Assembly, and Evolution of Protein Complexes. Annual Review of Biochemistry 84, 551–575. 10.1146/annurev-biochem-060614-034142.

6. Pleiner, T., Hazu, M., Tomaleri, G.P., Januszyk, K., Oania, R.S., Sweredoski, M.J., Moradian, A., Guna, A., and Voorhees, R.M. (2021). WNK1 is an assembly factor for the human ER membrane protein complex. Molecular Cell 81, 2693–2704.e12. 10.1016/j.molcel.2021.04.013.

7. Juszkiewicz, S., and Hegde, R.S. (2018). Quality Control of Orphaned Proteins. Molecular Cell 71, 443–457. 10.1016/j.molcel.2018.07.001.

8. Harper, J.W., and Bennett, E.J. (2016). Proteome complexity and the forces that drive proteome imbalance. Nature 537, 328–338. 10.1038/nature19947.

9. Yanagitani, K., Juszkiewicz, S., and Hegde, R.S. (2017). UBE2O is a quality control factor for orphans of multi-protein complexes. Science 357, 472–475. 10.1126/science.aan0178.

10. Sung, M.-K., Porras-Yakushi, T.R., Reitsma, J.M., Huber, F.M., Sweredoski, M.J., Hoelz, A., Hess, S., and Deshaies, R.J. (2016). A conserved quality-control pathway that mediates degradation of unassembled ribosomal proteins. eLife 5, e19105. 10.7554/eLife.19105.

11. Dephoure, N., Hwang, S., O’Sullivan, C., Dodgson, S.E., Gygi, S.P., Amon, A., and Torres, E.M. (2014). Quantitative proteomic analysis reveals posttranslational responses to aneuploidy in yeast. eLife 3, e03023. 10.7554/eLife.03023.

12. Manchado, E., Guillamot, M., and Malumbres, M. (2012). Killing cells by targeting mitosis. Cell Death Differ 19, 369–377. 10.1038/cdd.2011.197.

13. Natarajan, N., Foresti, O., Wendrich, K., Stein, A., and Carvalho, P. (2020). Quality Control of Protein Complex Assembly by a Transmembrane Recognition Factor. Molecular Cell 77, 108–119.e9. 10.1016/j.molcel.2019.10.003.

14. Babu, M., Vlasblom, J., Pu, S., Guo, X., Graham, C., Bean, B.D.M., Burston, H.E., Vizeacoumar, F.J., Snider, J., Phanse, S., et al. (2012). Interaction landscape of membrane-protein complexes in Saccharomyces cerevisiae. Nature 489, 585–589. 10.1038/nature11354.

15. Vilardi, F., Lorenz, H., and Dobberstein, B. (2011). WRB is the receptor for TRC40/Asna1-mediated insertion of tail-anchored proteins into the ER membrane. Journal of Cell Science 124, 1301–1307. 10.1242/jcs.084277.

16. Vilardi, F., Stephan, M., Clancy, A., Janshoff, A., and Schwappach, B. (2014). WRB and CAML Are Necessary and Sufficient to Mediate Tail-Anchored Protein Targeting to the ER Membrane. PLOS ONE 9, e85033. 10.1371/journal.pone.0085033.

17. Yamamoto, Y., and Sakisaka, T. (2012). Molecular Machinery for Insertion of Tail-Anchored Membrane Proteins into the Endoplasmic Reticulum Membrane in Mammalian Cells. Molecular Cell 48, 387–397. 10.1016/j.molcel.2012.08.028.

18. Colombo, S.F., Cardani, S., Maroli, A., Vitiello, A., Soffientini, P., Crespi, A., Bram, R.F., Benfante, R., and Borgese, N. (2016). Tail-anchored Protein Insertion in Mammals: FUNCTION AND RECIPROCAL INTERACTIONS OF THE TWO SUBUNITS OF THE TRC40 RECEPTOR *. Journal of Biological Chemistry 291, 15292–15306. 10.1074/jbc.M115.707752.

19. Rivera-Monroy, J., Musiol, L., Unthan-Fechner, K., Farkas, Á., Clancy, A., Coy-Vergara, J., Weill, U., Gockel, S., Lin, S.-Y., Corey, D.P., et al. (2016). Mice lacking WRB reveal differential biogenesis requirements of tail-anchored proteins in vivo. Sci Rep 6, 39464. 10.1038/srep39464.

20. McDowell, M.A., Heimes, M., Fiorentino, F., Mehmood, S., Farkas, Á., Coy-Vergara, J., Wu, D., Bolla, J.R., Schmid, V., Heinze, R., et al. (2020). Structural Basis of Tail-Anchored Membrane Protein Biogenesis by the GET Insertase Complex. Molecular Cell 80, 72–86.e7. 10.1016/j.molcel.2020.08.012.

21. McDowell, M.A., Heimes, M., Enkavi, G., Farkas, Á., Saar, D., Wild, K., Schwappach, B., Vattulainen, I., and Sinning, I. (2023). The GET insertase exhibits conformational plasticity and induces membrane thinning. Nat Commun 14, 7355. 10.1038/s41467-023-42867-2.

22. Wang, F., Chan, C., Weir, N.R., and Denic, V. (2014). The Get1/2 transmembrane complex is an endoplasmic-reticulum membrane protein insertase. Nature 512, 441–444. 10.1038/nature13471.

23. Cymer, F., Veerappan, A., and Schneider, D. (2012). Transmembrane helix–helix interactions are modulated by the sequence context and by lipid bilayer properties. Biochimica et Biophysica Acta (BBA) - Biomembranes 1818, 963–973. 10.1016/j.bbamem.2011.07.035.

24. Pazour, G.J., and Bloodgood, R.A. (2008). Chapter 5 Targeting Proteins to the Ciliary Membrane. In Current Topics in Developmental Biology (Elsevier), pp. 115–149. 10.1016/S0070-2153(08)00805-3.

25. Mill, P., Christensen, S.T., and Pedersen, L.B. (2023). Primary cilia as dynamic and diverse signalling hubs in development and disease. Nat Rev Genet 24, 421–441. 10.1038/s41576-023-00587-9.

26. The Primary Cilium as a Hedgehog Signal Transduction Machine (2009). In Methods in Cell Biology (Academic Press), pp. 199–222. 10.1016/S0091-679X(08)94010-3.

27. Reiter, J.F., and Leroux, M.R. (2017). Genes and molecular pathways underpinning ciliopathies. Nat Rev Mol Cell Biol 18, 533–547. 10.1038/nrm.2017.60.

28. Roberson, E.C., Dowdle, W.E., Ozanturk, A., Garcia-Gonzalo, F.R., Li, C., Halbritter, J., Elkhartoufi, N., Porath, J.D., Cope, H., Ashley-Koch, A., et al. (2015). TMEM231, mutated in orofaciodigital and Meckel syndromes, organizes the ciliary transition zone. J Cell Biol 209, 129–142. 10.1083/jcb.201411087.

29. Srour, M., Hamdan, F.F., Schwartzentruber, J.A., Patry, L., Ospina, L.H., Shevell, M.I., Désilets, V., Dobrzeniecka, S., Mathonnet, G., Lemyre, E., et al. (2012). Mutations in TMEM231 cause Joubert syndrome in French Canadians. Journal of Medical Genetics 49, 636–641. 10.1136/jmedgenet-2012-101132.

30. Shaheen, R., Ansari, S., Mardawi, E.A., Alshammari, M.J., and Alkuraya, F.S. (2013). Mutations in TMEM231 cause Meckel–Gruber syndrome. Journal of Medical Genetics 50, 160–162. 10.1136/jmedgenet-2012-101431.

31. Inglis, A.J., Page, K.R., Guna, A., and Voorhees, R.M. (2020). Differential Modes of Orphan Subunit Recognition for the WRB/CAML Complex. Cell Reports 30, 3691–3698.e5. 10.1016/j.celrep.2020.02.084.

32. Itakura, E., Zavodszky, E., Shao, S., Wohlever, M.L., Keenan, R.J., and Hegde, R.S. (2016). Ubiquilins Chaperone and Triage Mitochondrial Membrane Proteins for Degradation. Molecular Cell 63, 21–33. 10.1016/j.molcel.2016.05.020.

33. Gilbert, L.A., Horlbeck, M.A., Adamson, B., Villalta, J.E., Chen, Y., Whitehead, E.H., Guimaraes, C., Panning, B., Ploegh, H.L., Bassik, M.C., et al. (2014). Genome-Scale CRISPR-Mediated Control of Gene Repression and Activation. Cell 159, 647–661. 10.1016/j.cell.2014.09.029.

34. Horlbeck, M.A., Gilbert, L.A., Villalta, J.E., Adamson, B., Pak, R.A., Chen, Y., Fields, A.P., Park, C.Y., Corn, J.E., Kampmann, M., et al. (2016). Compact and highly active next-generation libraries for CRISPR-mediated gene repression and activation. eLife 5, e19760. 10.7554/eLife.19760.

35. Schulman, B.A., and Wade Harper, J. (2009). Ubiquitin-like protein activation by E1 enzymes: the apex for downstream signalling pathways. Nat Rev Mol Cell Biol 10, 319–331. 10.1038/nrm2673.

36. Olzmann, J.A., Kopito, R.R., and Christianson, J.C. (2013). The Mammalian Endoplasmic Reticulum-Associated Degradation System. Cold Spring Harb Perspect Biol 5, a013185. 10.1101/cshperspect.a013185.

37. Olzmann, J.A., Richter, C.M., and Kopito, R.R. (2013). Spatial regulation of UBXD8 and p97/VCP controls ATGL-mediated lipid droplet turnover. Proceedings of the National Academy of Sciences 110, 1345–1350. 10.1073/pnas.1213738110.

38. Shaheen, R., Szymanska, K., Basu, B., Patel, N., Ewida, N., Faqeih, E., Al Hashem, A., Derar, N., Alsharif, H., Aldahmesh, M.A., et al. (2016). Characterizing the morbid genome of ciliopathies. Genome Biology 17, 242. 10.1186/s13059-016-1099-5.

39. Radhakrishnan, P., Nayak, S.S., Shukla, A., Lindstrand, A., and Girisha, K.M. (2019). Meckel syndrome: Clinical and mutation profile in six fetuses. Clinical Genetics 96, 560–565. 10.1111/cge.13623.

40. Ridnõi, K., Šois, M., Vaidla, E., Pajusalu, S., Kelder, L., Reimand, T., and Õunap, K. (2019). A prenatally diagnosed case of Meckel–Gruber syndrome with novel compound heterozygous pathogenic variants in the TXNDC15 gene. Molecular Genetics & Genomic Medicine 7, e614. 10.1002/mgg3.614.

41. Yamazaki, S., Fujii, T., Chiba, S., Shin, H.-W., Nakayama, K., and Katoh, Y. (2024). TXNDC15, an ER-localized thioredoxin-like transmembrane protein, contributes to ciliary transition zone integrity. Journal of Cell Science 137, jcs262123. 10.1242/jcs.262123.

42. Breslow, D.K., Hoogendoorn, S., Kopp, A.R., Morgens, D.W., Vu, B.K., Kennedy, M.C., Han, K., Li, A., Hess, G.T., Bassik, M.C., et al. (2018). A CRISPR-based screen for Hedgehog signaling provides insights into ciliary function and ciliopathies. Nat Genet 50, 460–471. 10.1038/s41588-018-0054-7.

43. Guna, A., Page, K.R., Replogle, J.M., Esantsi, T.K., Wang, M.L., Weissman, J.S., and Voorhees, R.M. (2023). A dual sgRNA library design to probe genetic modifiers using genome-wide CRISPRi screens. BMC Genomics 24, 651. 10.1186/s12864-023-09754-y.

44. Lilley, B.N., and Ploegh, H.L. (2004). A membrane protein required for dislocation of misfolded proteins from the ER. Nature 429, 834–840. 10.1038/nature02592.

45. Bays, N.W., Gardner, R.G., Seelig, L.P., Joazeiro, C.A., and Hampton, R.Y. (2001). Hrd1p/Der3p is a membrane-anchored ubiquitin ligase required for ER-associated degradation. Nat Cell Biol 3, 24–29. 10.1038/35050524.

46. Stein, A., Ruggiano, A., Carvalho, P., and Rapoport, T.A. (2014). Key Steps in ERAD of Luminal ER Proteins Reconstituted with Purified Components. Cell 158, 1375–1388. 10.1016/j.cell.2014.07.050.

47. Christianson, J.C., Shaler, T.A., Tyler, R.E., and Kopito, R.R. (2008). OS-9 and GRP94 deliver mutant α1-antitrypsin to the Hrd1–SEL1L ubiquitin ligase complex for ERAD. Nat Cell Biol 10, 272–282. 10.1038/ncb1689.

48. Christianson, J.C., Olzmann, J.A., Shaler, T.A., Sowa, M.E., Bennett, E.J., Richter, C.M., Tyler, R.E., Greenblatt, E.J., Harper, J.W., and Kopito, R.R. (2011). Defining human ERAD networks through an integrative mapping strategy. Nat Cell Biol 14, 93–105. 10.1038/ncb2383.

49. Soldà, T., Galli, C., Guerra, C., Hoefner, C., and Molinari, M. (2024). TMX5/TXNDC15, a natural trapping mutant of the PDI family is a client of the proteostatic factor ERp44. Life Sci. Alliance 7, e202403047. 10.26508/lsa.202403047.

50. Hassink, G., Kikkert, M., Voorden, S. van, Lee, S.-J., Spaapen, R., Laar, T. van, Coleman, C.S., Bartee, E., Früh, K., Chau, V., et al. (2005). TEB4 is a C4HC3 RING finger-containing ubiquitin ligase of the endoplasmic reticulum. Biochemical Journal 388, 647–655. 10.1042/BJ20041241.

51. Habeck, G., Ebner, F.A., Shimada-Kreft, H., and Kreft, S.G. (2015). The yeast ERAD-C ubiquitin ligase Doa10 recognizes an intramembrane degron. Journal of Cell Biology 209, 261–273. 10.1083/jcb.201408088.

52. Kreft, S.G., Wang, L., and Hochstrasser, M. (2006). Membrane Topology of the Yeast Endoplasmic Reticulum-localized Ubiquitin Ligase Doa10 and Comparison with Its Human Ortholog TEB4 (MARCH-VI)*. Journal of Biological Chemistry 281, 4646–4653. 10.1074/jbc.M512215200.

53. Schmidt, C.C., Vasic, V., and Stein, A. (2020). Doa10 is a membrane protein retrotranslocase in ER-associated protein degradation. eLife 9, e56945. 10.7554/eLife.56945.

54. Fenech, E.J., Lari, F., Charles, P.D., Fischer, R., Laétitia-Thézénas, M., Bagola, K., Paton, A.W., Paton, J.C., Gyrd-Hansen, M., Kessler, B.M., et al. (2020). Interaction mapping of endoplasmic reticulum ubiquitin ligases identifies modulators of innate immune signalling. eLife 9, e57306. 10.7554/eLife.57306.

55. Timms, R.T., Menzies, S.A., Tchasovnikarova, I.A., Christensen, L.C., Williamson, J.C., Antrobus, R., Dougan, G., Ellgaard, L., and Lehner, P.J. (2016). Genetic dissection of mammalian ERAD through comparative haploid and CRISPR forward genetic screens. Nat Commun 7, 11786. 10.1038/ncomms11786.

56. Yang, J., Kim, S.-Y., and Hwang, C.-S. (2024). Delineation of the substrate recognition domain of MARCHF6 E3 ubiquitin ligase in the Ac/N-degron pathway and its regulatory role in ferroptosis. J Biol Chem 300, 107731. 10.1016/j.jbc.2024.107731.

57. Sine, S.M., and Engel, A.G. (2006). Recent advances in Cys-loop receptor structure and function. Nature 440, 448–455. 10.1038/nature04708.

58. Chih, B., Liu, P., Chinn, Y., Chalouni, C., Komuves, L.G., Hass, P.E., Sandoval, W., and Peterson, A.S. (2012). A ciliopathy complex at the transition zone protects the cilia as a privileged membrane domain. Nat Cell Biol 14, 61–72. 10.1038/ncb2410.

59. Smith, U.M., Consugar, M., Tee, L.J., McKee, B.M., Maina, E.N., Whelan, S., Morgan, N.V., Goranson, E., Gissen, P., Lilliquist, S., et al. (2006). The transmembrane protein meckelin (MKS3) is mutated in Meckel-Gruber syndrome and the wpk rat. Nat Genet 38, 191–196. 10.1038/ng1713.

60. Botsch, J.J., Junker, R., Sorgenfrei, M., Ogger, P.P., Stier, L., von Gronau, S., Murray, P.J., Seeger, M.A., Schulman, B.A., and Bräuning, B. (2024). Doa10/MARCH6 architecture interconnects E3 ligase activity with lipid-binding transmembrane channel to regulate SQLE. Nat Commun 15, 410. 10.1038/s41467-023-44670-5.

61. Wu, K., Itskanov, S., Lynch, D.L., Chen, Y., Turner, A., Gumbart, J.C., and Park, E. (2024). Substrate recognition mechanism of the endoplasmic reticulum-associated ubiquitin ligase Doa10. Nat Commun 15, 2182. 10.1038/s41467-024-46409-2.

62. Uhlén, M., Fagerberg, L., Hallström, B.M., Lindskog, C., Oksvold, P., Mardinoglu, A., Sivertsson, Å., Kampf, C., Sjöstedt, E., Asplund, A., et al. (2015). Tissue-based map of the human proteome. Science 347, 1260419. 10.1126/science.1260419.

63. Žegarac, Ž., Duić, Ž., and Bojanić, K. (2019). Meckel Gruber Syndrome: A rare and lethal anomaly. Clinical Journal of Obstetrics and Gynecology 2, 127–132. 10.29328/journal.cjog.1001035.

64. Skowyra, D., Craig, K.L., Tyers, M., Elledge, S.J., and Harper, J.W. (1997). F-Box Proteins Are Receptors that Recruit Phosphorylated Substrates to the SCF Ubiquitin-Ligase Complex. Cell 91, 209–219. 10.1016/S0092-8674(00)80403-1.

65. Madsen, L., Seeger, M., Semple, C.A., and Hartmann-Petersen, R. (2009). New ATPase regulators--p97 goes to the PUB. Int J Biochem Cell Biol 41, 2380–2388. 10.1016/j.biocel.2009.05.017.

66. Zuiderweg, E.R.P., Hightower, L.E., and Gestwicki, J.E. (2017). The remarkable multivalency of the Hsp70 chaperones. Cell Stress Chaperones 22, 173–189. 10.1007/s12192-017-0776-y.

67. Jost, M., Chen, Y., Gilbert, L.A., Horlbeck, M.A., Krenning, L., Menchon, G., Rai, A., Cho, M.Y., Stern, J.J., Prota, A.E., et al. (2017). Combined CRISPRi/a-Based Chemical Genetic Screens Reveal that Rigosertib Is a Microtubule-Destabilizing Agent. Mol Cell 68, 210–223.e6. 10.1016/j.molcel.2017.09.012.

68. Murray, T.A., Liu, Q., Whiteaker, P., Wu, J., and Lukas, R.J. (2009). Nicotinic acetylcholine receptor α7 subunits with a C2 cytoplasmic loop yellow fluorescent protein insertion form functional receptors. Acta Pharmacol Sin 30, 828–841. 10.1038/aps.2009.78.

69. Feng, S., Sekine, S., Pessino, V., Li, H., Leonetti, M.D., and Huang, B. (2017). Improved split fluorescent proteins for endogenous protein labeling. Nat Commun 8, 370. 10.1038/s41467-017-00494-8.

70. Götzke, H., Kilisch, M., Martínez-Carranza, M., Sograte-Idrissi, S., Rajavel, A., Schlichthaerle, T., Engels, N., Jungmann, R., Stenmark, P., Opazo, F., et al. (2019). The ALFA-tag is a highly versatile tool for nanobody-based bioscience applications. Nat Commun 10, 4403. 10.1038/s41467-019-12301-7.

71. Munro, S., and Pelham, H.R.B. (1987). A C-terminal signal prevents secretion of luminal ER proteins. Cell 48, 899–907. 10.1016/0092-8674(87)90086-9.

72. Arai, R., Ueda, H., Kitayama, A., Kamiya, N., and Nagamune, T. (2001). Design of the linkers which effectively separate domains of a bifunctional fusion protein. Protein Engineering, Design and Selection 14, 529–532. 10.1093/protein/14.8.529.

73. Replogle, J.M., Norman, T.M., Xu, A., Hussmann, J.A., Chen, J., Cogan, J.Z., Meer, E.J., Terry, J.M., Riordan, D.P., Srinivas, N., et al. (2020). Combinatorial single-cell CRISPR screens by direct guide RNA capture and targeted sequencing. Nat Biotechnol 38, 954–961. 10.1038/s41587-020-0470-y.

74. Pleiner, T., Bates, M., and Görlich, D. (2017). A toolbox of anti–mouse and anti–rabbit IgG secondary nanobodies. Journal of Cell Biology 217, 1143–1154. 10.1083/jcb.201709115.

75. Zhang, J.-P., Li, X.-L., Li, G.-H., Chen, W., Arakaki, C., Botimer, G.D., Baylink, D., Zhang, L., Wen, W., Fu, Y.-W., et al. (2017). Efficient precise knockin with a double cut HDR donor after CRISPR/Cas9-mediated double-stranded DNA cleavage. Genome Biol 18, 35. 10.1186/s13059-017-1164-8.

76. Yurtsev, E., and Friedman, J. (2015). FlowCytometryTools. Version zenodo (Zenodo). 10.5281/zenodo.32992 https://doi.org/10.5281/zenodo.32992.

77. Stevens, T.A., Tomaleri, G.P., Hazu, M., Wei, S., Nguyen, V.N., DeKalb, C., Voorhees, R.M., and Pleiner, T. (2024). A nanobody-based strategy for rapid and scalable purification of human protein complexes. Nat Protoc 19, 127–158. 10.1038/s41596-023-00904-w.

78. Jones, J., MacKrell, E.J., Wang, T.-Y., Lomenick, B., Roukes, M.L., and Chou, T.-F. (2023). Tidyproteomics: an open-source R package and data object for quantitative proteomics post analysis and visualization. BMC Bioinformatics 24, 239. 10.1186/s12859-023-05360-7.

79. Wickham, H. (2016). ggplot2: Elegant Graphics for Data Analysis (Springer International Publishing) 10.1007/978-3-319-24277-4.

80. Deutsch, E.W., Bandeira, N., Sharma, V., Perez-Riverol, Y., Carver, J.J., Kundu, D.J., García-Seisdedos, D., Jarnuczak, A.F., Hewapathirana, S., Pullman, B.S., et al. (2020). The ProteomeXchange consortium in 2020: enabling ‘big data’ approaches in proteomics. Nucleic Acids Research 48, D1145–D1152. 10.1093/nar/gkz984.

81. Sharma, A., Mariappan, M., Appathurai, S., and Hegde, R.S. (2010). In vitro dissection of protein translocation into the mammalian endoplasmic reticulum. Methods Mol Biol 619, 339–363. 10.1007/978-1-60327-412-8_20.

82. Walter, P., and Blobel, G. (1983). Preparation of microsomal membranes for cotranslational protein translocation. Methods Enzymol 96, 84–93. 10.1016/s0076-6879(83)96010-x.

83. Schindelin, J., Arganda-Carreras, I., Frise, E., Kaynig, V., Longair, M., Pietzsch, T., Preibisch, S., Rueden, C., Saalfeld, S., Schmid, B., et al. (2012). Fiji: an open-source platform for biological-image analysis. Nat Methods 9, 676–682. 10.1038/nmeth.2019.

84. Hansen, J.N., Rassmann, S., Stüven, B., Jurisch-Yaksi, N., and Wachten, D. (2021). CiliaQ: a simple, open-source software for automated quantification of ciliary morphology and fluorescence in 2D, 3D, and 4D images. Eur Phys J E Soft Matter 44, 18. 10.1140/epje/s10189-021-00031-y.

85. Griessl, M., Gutknecht, M., and Cook, C.H. (2017). Determination of suitable reference genes for RT-qPCR analysis of murine Cytomegalovirus in vivo and in vitro. Journal of Virological Methods 248, 100–106. 10.1016/j.jviromet.2017.06.012.

86. Wang, X., and Seed, B. (2003). A PCR primer bank for quantitative gene expression analysis. Nucleic Acids Res 31, e154. 10.1093/nar/gng154.

87. Spandidos, A., Wang, X., Wang, H., Dragnev, S., Thurber, T., and Seed, B. (2008). A comprehensive collection of experimentally validated primers for Polymerase Chain Reaction quantitation of murine transcript abundance. BMC Genomics 9, 633. 10.1186/1471-2164-9-633.

88. Spandidos, A., Wang, X., Wang, H., and Seed, B. (2010). PrimerBank: a resource of human and mouse PCR primer pairs for gene expression detection and quantification. Nucleic Acids Res 38, D792–799. 10.1093/nar/gkp1005.

89. Pfaffl, M.W. (2001). A new mathematical model for relative quantification in real-time RT-PCR. Nucleic Acids Res 29, e45. 10.1093/nar/29.9.e45.

90. Waterhouse, A.M., Procter, J.B., Martin, D.M.A., Clamp, M., and Barton, G.J. (2009). Jalview Version 2— a multiple sequence alignment editor and analysis workbench. Bioinformatics 25, 1189–1191. 10.1093/bioinformatics/btp033.

91. Abramson, J., Adler, J., Dunger, J., Evans, R., Green, T., Pritzel, A., Ronneberger, O., Willmore, L., Ballard, A.J., Bambrick, J., et al. (2024). Accurate structure prediction of biomolecular interactions with AlphaFold 3. Nature 630, 493–500. 10.1038/s41586-024-07487-w.

