## Supplemental figures and legends for "TXNDC15 modulated quality control at the endoplasmic reticulum shapes ciliogenesis"

[1]Division of Biology and Biological Engineering, California Institute of Technology, Pasadena, CA 91125, USA
[2]Current address: Altos Labs, San Francisco, CA 94065, USA
[3]Current address: Addition Therapeutics, San Francisco, CA 94080, USA
[4]David Geffen School of Medicine, University of California Los Angeles, Los Angeles, CA 90095, USA
[5] Current address: Department of Biochemistry and Biophysics, University of California San Francisco, San Francisco, CA 94143, USA.
[6]Howard Hughes Medical Institute, California Institute of Technology, Pasadena, CA 91125, USA

### SUPPLEMENTAL FIGURES AND LEGENDS

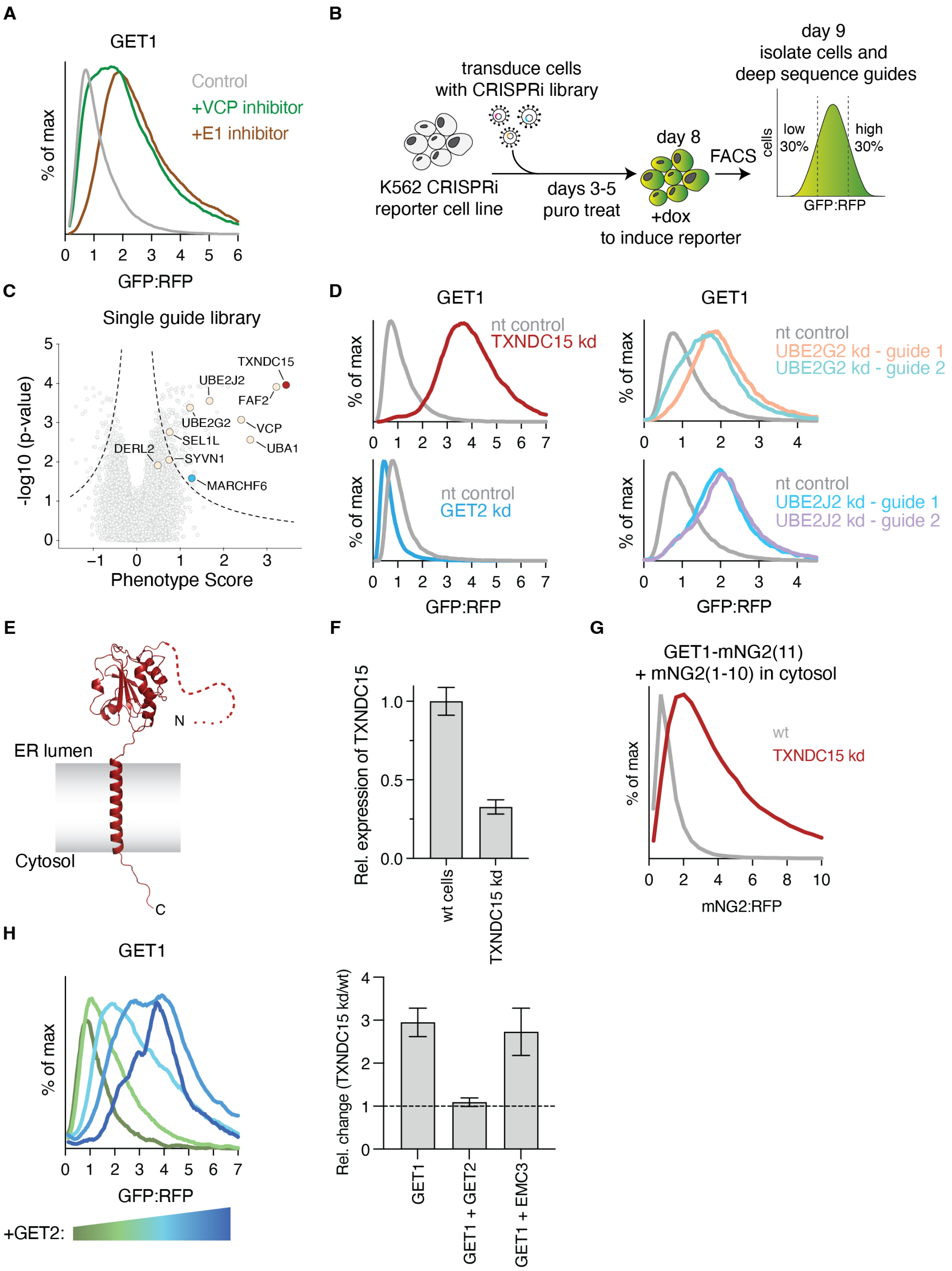


**Figure S1. A platform for genome wide CRISPRi screens for quality control of orphan GET1, related to Figure 1**

(A) Unassembled GET1 is stabilized upon inhibition of the ubiquitin proteasome pathway. A GET1-GFP reporter construct was expressed in HEK293T cells along with a normalization control (RFP). Cells were treated with either the VCP inhibitor CB-5083 (250 nM) or the E1 inhibitor MLN7243 (1.25 µM) for 6 hours before analysis by flow cytometry. Stabilization of GET1-GFP relative to a normalization marker (RFP) upon drug treatment is consistent with its

degradation by the ubiquitin proteasome system. For all experiments histograms are normalized such that the GFP:RFP ratio in wt cells is 1, as described in the methods.
(B) Workflow of the FACS-based CRISPRi screen used to identify quality control factors for orphan GET1. A K562 CRISPRi reporter cell line was constructed containing the GET1-GFP reporter under an inducible promoter. Following transduction with a genome wide v2-CRISPRi sgRNA library,[34] the GET1-GFP reporter was induced with doxycycline for 36 hours prior to cell sorting. Cells were isolated based on ratiometric changes in GFP relative to RFP expression by FACS and the enriched sgRNAs were identified using deep sequencing.
(C) Volcano plot of the genome wide CRISPRi screen for orphan GET1 highlighting known ERAD factors. Genes falling outside of the indicated dashed lines are considered statistically significant hits, including the indicated ERAD factors.
(D) Stability of the GET1 reporter was assessed in K562 cells expressing a nontargeting (nt) control or the indicated sgRNA. GET1-GFP expression relative to a normalization control was determined by flow cytometry and displayed as a histogram. Consistent with the results of the screen, depletion of TXNDC15 or the indicated ER-resident E2 ubiquitin-conjugating enzymes (refer to (C)) leads to stabilization of GET1 while depletion of GET2 leads to destabilization of GET1.
(E) AlphaFold 3 predicted model of TXNDC15. After the signal sequence, the N-terminal 175 residues of the mature protein are predicted to be unstructured followed by a thioredoxin-like domain in the ER lumen and a single C-terminal TM.
(F) Characterization of a HEK293T TXNDC15 stable knockdown cell line. HEK293T cells were transiently transfected with Cas9 and an sgRNA targeting TXNDC15, along with GFP as a transfection marker. Transfected cells were sorted to generate a monoclonal population. Relative expression of TXNDC15 in a stable HEK293T TXNDC15 kd cell line compared to a wt control was quantified by qPCR. Error bars represent errors as determined by error propagation, calculated from the primer efficiency and measurements taken from 3 technical replicates.
(G) To determine whether labeling of GET1 with GFP influences its dependence on TXNDC15, a C-terminal fusion of GET1 with the 11$^{th}$ b-strand of monomeric green2 (mNG2(11)) was expressed in HEK293T wt and TXNDC15 kd cells. At the same time, the remaining ten b-strands of mNG2 (mNG2(1-10)) were expressed in the cytosol. Binding of mNG2(1-10) to mNG2(11) results in complementation and thereby fluorescence relative to a normalization control (RFP). Using our ratiometric reporter assay we determined that the post-translational stability of GET1-mNG2(11) was also dependent on TXNDC15 kd, excluding artifacts of labeling GET1 with full length GFP.
(H) (Left) GET2 expression stabilizes the ratiometric GET1 fluorescent reporter. The GET1-GFP-2A-RFP reporter construct used for genome wide screens described in Figure 1 was transduced in HEK293T cells along with GET2-2A-BFP and analyzed by flow cytometry. All histograms were normalized relative to the median of the GFP:RFP-ratio in wt cells without GET2. Increasing amounts of BFP expression, a proxy for GET2 levels, leads to a commensurate increase in GET1-GFP:RFP ratio. This GET2 dependent stabilization of GET1 suggests that our ratiometric reporter is predominantly unassembled and can therefore be used to study the quality control of orphan GET1. (Right) Additionally, the relative change in GET1 stability in TXNDC15 depleted compared to wt HEK293T cells was measured in the presence and absence of saturating levels of exogenously expressed GET2. As a specificity control, the effect of EMC3 expression on GET1 stability is also displayed. The relative change (TXNDC15 kd/wt) was calculated from the medians of the cell distributions and is shown here as a bar graph. The dashed line at 1 is the relative change expected for substrates that are not dependent on TXNDC15.

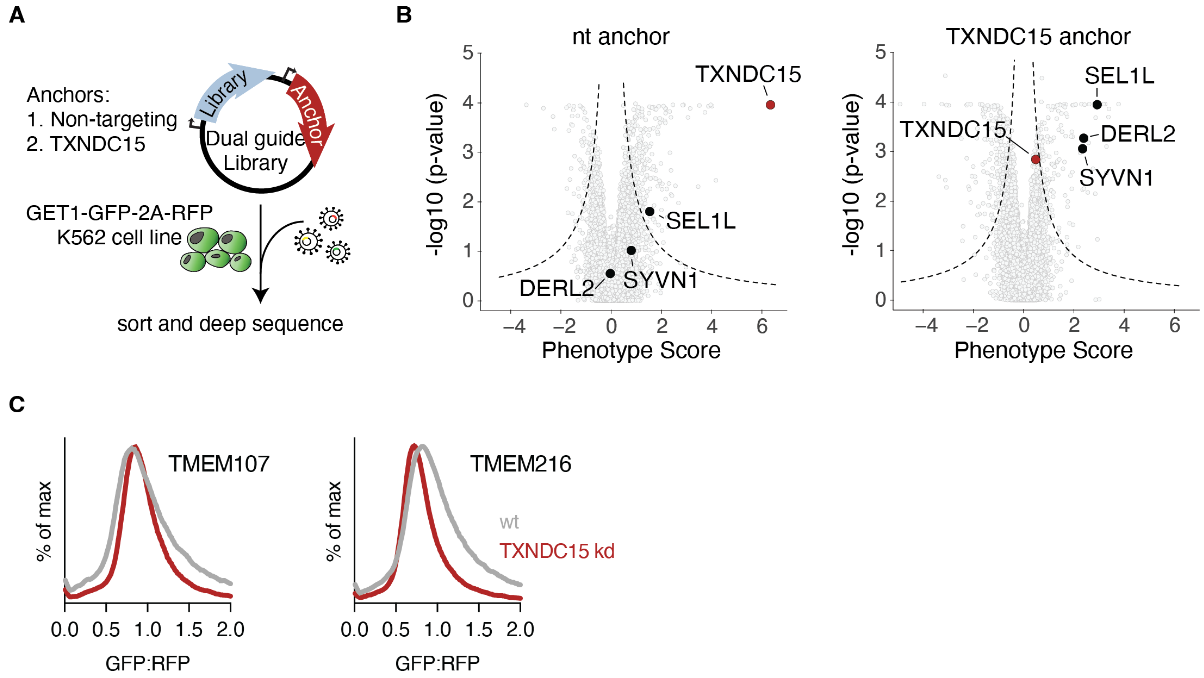


**Figure S2. TXNDC15 acts in a parallel pathway to canonical ERAD factors like SYVN1 and has no effect on additional MKS complex proteins, related to Figure 1**

(A) Schematic of the dual guide genetic modifier screening approach.[43]

(B) To identify genetic modifiers of TXNDC15 across the genome we performed two screens using dual libraries expressing either a nt control (left) or TXNDC15 guide (right) at the anchor position and all guides in the CRISPRi-v2 library at the variable position. Volcano plots summarizing the resulting CRISPRi modifier screens are shown, and statistically significant hits indicated by the dashed lines. TXNDC15 (red) and several known ERAD factors (black) are highlighted.

(B) The indicated membrane protein subunits of the MKS complex were analyzed in HEK293T wildtype (wt) and TXNDC15 knockdown (kd) cells. Histograms were normalized to the median GFP:RFP ratio of the reporter in wt cells, only minor changes can be observed.

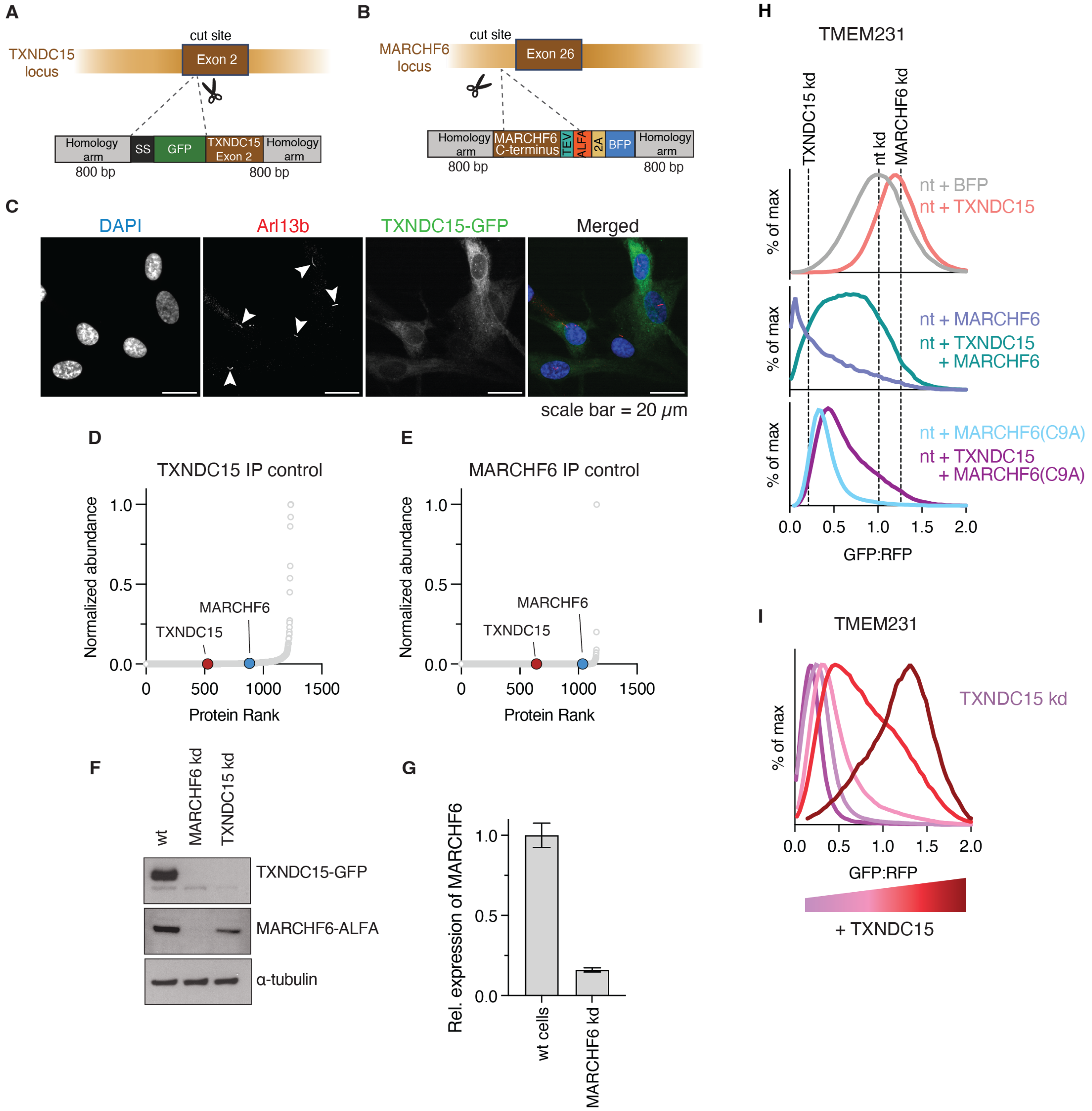


**Figure S3. TXNDC15 acts together with MARCHF6 in the ER, related to Figure 2**

(A) Schematic of strategy for tagging endogenous TXNDC15 with a GFP at the N-terminus of the mature domain in HEK293T cells. A vector encoding GFP and 800 bp of homology on either side of the PAM cas9 cut site within the TXNDC15 locus was cloned. The GFP insertion site was designed such that the native TXNDC15 signal sequence (SS) and signal cleavage site would remain intact (see materials and methods).

(B) Schematic of strategy for tagging endogenous MARCHF6 with an ALFA tag. A ribosomal skipping site 2A followed by BFP was added for subsequent cell sorting.

(C) Confocal imaging of RPE1 cells transduced with TXNDC15-GFP (green) subjected to serum starvation to induce cilia formation. Shown is staining with DAPI (blue) for nuclear DNA and Arl13b (red) as a ciliary marker. Cilia are demarcated with arrows in the Arl13b channel. Even upon overexpression of TXNDC15, we did not observe colocalization of TXNDC15 with the

Arl13b marker. These data, together with imaging of the endogenously tagged GFP-TXNDC15 shown in Figure 2A suggested that TXNDC15 is localized primarily to the ER.

(D) To ensure that the observed interaction of MARCHF6 with TXNDC15 shown in Figure 2B was specific, we used wt HEK293T cells containing untagged TXNDC15 as the background control for immunoprecipitation of endogenously tagged GFP-TXNDC15. Non-specific interaction partners of the GFP nanobody were analyzed by mass spectrometry and were ranked based on the abundance and plotted as protein rank versus abundance normalized on the most abundant protein. No enrichment of TXNDC15 or MARCHF6 was detected.

(E) Mass spectrometry analysis of the background control analyzed as in (D) for the immunoprecipitation of exogenously expressed MARCHF6 in Figure 2C. Expi293 cells overexpressing MARCHF6(C9A)-GFP were purified in the absence of the GFP nanobody to assess non-specific interactors. Similar to (D), MARCHF6 and TXNDC15 were not enriched under these conditions, suggesting that the interaction between MARCHF6 and TXNDC15 is specific.

(F) HEK293T cells tagged at the endogenous loci GFP-TXNDC15 and/or MARCHF6-ALFA were depleted of either TXNDC15 or MARCHF6 and analyzed by western blot. While depletion of TXNDC15 in MARCHF6-ALFA tagged cells reduced the amounts of MARCHF6, depletion of MARCHF6 in GFP-TXNDC15 tagged cells resulted in loss of TXNDC15 beyond the detection limit. As a control, blots of double-tagged GFP-TXNDC15 MARCHF6-ALFA cells are shown.

(G) Characterization of a HEK293T MARCHF6 stable knockdown cell line. HEK293T cells were transiently transfected with Cas9 and an sgRNA targeting MARCHF6, along with GFP as a transfection marker. Transfected cells were sorted. Relative expression of MARCHF6 in a HEK293T MARCHF6 stable kd cell line compared to a wt control was quantified by qPCR. Error bars represent errors as determined by error propagation, calculated from the primer efficiency and measurements taken from 3 technical replicates.

(H) To delineate the contributions of MARCHF6 and TXNDC15 to TMEM231 stability, we tested how their overexpression in wildtype cells affected the stability of TMEM231. Data for part of this experiment is presented in Figure 2H. K562 cells carrying an inducible TMEM231 reporter were transduced with a non-targeting (nt) control sgRNA. Afterwards, cDNA constructs expressing TXNDC15, MARCHF6, or the catalytically inactive mutant MARCHF6(C9A) were overexpressed and reporter stability was determined by flow cytometry. Histograms were normalized to the median of the non-targeting control which is the same as shown in Figure 2H. The mode of the nt kd, TXNDC15 kd and MARCHF6 kd are indicated by vertical dotted lines (for reference, histograms are shown in Figure 2H). Analysis of these four conditions suggests the following. (i) Overexpression of TXNDC15 alone phenocopies MARCHF6 kd, resulting in stabilization of TMEM231. This is consistent with a model where altering the equilibrium of TXNDC15-MARCHF6 binding such that all MARCHF6 is occupied by TXNDC15 prevents degradation of TMEM231. (ii) Overexpression of MARCHF6 alone, so that it is expressed in excess of endogenous levels of TXNDC15, destabilized TMEM231. These results suggested that without the protective effect of TXNDC15, MARCHF6 is able to catalyze the degradation of TMEM231. (iii) Overexpression of both TXNDC15 and MARCHF6 introduces more complexed and free MARCHF6 into the ER, which led to a small destabilization of TMEM231 compared to overexpression of only MARCHF6. (iv) Finally, overexpression of the catalytically inactive MARCHF6(C9A) mutant leads to a destabilization of TMEM231. These results suggest that excess inactive MARCHF6(C9A) binds TXNDC15, thereby freeing endogenous wildtype MARCHF6 to degrade TMEM231.

(I) Using our ratiometric fluorescent reporter assay we tested the effects of TXNDC15 expression on TMEM231 stability. Following TXNDC15 depletion, TXNDC15 was reintroduced by overexpression of TXNDC15-2A-BFP. BFP levels can be used as a proxy for TXNDC15 expression in each cell. Cells were gated for 'no BFP' (TXNDC15 kd) and increasing amounts of BFP (violet to red gradient). Higher expression of TXNDC15 lead to a commensurate increase in TMEM231 stabilization. We therefore concluded that the higher the occupancy of TXNDC15 at MARCHF6, the greater protection of TMEM231 from degradation.

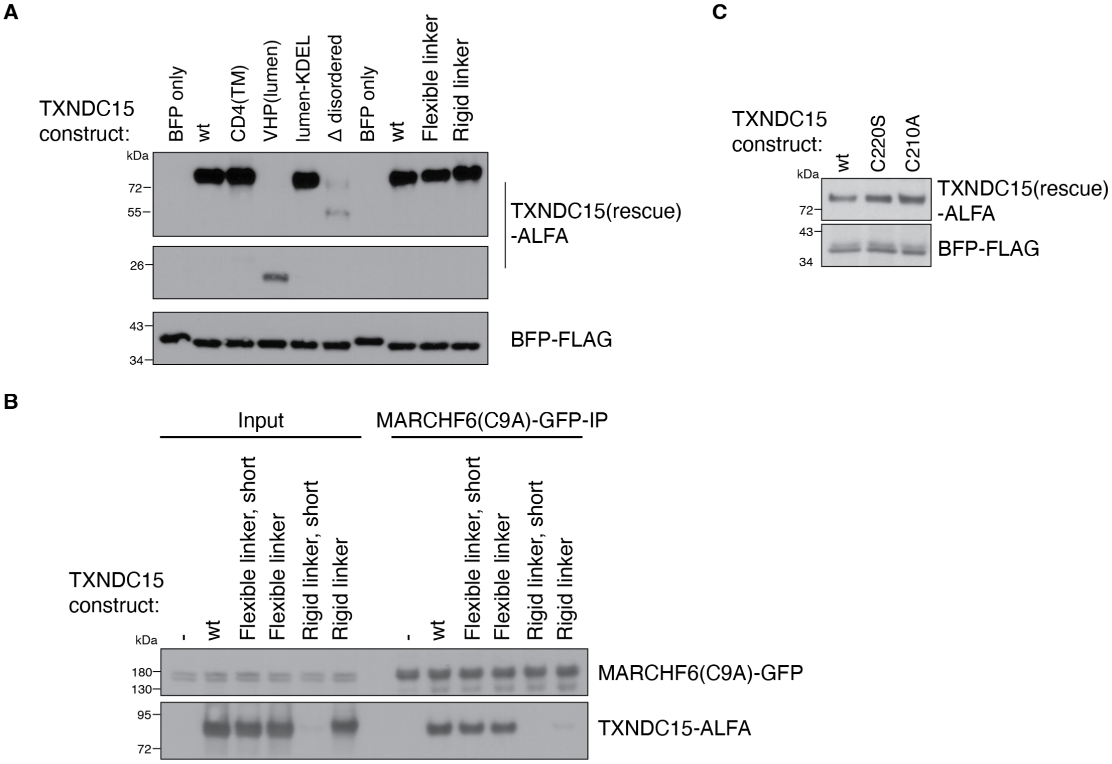


**Figure S4. Mutational analysis of TXNDC15, related to Figure 3**

(A) Western blots of cells expressing the TXNDC15 rescue constructs described in Figure 3A to ensure that mutants accumulate to similar levels. Therefore, the observed differences in their quality control activity cannot be explained by differences in expression level or protein stability. Blots were normalized to BFP levels to account for differences in transduction of the TXNDC15-2A-BFP rescue. Note, though the Δ disordered construct accumulated at lower levels than wildtype TXNDC15, it was still capable of rescuing while the VHP(lumen) construct is not functional despite being expressed at relatively higher levels.

(B) To test whether mutations to TXNDC15 affected their ability to bind MARCHF6, we performed an immunoprecipitation assay in human cells. Expi293 cells expressing MARCHF6(C9A)-GFP were transduced with the TXNDC15-ALFA rescue mutants as indicated. After anti-GFP immunoprecipitation, samples were analyzed by western blot for co-immunoprecipitation of the indicated TXNDC15 mutants. The short versions of flexible and rigid linkers were not included in the final experiment in Figure 3A.

(C) Western blot of the TXNDC15 mutant rescue constructs shown in Figure 3D in HEK293T cells, as described in (A).

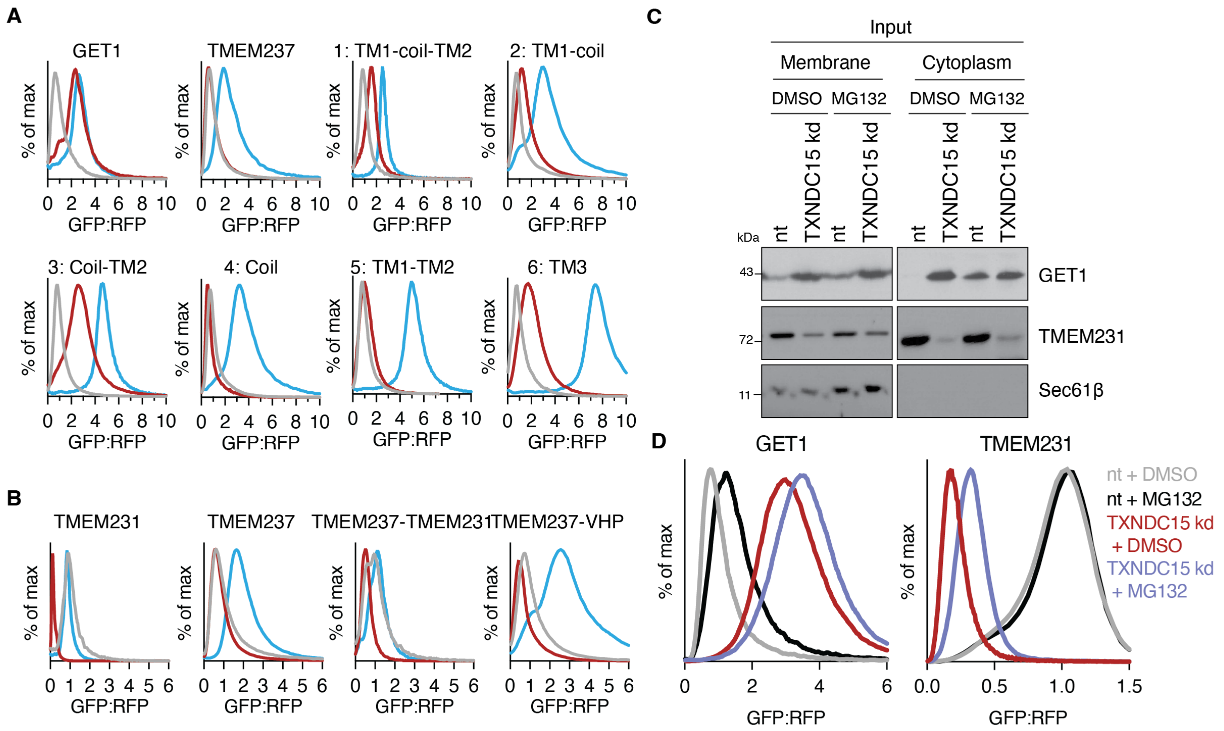


**Figure S5. TXNDC15 regulates the recruitment and interaction of substrates to MARCHF6 based on their structural features, related to Figure 4**

(A) Representative histograms of the fluorescence reporter assay that is quantified in Figure 4A. GET1, TMEM237 and their chimeras were fused to a GFP-2A-RFP reporter cassette and expressed in HEK293T wt, TXNDC15 kd or MARCHF6 kd cells. Histograms were normalized to the median GFP:RFP ratio in wt cells.

(B) Representative histograms of the TMEM231 and TMEM237 chimeras as described in (A) and quantified in Figure 4B.

(C) Input samples of retrotranslocation experiment. As described in Figure 4F, K562 cells carrying the inducible reporter cassettes for either GET1-GFP or TMEM231-GFP were transduced with either a TXNDC15 or non-targeting (nt) control sgRNA and treated with the proteasome inhibitor MG132 or a DMSO control. Cells were divided into cytosol and membrane fractions by ultracentrifugation, and GET1 or TMEM231 were enriched by GFP-IP. Input samples were analyzed by western blot using anti-GFP and anti-SEC61b antibodies. While depletion of TXNDC15 results in stabilization of GET1, TMEM231 levels are decreased compared to wildtype cells. SEC61b is only found in the membrane fraction, not the cytosol, which indicates a clean separation of the two samples.

(D) Fluorescence reporter assay of GET1 (left) or TMEM231 (right) in K562 cells treated with proteasome inhibitor. Samples described in (C) and Figure 4F were analyzed by flow cytometry at the time of harvest after 6 h of proteasome inhibitor treatment. Histograms were normalized based on the median GFP:RFP ratio of the nt + DMSO control. GET1 is stabilized by the inhibitor treatment in nt and TXNDC15 kd while TMEM231 is only stabilized in TXNDC15 kd cells. This potentially points towards a change in the dominant degradation pathway between kd and wt cells.

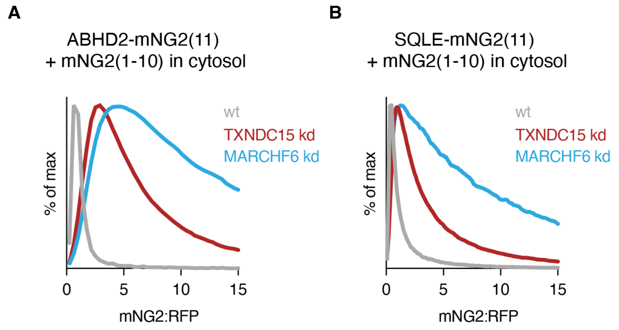


**Figure S6. TXNDC15 regulates membrane protein homeostasis in the ER, thereby regulating ciliogenesis, related to Figure 5**

(A) Two hits identified through our proteomics assays (Figure 5B) were tested using the ratiometric fluorescent reporter to ascertain whether TXNDC15 affected the post-translational stability of these nascent proteins. Using our split mNeon system (described in Figure S1G) ABHD2-mNG2(11) was expressed in HEK293T wt, TXNDC15 kd and MARCHF6 kd cells that were co-transduced with mNG2(1-10) in the cytosol. Histograms were normalized to the median mNG2:RFP ratio of wt cells. Similar to GET1, loss of TXNDC15 impairs degradation of ABHD2.

(B) Fluorescence reporter assay as described in (A) for SQLE, also identified through our proteomics experiment.

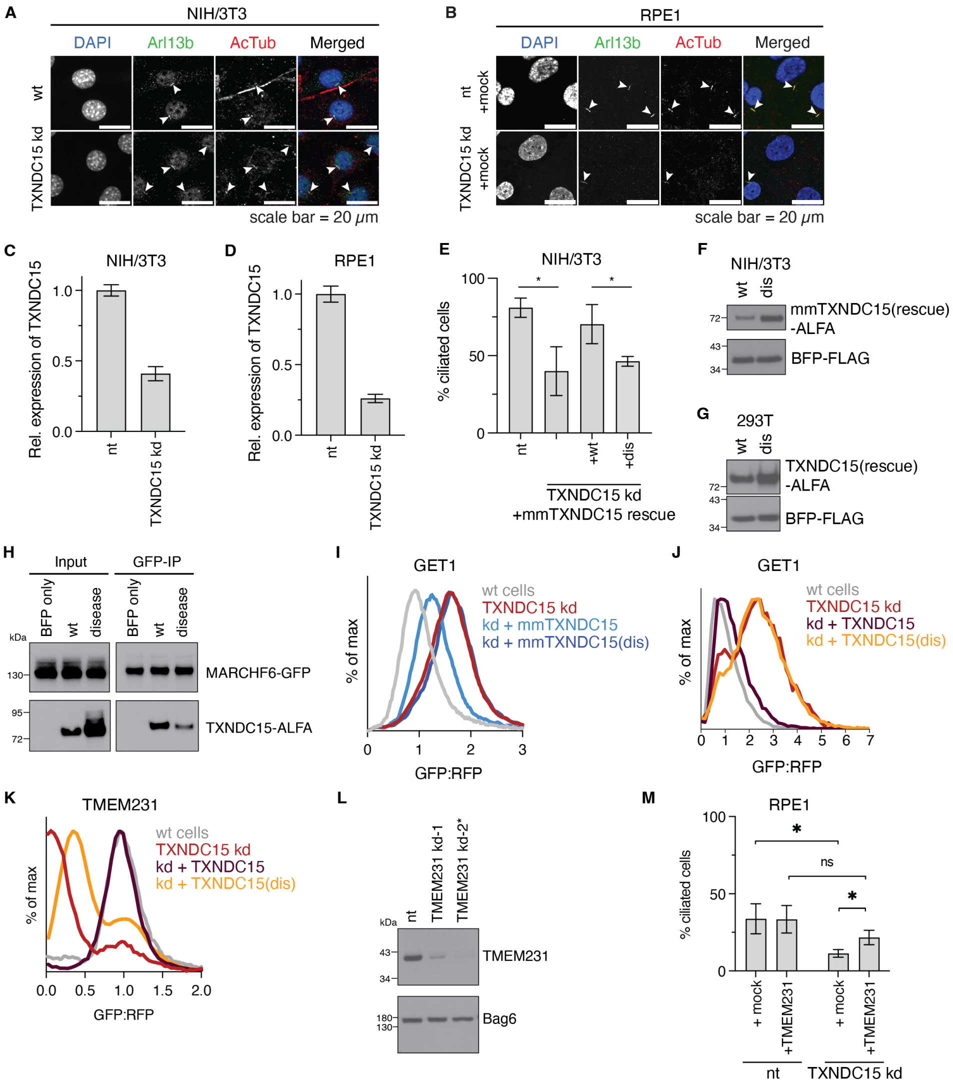


**Figure S7. TXNDC15 regulates membrane protein homeostasis in the ER, thereby regulating ciliogenesis, related to Figure 6**

(A) Comparison of Arl13b and acetylated tubulin stains in NIH/3T3 cells as shown in Figure 6B. Acetylated tubulin is a robust cilia stain that overlaps with Arl13b in wt and TXNDC15 kd cells. The same images are shown as in Figure 6B. All subsequent quantification of cilia was performed using Arl13b because the antibody contained less background, simplifying analysis.

(B) Comparison of Arl13b and acetylated tubulin immunofluorescence in RPE1 cells as shown in Figure 6D. In the absence of Arl13b stained cilia, no cilia were detected using the acetylated tubulin stain. All subsequent quantification of cilia was performed using Arl13b because the antibody contained less background, simplifying analysis.

(C) Characterization of a murine NIH/3T3 TXNDC15 stable CRISPRi knockdown cell line used in Figure 6B,C. NIH/3T3 cells expressing dCas9 were transduced with a guide targeting TXNDC15 along with a puromycin selection marker. Transduced cells were subjected to puromycin selection. Relative expression of TXNDC15 in the NIH/3T3 TXNDC15 stable knockdown (kd) cell line compared to a non-targeting (nt) control was quantified by qPCR. Error bars represent errors as determined by error propagation, calculated from the primer efficiency and measurements taken from 3 technical replicates.

(D) Characterization of RPE1 cells subjected to CRISPRi TXNDC15 knockdown that were used in Figure 6D,E. RPE1 cells expressing dCas9 were transduced with sgRNA targeting TXNDC15 along with a puromycin selection marker. Transduced cells were subjected to puromycin selection. Relative expression of TXNDC15 in RPE1 cells transduced with TXNDC15 sgRNA guide compared to a nt control was quantified by qPCR. Error bars represent errors as determined by error propagation, calculated from the primer efficiency and measurements taken from 3 technical replicates.
(E) Quantification of ciliated NIH/3T3 cells in wt and TXNDC15 kd cell lines. TXNDC15 kd was rescued by overexpression of either murine mmTXNDC15 wt or the mmTXNDC15 ciliopathy-mutant (dis, Δ209-213). Ciliated cells were imaged by immunofluorescence microscopy (see Figure 6B) and the number of cilia and cells per microscope slide were counted. Mean and standard deviation for 3 replicates are shown (wt: 420 cells; TXNDC15 kd: 445 cells; mmTXNDC15 wt rescue: 207 cells; mmTXNDC15(dis): 184 cells). Statistical significance was evaluated using a t-test (($p$(nt/TXNDC15 kd) $< 0.014$; $p$(TXNDC15wt rescue/TXNDC15dis rescue) $= 0.034$).
(F) Determination of expression levels by western blot analysis of overexpressed murine mmTXNDC15 in TXNDC15 kd NIH/3T3 cells as used in (C), (E), (I) and Figure 6B,C. ALFA-tagged mmTXNDC15 was overexpressed from the same mRNA as BFP-FLAG so that the sample was normalized to the number of cells expressing BFP. Western blots were performed using the anti-ALFA-Nb-HRP and anti-FLAG-HRP antibody, respectively.
(G) Western blot analysis of overexpression levels of TXNDC15 wt and the ciliopathy mutant (dis, Δ225-229) in TXNDC15 kd HEK293T cells as used in Figure 6H. Samples were analyzed as described in (F).
(H) MARCHF6-TXNDC15 interaction assay. Expi293 cells expressing MARCHF6(C9A)-GFP were transduced with the TXNDC15-ALFA rescue mutants as indicated. Following a GFP-IP, samples were analyzed by western blot using anti-GFP and anti-ALFA antibodies.
(I) Analysis of the rescue efficiencies of mmTXNDC15 wt and mmTXNDC15(dis) in NIH/3T3 cells. The stability of the GET1-GFP-2A-RFP reporter was analyzed by flow cytometry and curves were normalized to the median GFP:RFP ratio of the nt control. mmTXNDC15 wt overexpression leads to a partial rescue of the TXNDC15 kd phenotype while the ciliopathy mutant does not rescue GET1 degradation. These results are consistent with those in Figure 6H, where the same assay was performed in human HEK293T cells.
(J) Analysis of rescue efficiencies of the TXNDC15 wt and (dis, Δ225-229) ciliopathy mutant in RPE1 cells as described in (I). TXNDC15 wt can fully rescue the TXNDC15 kd phenotype of the GET1 reporter while the ciliopathy mutant cannot.
(K) Analysis of rescue efficiencies of the TXNDC15 wt and (dis, Δ225-229) ciliopathy mutant in RPE1 cells as described in (I) using TMEM231 as reporter. Similarly to GET1, the ciliopathy mutant cannot restore TMEM231 stabilization to wildtype levels.
(L) RPE1 cells were transduced with two different TMEM231 kd sgRNAs or a nt control. Samples were analyzed by western blot and guide 2 (marked with *) was selected for further experiments shown in Figure 6I,J.
(M) Quantification of ciliated RPE1 cells in wt and TXNDC15 kd cells in the presence of absence of TMEM231 overexpression. Ciliated cells were imaged by immunofluorescence microscopy (see Figure 6B) and the number of cilia and cells per microscope slide were counted. Mean and standard deviation for 4 replicates are shown (nt + mock: 343 cells; nt + TMEM231: 540 cells; TXNDC15 kd + mock: 262 cells; TXNDC15 kd + TMEM231: 545 cells). Statistical significance was evaluated using Welch's t-test (($p$(nt + mock/TXNDC15 kd + mock) $= 0.016$;

p(nt + TMEM231/TXNDC15 kd + TMEM231) = 0.072; p(TXNDC15 kd + mock/TXNDC15 kd + TMEM231) = 0.014).
